# Mosquitoes of Macha – new mosquito species records from Macha and surrounding locations in southern Zambia with photographs

**DOI:** 10.64898/2026.01.09.698444

**Authors:** Isabella S. Flores, Valerie T. Nguyen, Jun Soo Bae, Alexander Urlaub, Raymond Gellner, Rebekah C. Kading, Ana L. Romero-Weaver, Limonty Simubali, Kochelani Saili, Edgar Simulundu, Lawrence E. Reeves, Yoosook Lee

**Author notes:** These authors have contributed equally to this work and share senior authorship (Lawrence E. Reeves, Yoosook Lee).

## Abstract

The biodiversity of mosquito taxa beyond *Anopheles* malaria vectors has been understudied in many parts of Africa. Here, we provide a new species record of mosquitoes from Macha and its surrounding locations in southern Zambia, as well as an updated list of species from the region. With the addition of 19 new species records in this region, the total list of mosquito species reported in Macha is now 46. We present high-resolution focus-stacked photographs of some of these species, many of which were only previously available as text or drawing descriptions in mosquito identification books and articles. We also include 34 COI sequences from Zambian mosquito species, five of which had no prior genetic information on Genbank. This visual guide as well as additional COI sequences provide essential tools for accurate mosquito surveillance in southern Zambia and establishes a baseline data for future invasive species monitoring.

## Introduction

Accurate records of mosquito species in a region are important for biodiversity management, disease/pathogen surveillance, and invasive species monitoring and control. Because of their reliance on blood for reproduction, mosquito diversity often reflects the diversity of animals within the ecosystems they inhabit. Therefore, studying the mosquito fauna and its host associations could reveal hidden biodiversity (Reeves et al. 2021).

With efforts to understand diseases of medical and veterinary importance, such as Rift Valley fever and other arboviruses, or avian malaria, more detailed mosquito community data and taxonomy work have been conducted, illuminating the diversity of mosquito fauna in Africa (Reinert et al. 2009, Mutebi et al. 2012, Kaddumukasa et al. 2014, Mayanja et al. 2014, Wilkerson et al. 2015, Muiruri et al. 2015, Cornel et al. 2018, Mutebi et al. 2018, Cornel et al. 2020, Tumusiime et al. 2023). Due to public health importance, however, research on African mosquitoes has been heavily focused on malaria vectors contained within the *Anopheles* genus (Nguyen et al. 2023, Irish et al. 2020). Decades of such funding practice created a knowledge gap for other African mosquitoes, leaving valuable mosquito identification tools for some genera and subgenera untouched for over 80 years (Cornel et al. 2020).

With increasing risks of invasive mosquito introductions and disease threats associated with invasive species globally, the need for accurate documentation on the native ranges of a particular mosquito species has been expressed (Seok et al. 2024). For example, *Anopheles stephensi* was introduced in Djibouti in 2012 and now has spread to six other countries, including Ethiopia, Somalia, Kenya, Sudan, Nigeria, and Ghana (WHO 2023). Unlike native malaria vectors, this invasive species thrives in urban environments, adding a malaria burden in densely populated areas using containers of varying sizes holding water (Yared et al. 2025). Following the introduction of this invasive species, Djibouti alone experienced a surge of malaria cases from 27 in 2012 to 58,916 in 2022 (WHO 2023). In this case, because the introduced species were from different continents, it was relatively straightforward to label this an invasive species. However, given the paucity of information, it would be much harder to determine invasive vs. non-native species in one region of Africa if that species originated from a different African region.

Africa also faces severe headwind in preserving biodiversity in the face of rapid development and urbanization (Seto et al. 2012, Simkin et al. 2022). In 2015, Africa had urban land cover totaling 365,626 km^2^ in the continent, according to the Center for International Earth Science Information Network (World Bank, 2025). And the United Nations Department of Economic and Social Affairs (UN DESA) expects the global urban area will be doubled by 2050 (UN DESA 2018). Africa is also undergoing rapid urbanization, and the need to document the diversity of mosquito fauna in the continent is more urgent than ever if we are to be able to accurately assess the impact of environmental changes on the region’s biodiversity.

The first list and images of mosquitoes from Macha was created in 2006 (Kent 2006). A team of our scientists revisited the site in 2024 and collected mosquito specimens, which were then catalogued using high-resolution stacked images to provide updated records of mosquito species present in Macha, Zambia, and surrounding areas. This pictorial reference will assist with entomological research in Africa by providing an additional tool to identify common culicine species that may be of academic or public/veterinary health interest.as well as in assessing biodiversity

## Material and methods

### Mosquito collection

Mosquitoes were collected in Macha (16° 24’ S, 26° 47’ E) and neighboring towns Sinazongwe (17° 15’ S, 27° 28’ E) and Livingstone (17° 51’ S, 25° 52’ E). A variety of catch methods, including pyrethrum spray catch, CDC light traps, landing catches, aspirations using a Entovate^®^ 200mm aspirator (Entovate^®^, Melbourne, FL; Figure 1A), and larval collections using plastic dippers and transfer pipettes (Figure 1B and 1C) were used to collect mosquito specimens. The larvae were held in Whirl-Pak plastic bags (Whirl-Pak, Austin, TX) for transport between collection sites and the Macha Research Trust Laboratory (Figure 1D). Adults were held in modified RESTCLOUD 12in x 12in x 12in insect cages (China, company info unavailable) with a 4” diameter stockinette sewn on one side of the cage for access to minimize mosquito escapes (Figure 1E). At Macha Research Trust, the larvae were held in plastic Ziploc containers or Whirl-Pak plastic bags (Whirl-Pak, Austin, TX) until they turned into pupae. The pupae are then transferred to a cage or 7-dram polystyrene plastic vial as shown in Figure 1F. Emerged adults were frozen at 0 °C overnight for photography.

**Figure 1.**
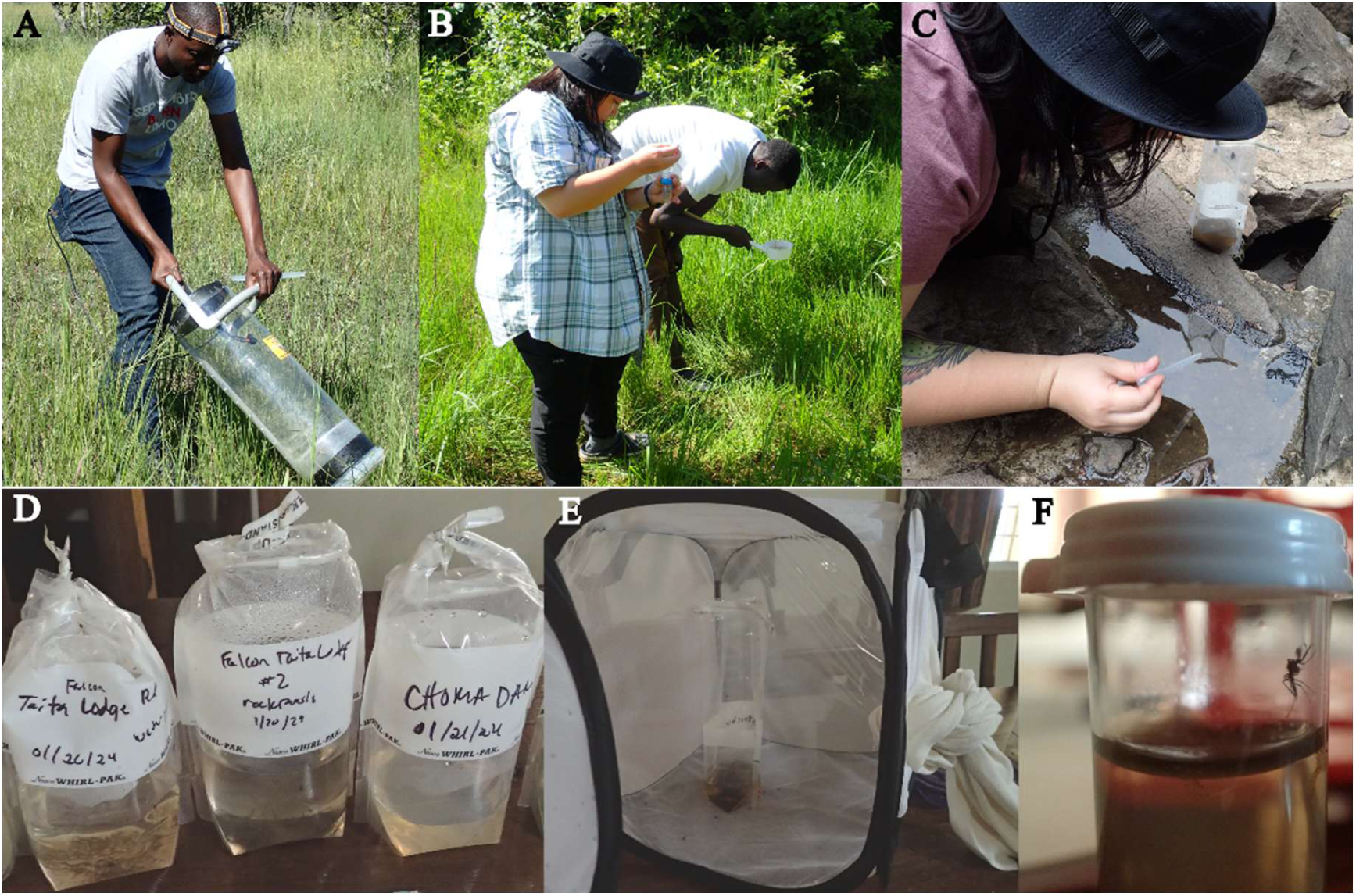
Methods of mosquito collection in Macha, Zambia. A: Entovate^®^ 200mm aspirator operation by co-author, Limonty Simubali. B: larval collection using dipper demonstrated by co-author Valerie T. Nguyen and Macha Research Trust Entomology team staff. C: larval collections directly from a rock pool using transfer pipettes demonstrated by co-author Valerie T. Nugyen. D: Whirl-Pak holding mosquito larvae for transport. E: A modified insect cage with a 4” diameter stockinette sewn on one side of the cage for access to adult mosquitoes, minimizing their escape. F: 7 Dram vial holding a mosquito that emerged from the pupal stage. All images are taken by the authors and original to this manuscript.

### High-resolution imaging

For high-resolution stacked photographs, the collected adult mosquitoes were frozen at 0 °C overnight and mounted on a glass microscope slide within 24 hours. The mounted mosquito was photographed with a Canon 5D SR digital SLR camera using a focus-stacking system that consisted of a 5x or 10x Mitutoyo infinity-corrected microscope objective attached to a Canon 200mm L prime lens. The camera and lens were mounted onto an automated StackShot rail (Cognisys Inc., Traverse City, MI). The camera was moved so that the mosquito specimen was just out of focus, then moved again in steps of either 7μm (5x) or 15μm (10x), with one photograph taken between each step until the mosquito was out of focus in the opposite direction. After taking photographs, a leg of each specimen was preserved in alcohol for DNA barcoding. All photographs were taken at Macha Research Trust in Zambia.

The raw images were digitally merged to create one image with the entire specimen in focus using the Helicon Focus program version 8.2.13 and then edited to clean the background using Adobe Photoshop version 25.5.1.

### Mosquito species identification

Mosquitoes were identified to the species level based on morphology and DNA barcoding. For morphological identification, we used a variety of keys, including Edwards (1941), Gilles and Coetzee (1987), Jupp (1996), Huang and Ward (1981), Service (1990), Huang (2001), Huang (2004), Huang and Rueda (2015), Huang and Rueda (2016), and Coetzee (2020). Genus, subgenus, and species names follow Wilkerson et al. system (2015) for the Aedini tribe and Mosquito Taxonomy Inventory (Harbach 2013) for the others.

For DNA barcoding, a region of the cytochrome c oxidase subunit I (COI) was amplified by polymerase chain reaction (PCR) using the primer pair LCO1490 (5’ GGT CAA CAA ATC ATA AAG ATA TTG G 3’) and HCO2198 (5’ TAA ACT TCA GGG TGA CCA AAA AAT CA 3’; Hebert et al. 2004). We used the PCR reaction protocol described in Reeves et al. (2021).

The DNA sequences were generated using commercial Sanger sequencing services. The resulting sequences were trimmed to remove low-quality bases before searching for matches in online databases like the Barcode of Life (BOL) (Ratnasingham and Hebert, 2007) or NCBI Nucleotide database using BLASTN (NCBI 1988). Species with 98% COI sequence similarity or higher were used to determine the species of a sample. The phylogenetic tree based on COI sequences was generated using the Maximum Likelihood method with 500 bootstrap MEGA12 version 12.0.14 (Kumar et al. 2024). For brevity, genus names in the tree figure are abbreviated as suggested by Reinert (2001).

### Occurrence Record Analysis

We used the occurrence records in the Global Biodiversity Information Facility (GBIF) Backbone Taxonomy (GBIF Secretariat, 2023) to note whether our observation is the first record in Zambia or a geographic region. GBIF aggregates data from multiple sources such as Genbank, Barcode of Life (BOL), research-grade iNaturalist data, VectorBase, and other museum or research institute collections. In addition, we used Edwards (1941), which noted the distribution of a species. We used the geographic regional groups according to the UN M49 Standard country and area codes for statistical use (UNSD, 1999). The UN M49 Standard classifies Africa into 4 subgroups, namely Eastern, Middle, Southern, and Western Africa. For example, Zambia is grouped as the Eastern African country by the M49 standard.

## Results

A total of 46 mosquito species have been documented in Macha, Zambia as well as in neighboring southern Zambia (Livingstone and Sinazongwe; Table 1). This includes 19 new species records we added to the list from Kent (2006) and we provided high resolution photographs for 43 species. We included a short description of the morphology of each species and used female mosquitoes for photographs to remain consistent with Reeves’ previous photograph collections. While pristine conditions are ideal for consistency, we had limited samples available for some species. Therefore, some mosquitoes appear with bloated abdomens because they recently fed on sugar (e.g. *Aedes hirsutus* and *Aedes circumluteolus*) or blood (e.g. *Aedes embuensis*) at the time of our capture.

**TABLE 1.**
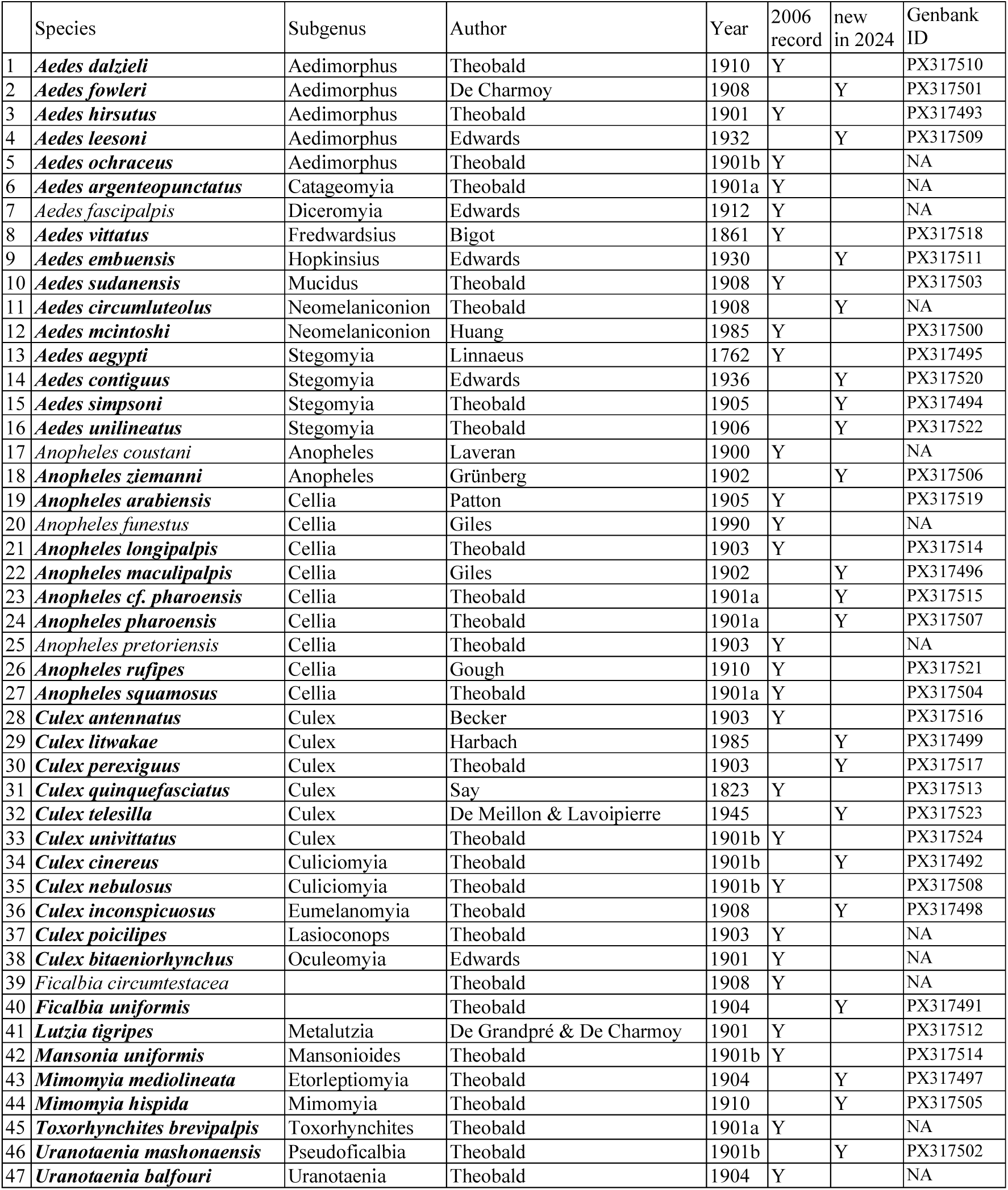
List of mosquito species reported in Macha, Zambia. Genus, subgenus, and species names follow Wilkerson et al. system (2015) for Aedini tribe and Mosquito Taxonomy Inventory (Harbach 2013) for the others. Species with high resolution images are marked in bold text. Genbank ID for COI sequence data is provided when available. Some species failed to amplify COI fragments are noted as NA (=not available).

### *Aedes (Aedimorphus) dalzieli* Theobald 1910 (Diptera: Culicidae)

*Aedes dalzieli* is a dark mosquito with a large portion of the sterna mostly pale (Figure 2A). No pale spots are present in the scutum (Figure 2B). Patches of pale scales are present above the wing base and prescutellar area (Figure 2B). The hind femora as well as tarsi are dark (Figure 2C). No discernible features are mentioned for head, palpi, or proboscis for this species in other literature (Figure 2D). This species has been primarily recorded in Western Africa by the Walter Reed Biosystematics Unit (WRBU, 2018) and the Research Institute for Development (IRD; Ibrahim and Granouillac, 2012), with the exception of Madagascar (Ibrahim and Granouillac, 2012). Edwards (1941) reports that this species occurs in other Eastern African countries like Mozambique, Sudan, and South Rhodesia (now known as Zimbabwe). Therefore, Kent (2006) marks the first record of this species in Zambia.

**FIGURE 2.**
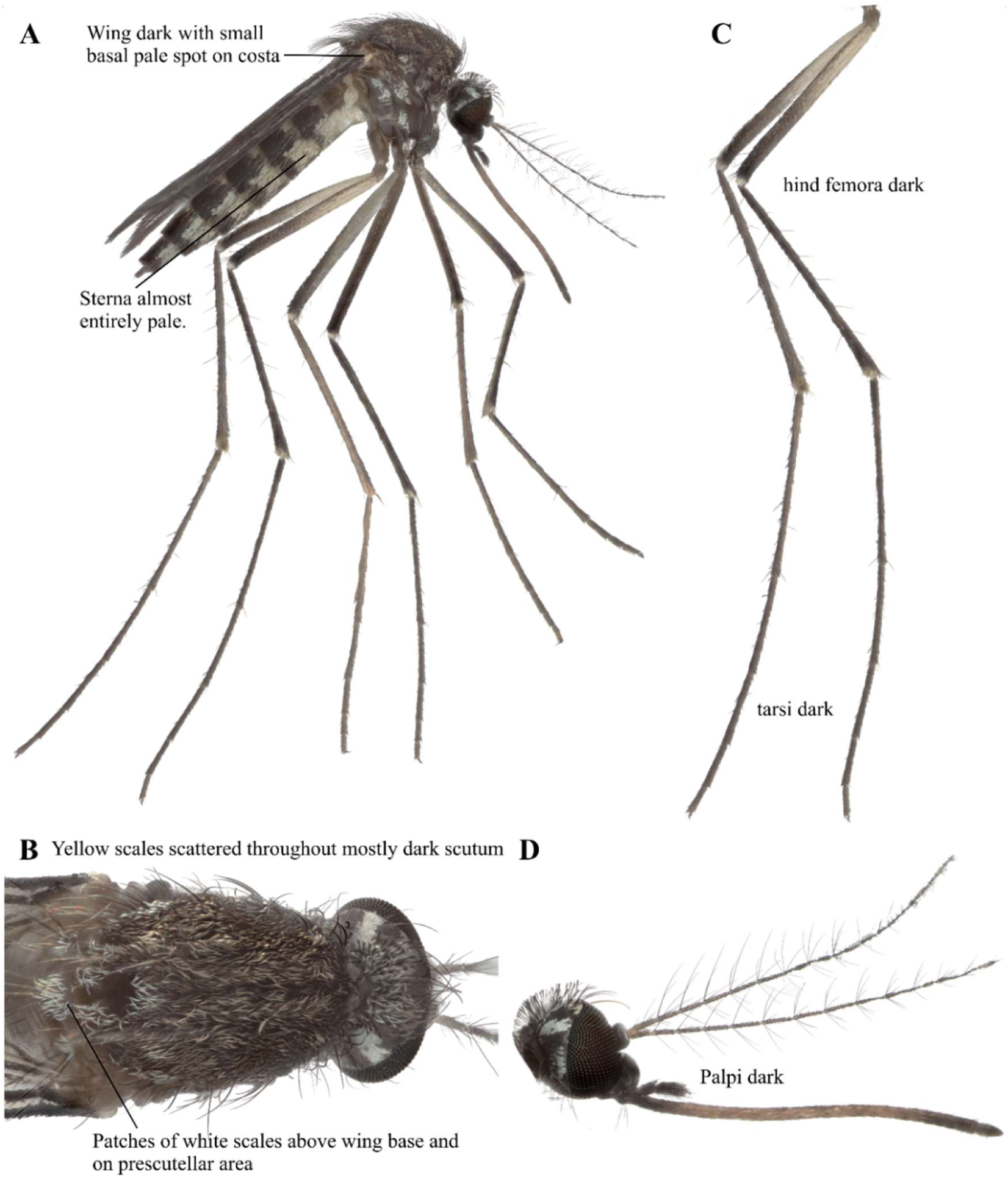
*Ae. dalzieli* focus-stacked images. A: Lateral view. B: Scutum. C: Legs. D: Head.

### *Aedes (Aedimorphus) fowleri* De Charmoy 1908 (Diptera: Culicidae)

*Aedes fowleri* has a pleuron with broad white scales (Figure 3A). The terga have basolateral and median patches (Figure 3A). Scutum does not have obvious features for species identification (Figure 3B). Tarsomeres 1-5 of all legs have broad white basal bands (Figure 3C). Palpi have a median white band (Figure 3D). The proboscis is slightly speckled in the center (Figure 3D). While this species has been observed in Eastern, Southern, and Western Africa (GBIF Secretariat, 2023; Edwards 1941), our 2024 observation marks the first record of occurrence in Zambia.

**FIGURE 3.**
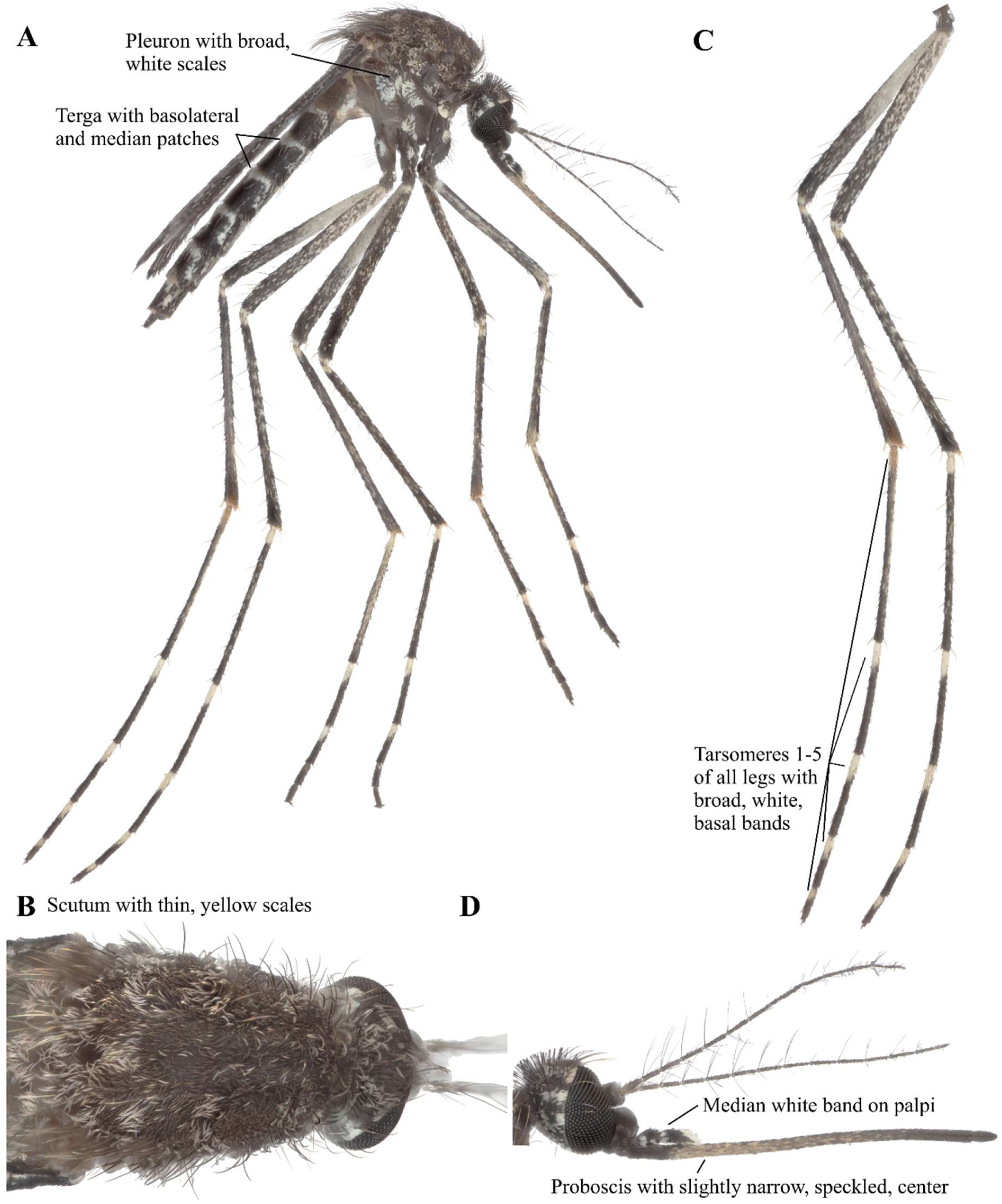
*Ae. fowleri* focus-stacked images. A: Lateral view. B: Scutum. C: Legs. D: Head.

### *Aedes (Aedimorphus) hirsutus* Theobald 1901 (Diptera: Culicidae)

*Aedes hirsutus* is a speckled mosquito with terga 2-6 having basal white bands (Figure 4A). Scutum is mostly dark toward the eyes with narrow yellow scales near the scutellum. Two yellow spots are present in the scutum toward the eyes (Figure 4B). The femora, tibia, and tarsomere 1 are all speckled (Figure 4C). The proboscis has a wide pale region in the center with a black tip (Figure 4D). This species has been previously documented in Macha, Zambia (Kent 2006). While this species has been previously documented in Eastern, Southern, and Western Africa (GBIF Secretariat, 2023; Edwards 1941), Kent’s 2006 observation (Kent 2006) marks the first record of occurrence in Zambia.

**FIGURE 4.**
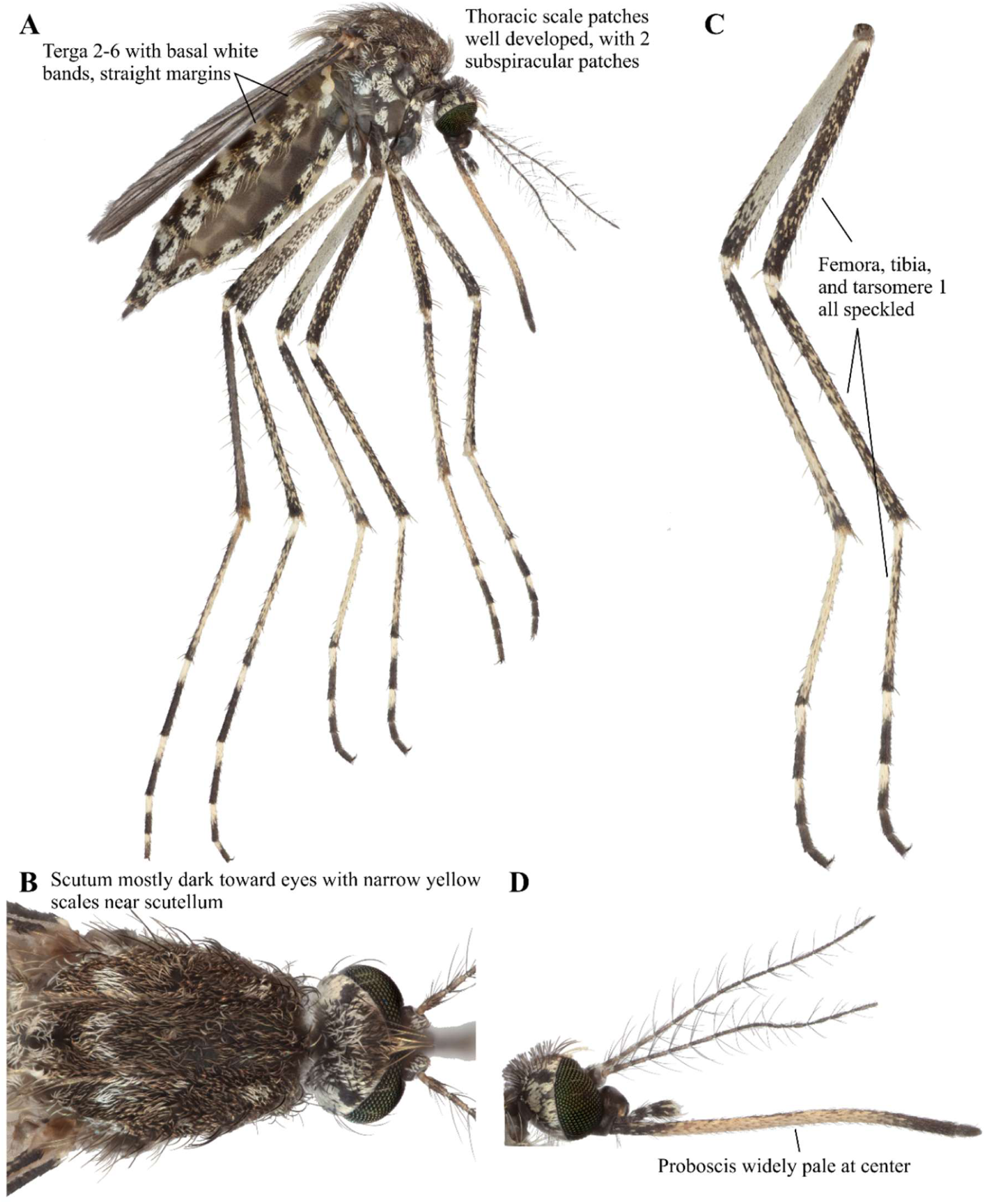
*Ae. hirsutus* focus-stacked images. A: Lateral view. B: Scutum. C: Legs. D: Head.

### *Aedes (Aedimorphus) leesoni* Edwards 1932 (Diptera: Culicidae)

*Aedes leesoni* is a brown mosquito with a banded abdomen (Figure 5A). Scutum has roughly an equal amounts of dark and creamy scales and scutellar scales are pale yellow or creamy (Figure 5B). The mid and hind femora have no spots, and the tarsi are dark (Figure 5C). Palpi and proboscis are dark (Figure 5D). There is only one other occurrence recorded by IRD (Ibrahim and Granouillac, 2012) with an unknown location at GBIF (GBIF Secretariat, 2023). Edwards (1941) reports that this species occurs in Zimbabwe and South Africa. Therefore, this is the first record of the occurrence of this species in Zambia.

**FIGURE 5.**
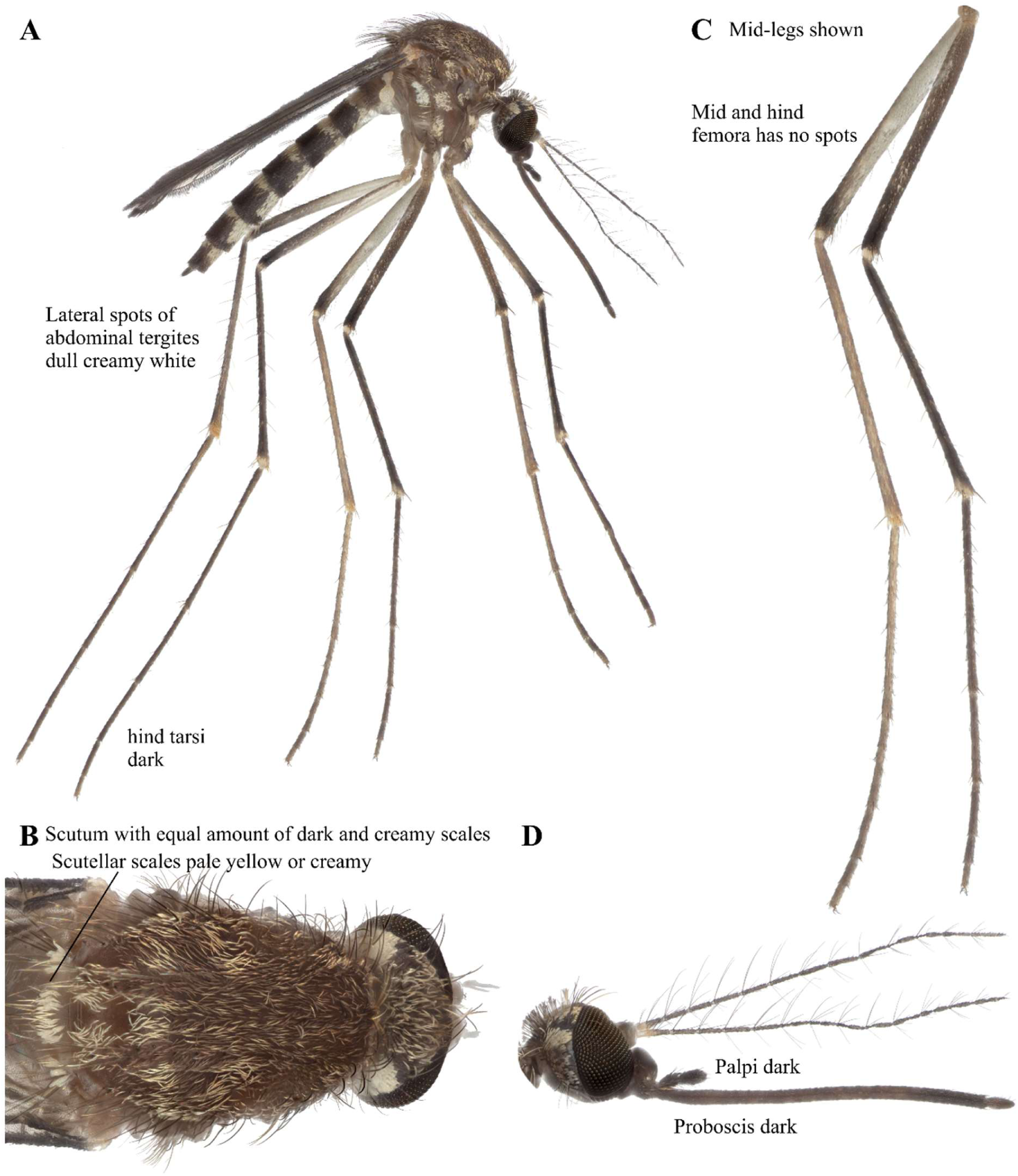
*Ae. leesoni* focus-stacked images. A: Lateral view. B: Scutum. C: Legs. D: Head.

### *Aedes (Aedimorphus) ochraceus* Theobald 1901 (Diptera: Culicidae)

*Aedes ochraceus* is a large yellow and brown mosquito (Figure 6A). Scutum is mostly dark toward the eyes with narrow yellow scales near the scutellum. Two yellow spots are present on the scutum toward the eyes (Figure 6B). The femora, tibia, and tarsomere 1 are all speckled (Figure 6C). Tarsi are largely yellowish. Palpi are brownish and about 1/6 the length of the proboscis (Figure 6D). Proboscis is yellowish with base and tip black (Figure 6D). This species has been previously documented in Macha, Zambia (Kent 2006). While this species has been observed in Eastern, Southern, and Western Africa (GBIF Secretariat, 2023; Edwards 1941), the Kent 2006 record marks the first occurrence of this species in Zambia.

**FIGURE 6.**
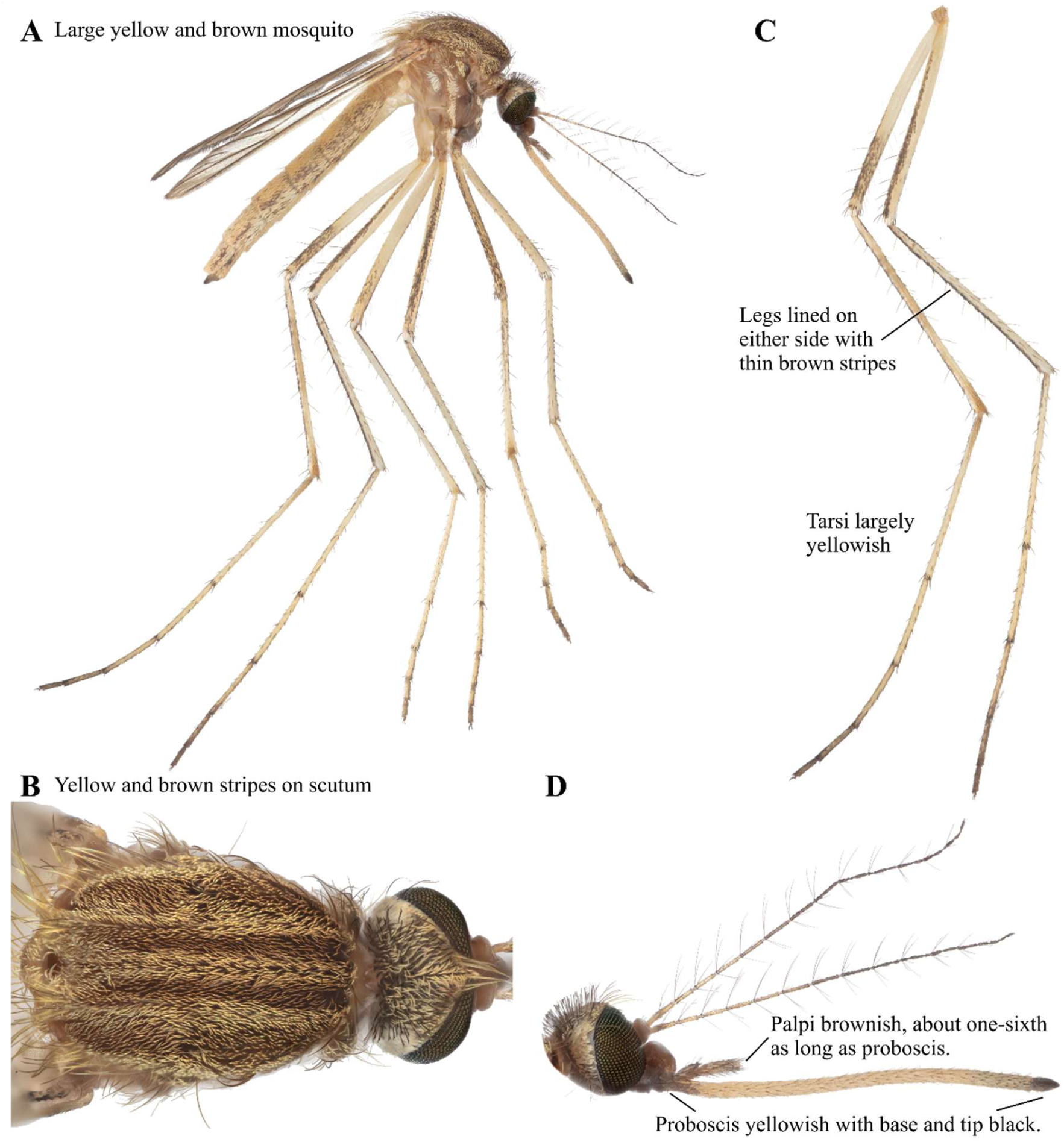
*Ae. ochraceus* focus-stacked images. A: Lateral view. B: Scutum. C: Legs. D: Head.

### *Aedes* (*Catagomyia) argenteopunctatus* Theobald 1901 (Diptera: Culicidae)

*Aedes argenteopuctatus* is a dark mosquito with broad, silvery white scale patches (Figure 7A). Four small white spots of similar size are present on the scutum (Figure 7B). The mid and hind femora have small white spots about 3/4 down the length of the femora (Figure 7C). The palpi are dark and about 1/8 the length of the proboscis (Figure 7D), which is dark brown (Figure 7D). This species has been previously documented in Macha, Zambia (Kent 2006). While this species has been observed in all four regions of sub-Saharan Africa (GBIF Secretariat, 2023; Edwards 1941), the Kent 2006 record marks the first occurrence of this species in Zambia.

**FIGURE 7.**
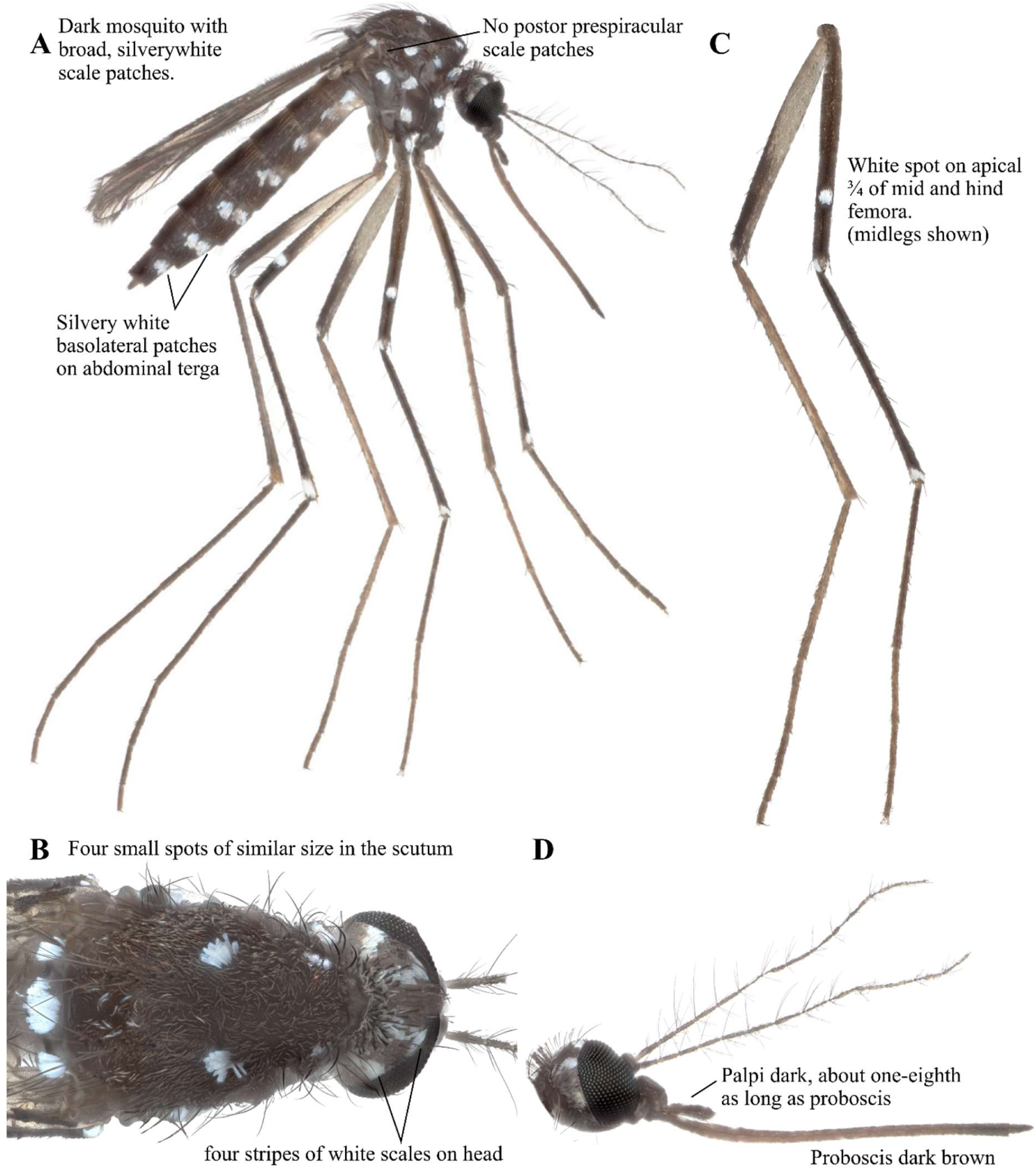
*Ae. argenteopunctatus* focus-stacked images. A: Lateral view. B: Scutum. C: Legs. D: Head.

### *Aedes (Diceromyia) fascipalpis* Edwards 1912 (Diptera: Culicidae)

*Aedes fascipalpis* is a black mosquito with some pale scale markings (Kent 2006). Scutum has creamy white scales with patches of dark brown (Edwards 1941). Their legs are mostly black with small white knee spots (Kent 2006, Edwards 1941). The first three segments of the anterior tarsi and all segments of the hind tarsi have white basal rings (Kent 2006, Edwards 1941). Proboscis is black and can have some scattered pale scales in the middle (Edwards 1941). Palpi are dark and about 1/3 the length of the proboscis (Edwards 1941). This species has been previously documented in Macha, Zambia (Kent 2006), but was not detected in the January 2024 collection trip, and thus high-resolution images are not currently available for this species. The only record of occurrence of this species in GBIF is from South Africa, recorded by the European Bioinformatics Institute (EMBL-EBI, 2025). Edwards (1941) reports that this species occurs in Tanzania, Malawi, Zimbabwe, and South Africa. Therefore, the Kent 2006 record marks the first occurrence of this species in Zambia.

### *Aedes (Fredwardsius) vittatus* Bigot 1861 (Diptera: Culicidae)

*Aedes vittatus* is a black mosquito with silvery white markings (Figure 8A). Scutum has three pairs of small white spots (Figure 8B). The femora are all speckled and have subapical white bands (Figure 8C). The tarsi are banded. Palpi have white tips and some white scales in the middle (Figure 8D). Proboscis is dark with pale scales in the middle (Figure 8D). This species has been previously documented in Macha, Zambia (Kent 2006). While this species has been observed in many countries in Eastern Africa (GBIF Secretariat, 2023; Edwards 1941), the Kent 2006 record marks the first occurrence of this species in Zambia. This species has been spotted in non-native locations like Spain in Europe (iNaturalist contributors, 2025), Caribbean nations like the Dominican Republic (Alarcón-Elbal et al. 2020), Cuba (Pagac et al. 2021, Díaz-Martínez et al. 2021), and Jamaica (Noble et al. 2025) since 2019. Therefore, this species may be of importance in monitoring non-native species in Europe and the Americas.

**FIGURE 8.**
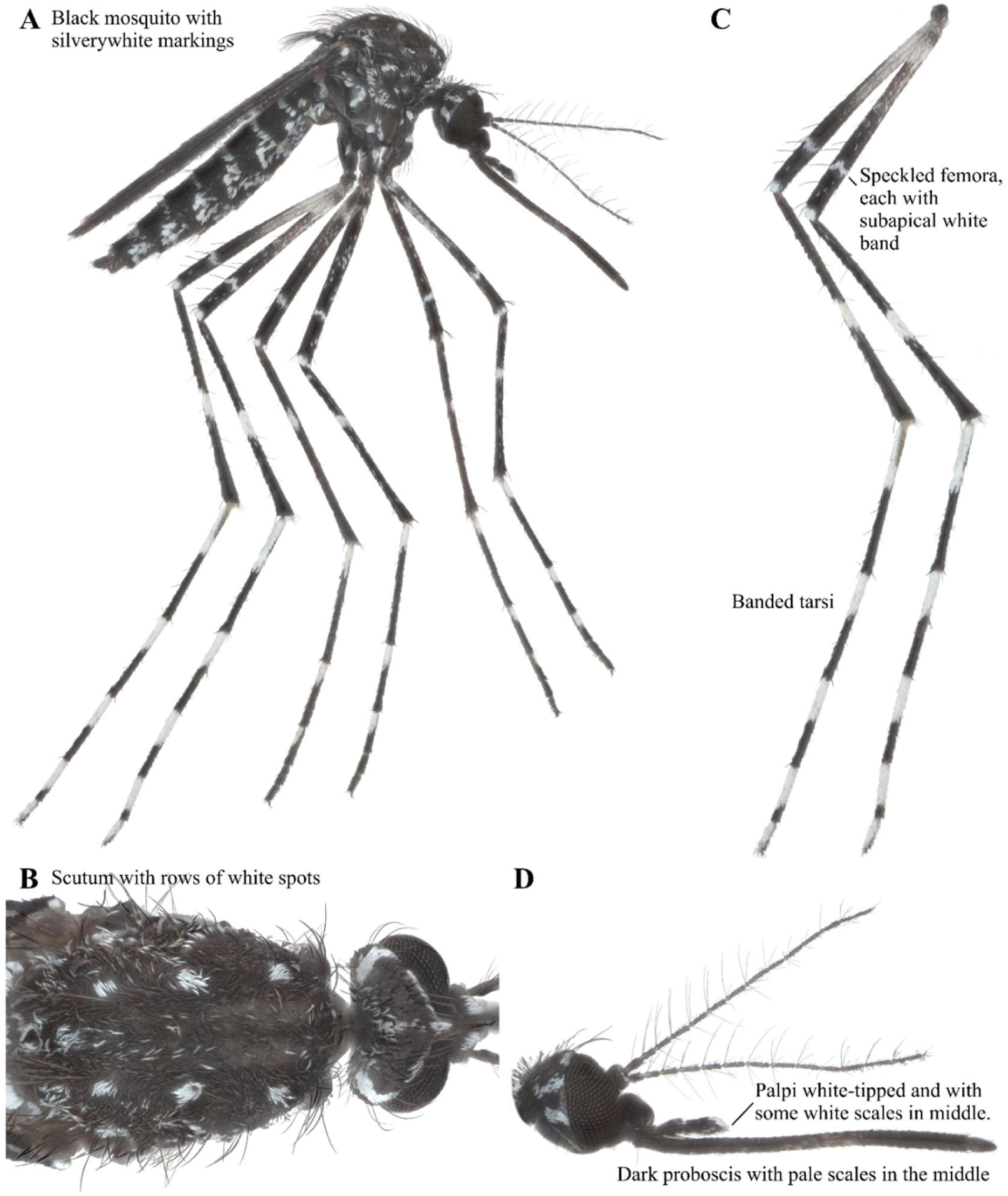
*Ae. vittatus* focus-stacked images. A: Lateral view. B: Scutum. C: Legs. D: Head.

### *Aedes (Hopkinsius) embuensis* Edwards 1930 (Diptera: Culicidae)

*Aedes embuensis* has a patch of creamy-white scales at the base of the costa (Figure 9A). It has scales on the subspiracular area. While the scutal scales of our sample have rubbed off, narrow golden scales on the midlobe of its scutellum are a diagnostic feature to identify this species (Figure 9B; Huang and Rueda, 2017). Hind femora have a subapical patch of pale scales, and the hind tibiae are all dark (Figure 9C). The palpi and proboscis are dark (Figure 9D). This specimen was collected in Sinazongwe on the road to Lake Kaliba in southern Zambia, 120km South-southeast of Macha. This is a new record of occurrence in Zambia (GBIF Secretariat, 2023). The only record of occurrence in GBIF is in Ethiopia, collected by IRD (Ibrahim and Granouillac, 2012), and Edwards (1941) noted that this species occurs in Kenya.

**FIGURE 9.**
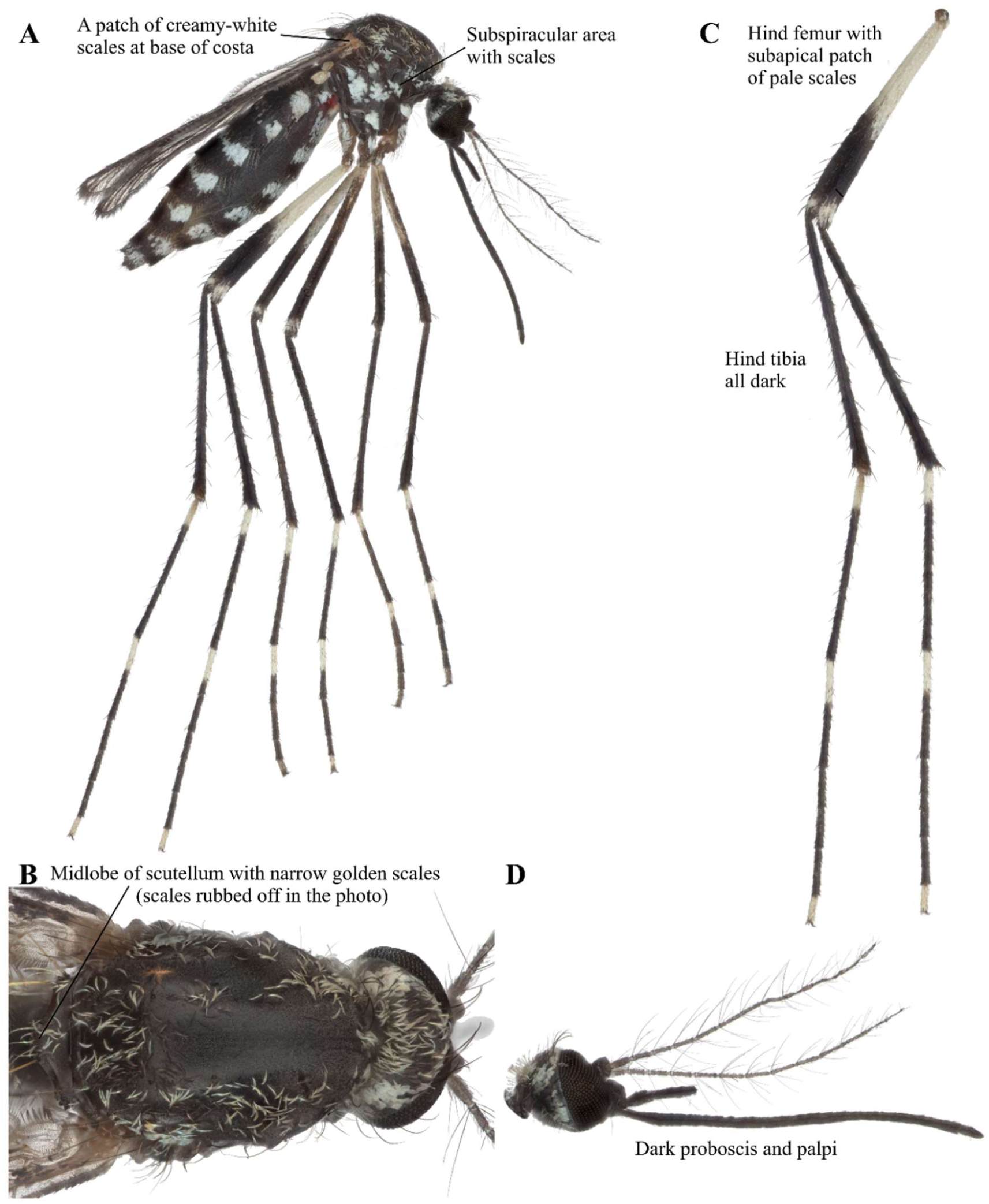
*Ae. embuensis* focus-stacked images. A: Lateral view. B: Scutum. C: Legs. D: Head.

### *Aedes (Mucidus) sudanensis* Theobald 1908 (Diptera: Culicidae)

*Aedes sudanensis* is a large mosquito with broad and erect scales covering its body and legs (Figure 10A). It has relatively long palpi for *Aedes* (Figure 10A). The scutum has mainly white scales with yellow (Figure 10B). Tarsomere 1 of all the legs have a median white band (Figure 10C). The wings have several veins with broad black plume scales (Figure 10E). While this species has been observed in other Eastern African countries like South Africa, Namibia, Malawi, and one occurrence in Senegal (GBIF Secretariat, 2023), the Kent (2006) record marks the first occurrence of this species in Zambia.

**FIGURE 10.**
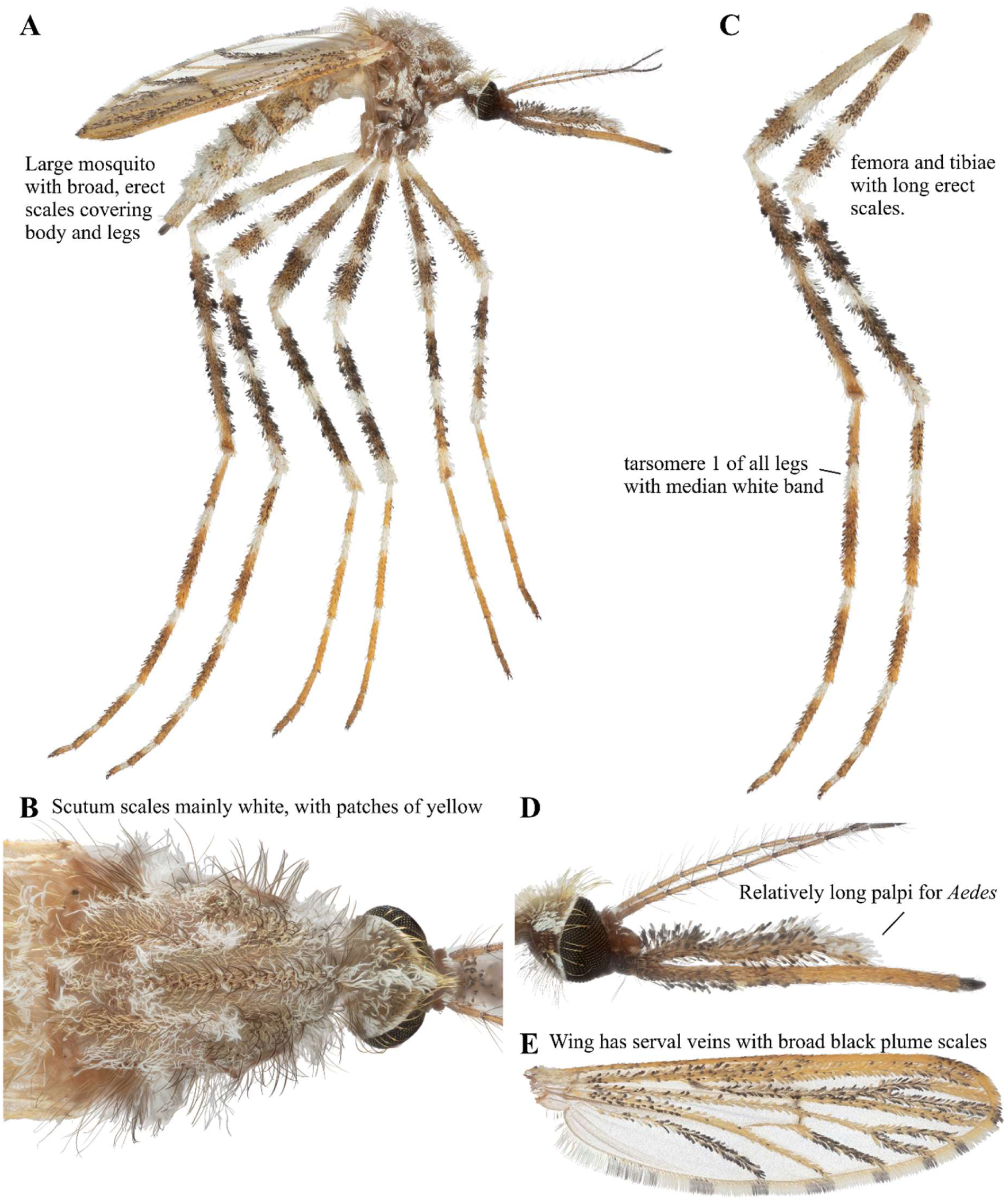
*Ae. sudanensis* focus-stacked images. A: Lateral view. B: Scutum. C: Legs. D: Head. E: Wing.

### *Aedes (Neomelaniconion) circumluteolus* Theobald 1908 (Diptera: Culicidae)

*Aedes circumluteolus* is a yellow-scaled, dark-bodied mosquito (Figure 11A). Its abdomen is banded with the sterna almost entirely yellow-scaled (Figure 11A). Margin of scutum broadly yellow-scaled (Figure 11B). Hind femora are yellowish on the basal half of the anterior (Figure 11C). Proboscis is entirely dark (Figure 11D). While this species has been observed in all four regions of sub-Saharan Africa (GBIF Secretariat, 2023; Edwards 1941), this is the first record of occurrence in Zambia.

**FIGURE 11.**
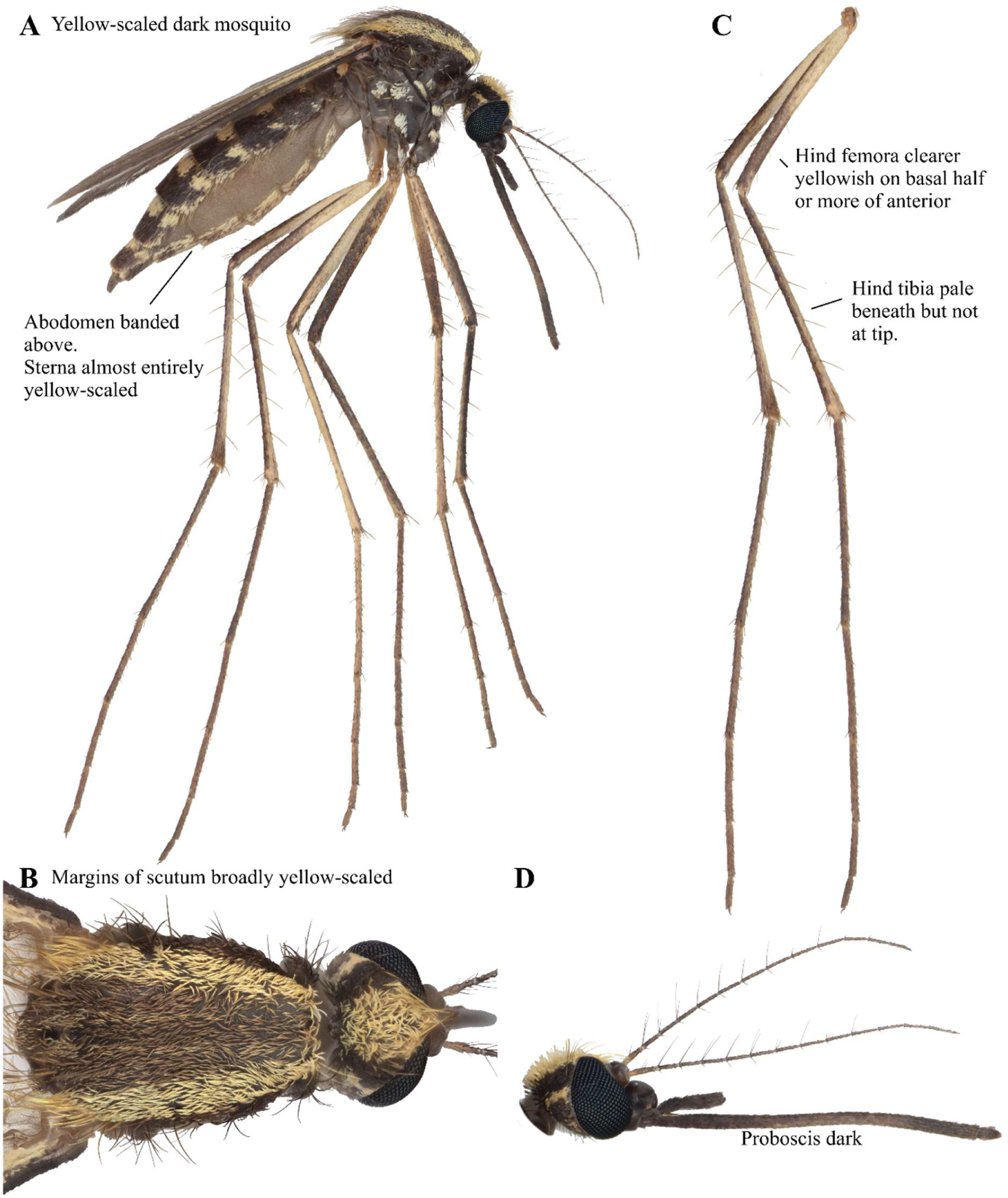
*Ae. circumluteolus* focus-stacked images. A: Lateral view. B: Scutum. C: Legs. D: Head.

### *Aedes (Neomelaniconion) mcintoshi* Huang 1985 (Diptera: Culicidae)

*Aedes mcintoshi* is a dark mosquito with bright yellow scales (Figure 12A). Half or more of the scales are dark in the sterna and the pleuron has subspiracular yellow and narrow scales (Figure 12A). It has lateral bright yellow bands on the scutum (Figure 12B). Basal half or more of the hind femora are pale, and hind tibia and tarsi are dark (Figure 12C). Proboscis is entirely dark (Figure 12D). This species has been previously documented in Macha, Zambia (Kent 2006). While this species has been observed in Eastern, Southern, and Western Africa (GBIF Secretariat, 2023), the Kent (2006) record marks the first record of this species in Zambia. Huang (1985) noted that this species is predominantly in Eastern and Southern Africa, and the records from other African regions may need to be reexamined due to its similarity with other mosquitoes, such as *Ae. circumluteolus* (Figure 11).

**FIGURE 12.**
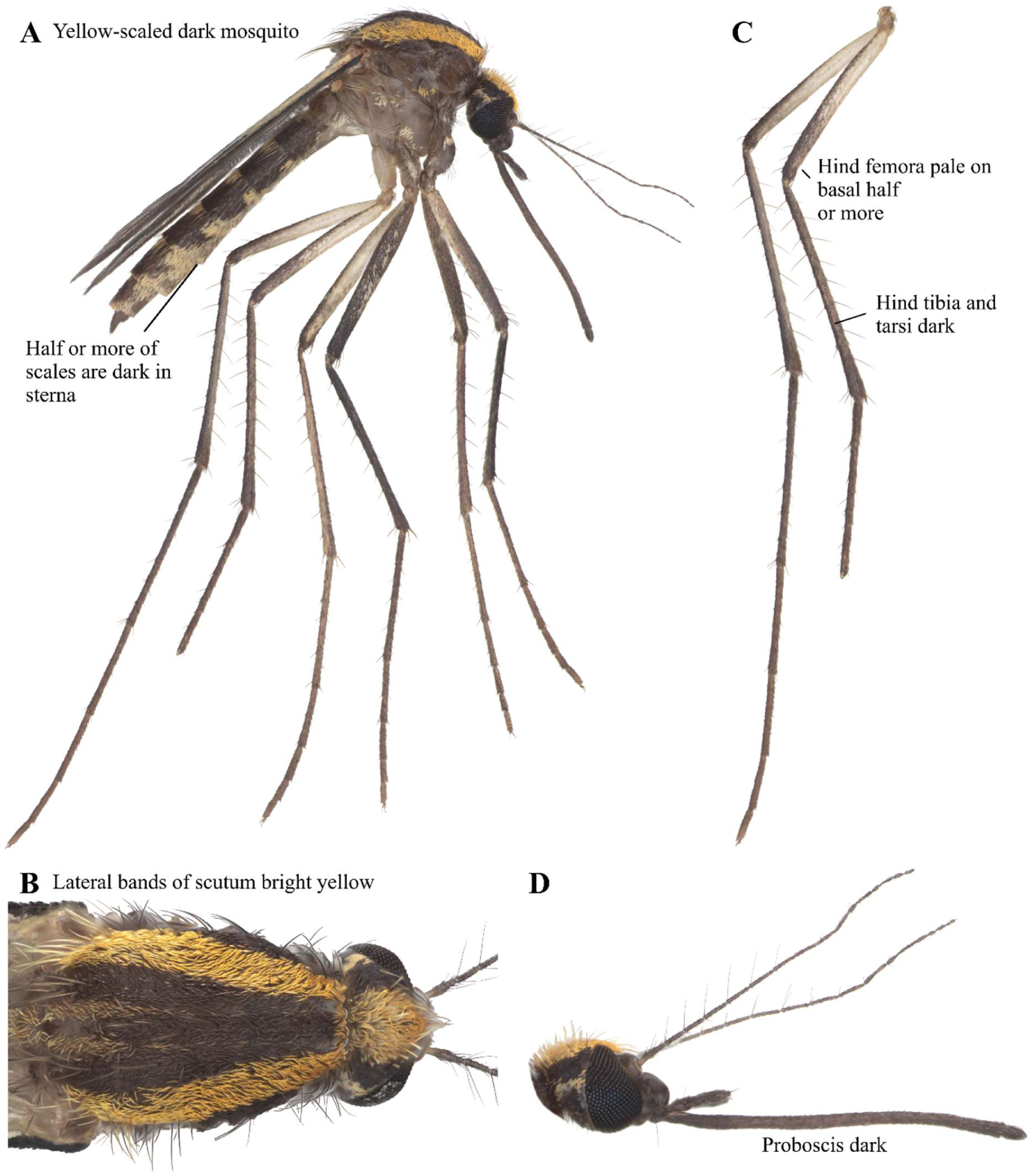
*Ae. mcintoshi* focus-stacked images. A: Lateral view. B: Scutum. C: Legs. D: Head.

### *Aedes (Stegomyia*) *aegypti* Linnaeus 1762 (Diptera: Culicidae)

*Aedes aegypti* is a black mosquito with silver markings (Figure 13A). Middle femora have a white stripe in front from base nearly to tip (Figure 13A). The scutum has a characteristic lyre shaped scale pattern formed with white scales (Figure 13B). The tibia is dark (Figure 13C). Proboscis is all dark (Figure 13D). Two forms of this species include *Ae. aegypti aegypti* and *Ae. aegypti formosus*. *Aedes aegypti formosus* is associated with forested, sylvatic environments and can be distinguished by the lack of a white medial scale patch on the first abdominal tergite that is present in *Ae. aegypti aegypti* (Huang 2004). *Aedes aegypti aegypti* is highly invasive and has been observed in all continents except Antarctica (GBIF Secretariat, 2023). The earliest records of this species are from human observation in 2011 (iNaturalist contributors, 2025), but the Kent (2006) record marks the first occurrence of this species in Zambia known to date.

**FIGURE 13.**
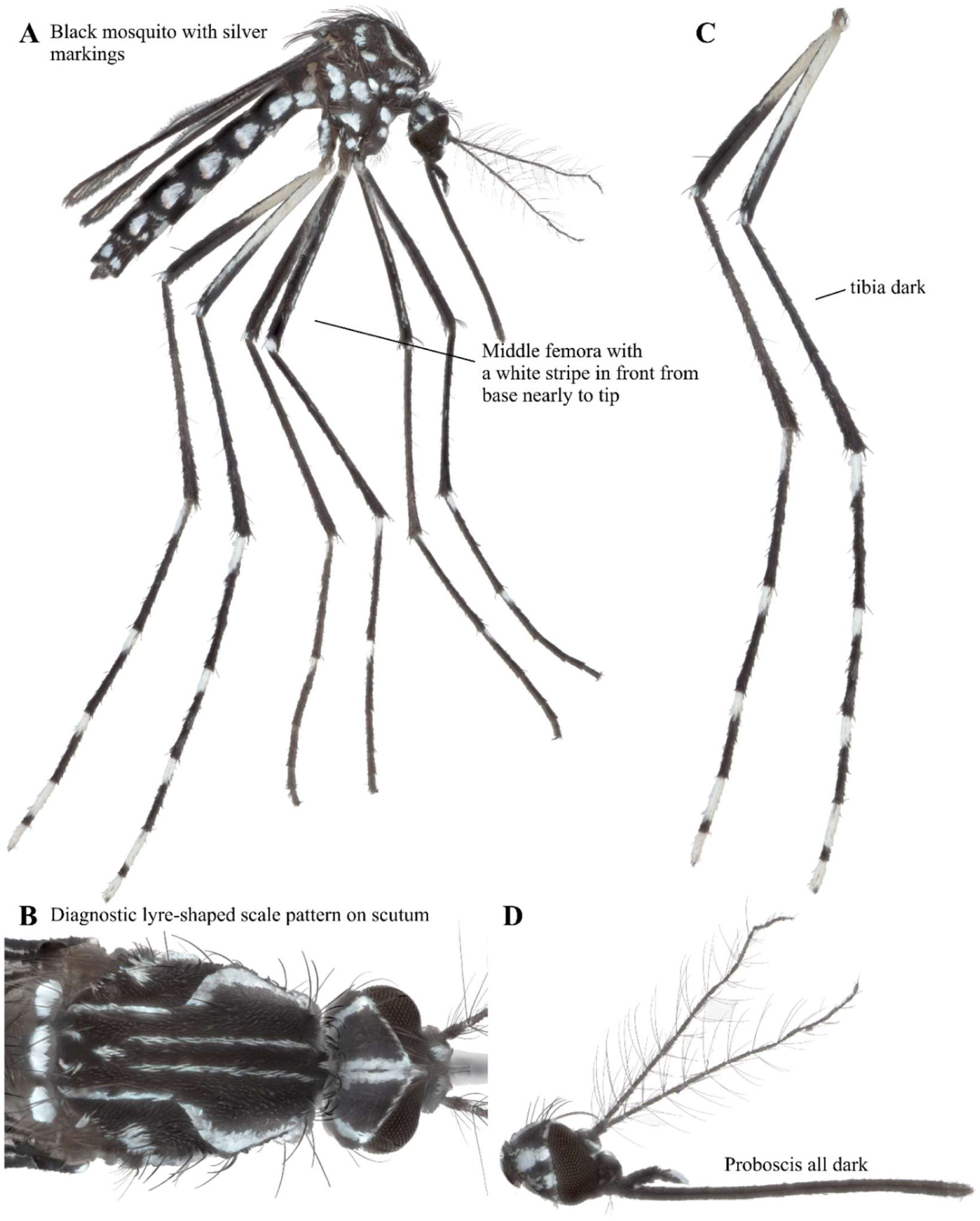
*Ae. aegypti* focus-stacked images. A: Lateral view. B: Scutum. C: Legs. D: Head.

### *Aedes (Stegomyia) contiguus* Edwards 1936 (Diptera: Culicidae)

*Aedes contiguus* has small white spots at the tips of the middle and hind femora (Figure 14A). The middle tibiae are all dark (Figure 14C). Scutum has a pair of large oval white markings (Figure 14B). Proboscis is all black (Figure 14D). The past identification guides (Edwards 1941; Jupp 1996) had conflicting information on the morphological features of this species and this particular specimen could potentially be classified as *Ae. ledgeri.* Moreover, our COI sequence matches with *Ae. contiguus* but based on only one sample. We cannot rule out the current COI sequence labeled as *Ae. contiguus* could have been misidentified. *Aedes contiguus* has been observed in the broader region of Eastern (Ethiopia) and Southern (Zimbabwe, South Africa) Africa as well as Middle Africa (Cameroon) (GBIF Secretariat, 2023; Edwards 1941). GBIF records on *Ae. ledgeri* are limited to Southern Africa (Botswana, South Africa) but Huang (1981) reported much broader distribution of *Ae. ledgeri* in Eastern and Southern Africa including Livingstone, Zambia. Further work is needed to confirm this species call. If this species is indeed *Ae. contiguus,* this marks the first record of occurrence in Zambia.

**FIGURE 14.**
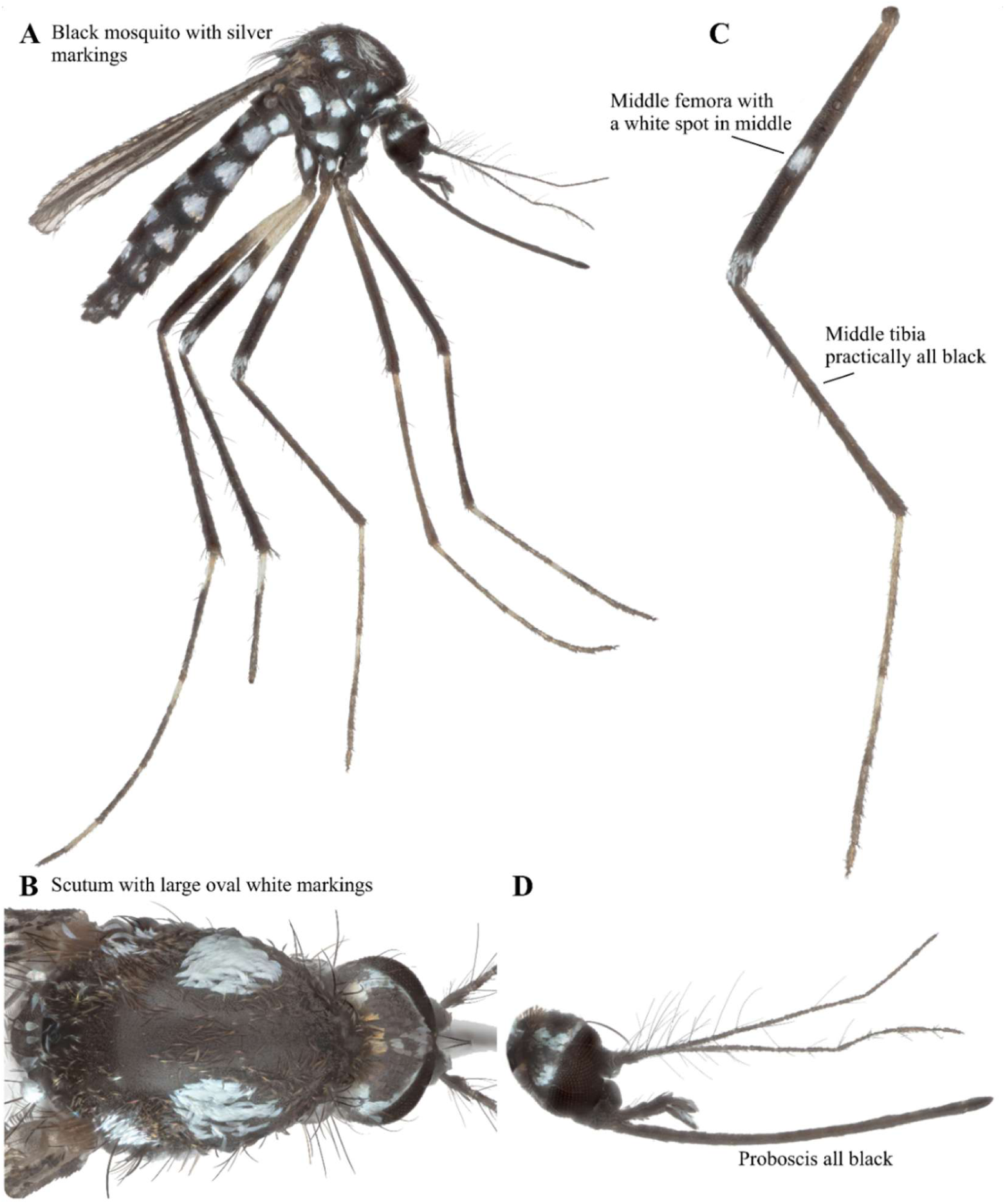
*Ae. contiguus* focus-stacked images. A: Lateral view. B: Scutum. C: Legs. D: Head.

### *Aedes (Stegomyia) simpsoni* Theobald 1905 (Diptera: Culicidae)

*Aedes simpsoni* can look similar to *Ae. contiguus* but the markings differ on scutum and femora (Figure 15A). It has broad white scales on each side of the scutum (Figure 15B). Lateral lobes of scutellum are white-scaled. This mosquito has a white spot on the mid and hind femora (Figure 15C). The hind tibia and tarsomere 4 are all black (Figure 15C). Palpi and Proboscis do not have diagnostic features, but are entirely dark (Figure 15D). While this species has been observed in all four regions of sub-Saharan Africa (GBIF Secretariat, 2023; Edwards 1941), this is the first record of occurrence in Zambia.

**FIGURE 15.**
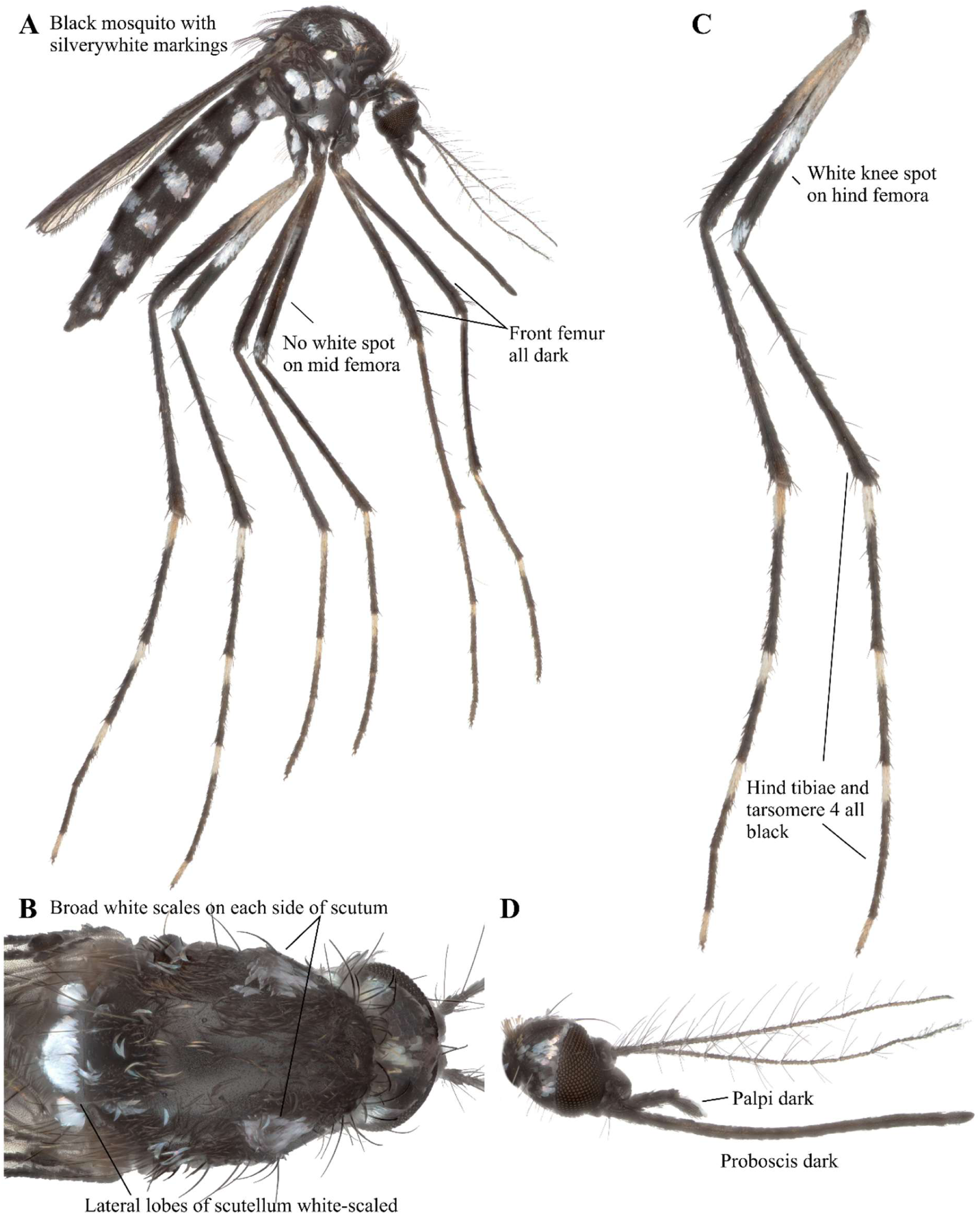
*Ae. simpsoni* focus-stacked images. A: Lateral view. B: Scutum. C: Legs. D: Head.

### *Aedes (Stegomyia) unilineatus* Theobald 1906 (Diptera: Culicidae)

*Aedes unilineatus* has an abdomen with white basal bands and lateral spots on tergites (Figure 16A). Scutum has a bold median line with small white spots on either side of the line (Figure 16B). Femur has white apical spots (Figure 16C). Tibia is entirely dark, and hind tarsi have white bands on tarsomeres 1-4. Tarsomere 5 is white (Figure 16C). Proboscis is dark and male palpi have two white rings on the shaft (Figure 16D). According to Edwards (1941), this species was observed in Ghana, Sudan, Kenya, Malawi and present-day south-central Africa. More recently, occurrences have been observed in Nigeria, South Africa, Burkina Faso, Central African Republic, and Senegal (GBIF Secretariat, 2023). Our observation marks a new record of occurrence in Zambia.

**FIGURE 16.**
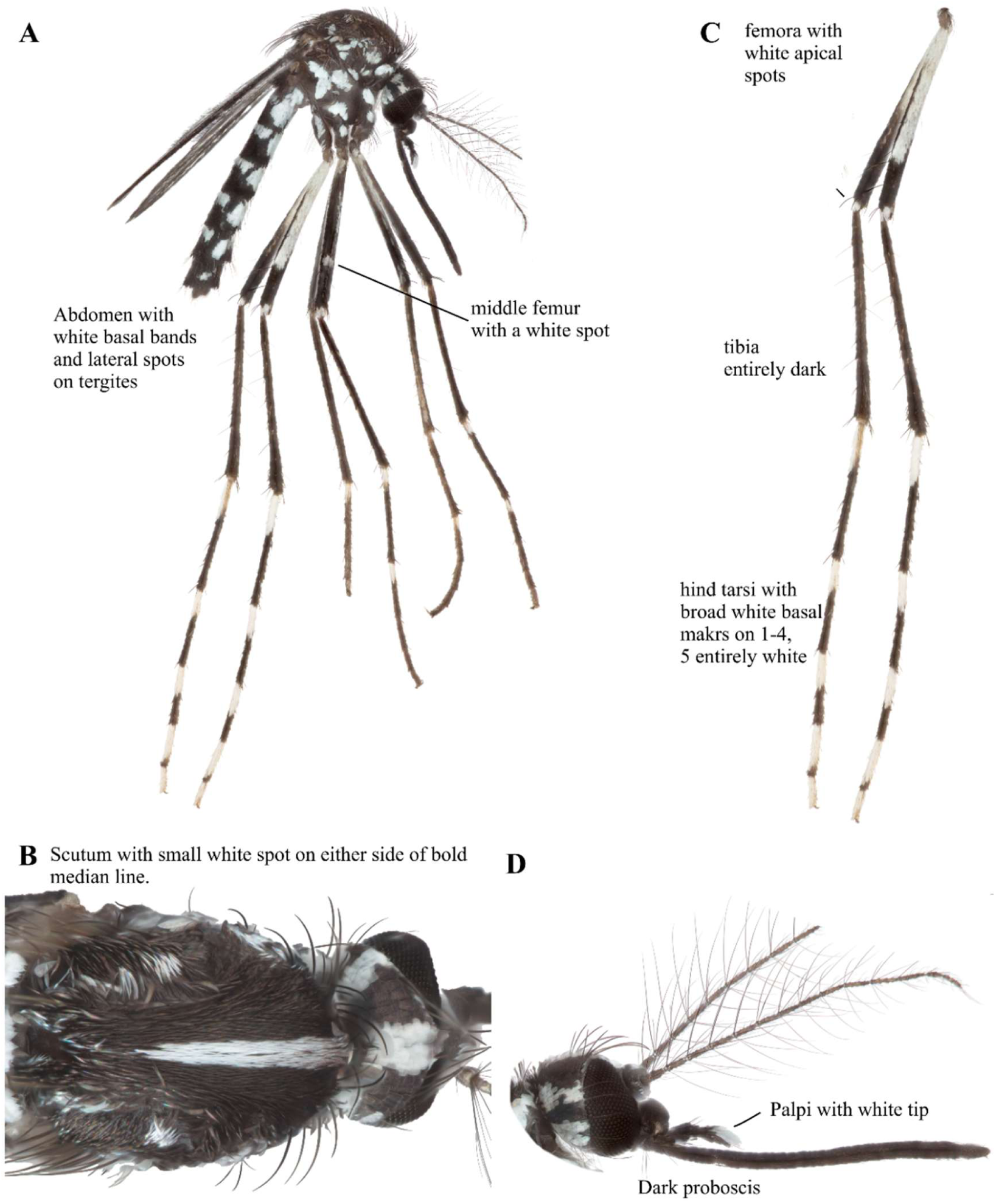
*Ae. unilineatus* male focus-stacked images. A: Lateral view. B: Scutum. C: Legs. D: Head.

### *Anopheles (Anopheles) coustani* s.l. Laveran 1900 (Diptera: Culicidae)

*Anopheles coustani* is a species complex (Gillies and Coetzee 1987). It has shaggy maxillary palpi with pale bands. Legs are not speckled, and most of the 3rd, 4th, and 5th tarsal segments are white (Gillies and Coetzee 1987). This species has been previously documented in Macha, Zambia (Kent 2006), but was not detected in the January 2024 collection trip, and thus high-resolution images are not currently available for this species. Its distribution is spread through western, southern, and parts of eastern Africa, with the highest occurrence observed in Cameroon, Nigeria, Madagascar, Sao Tome & Principe, Zambia, and South Africa (GBIF Secretariat, 2023).

### *Anopheles (Anopheles) ziemanni* Grünberg 1902 (Diptera: Culicidae)

*Anopheles ziemanni* is a brown mosquito with spotted wings (Figure 17A). It has no discernable markings on the pleuron (Figure 17B). Legs are not speckled (Figure 17C). There is a pale streak on hind tibia 1-3 times as long as broad and the hind tarsomere 2 has apical narrow and pale bands (Figure 17C). Its palpi are shaggy with pale bands (Figure 17D). Wing is not used for discerning this species, but the stem of vein 6 is white with two medium size dark bands on the 2^nd^ half of vein 6 (Figure 17E). This species has been observed in Cameroon, Central African Republic, Gabon, Gambia, Kenya, Senegal, Botswana, Burkina Faso, Morocco, South Africa and Zambia (GBIF Secretariat, 2023). We have observed this species in Macha, Zambia.

**FIGURE 17.**
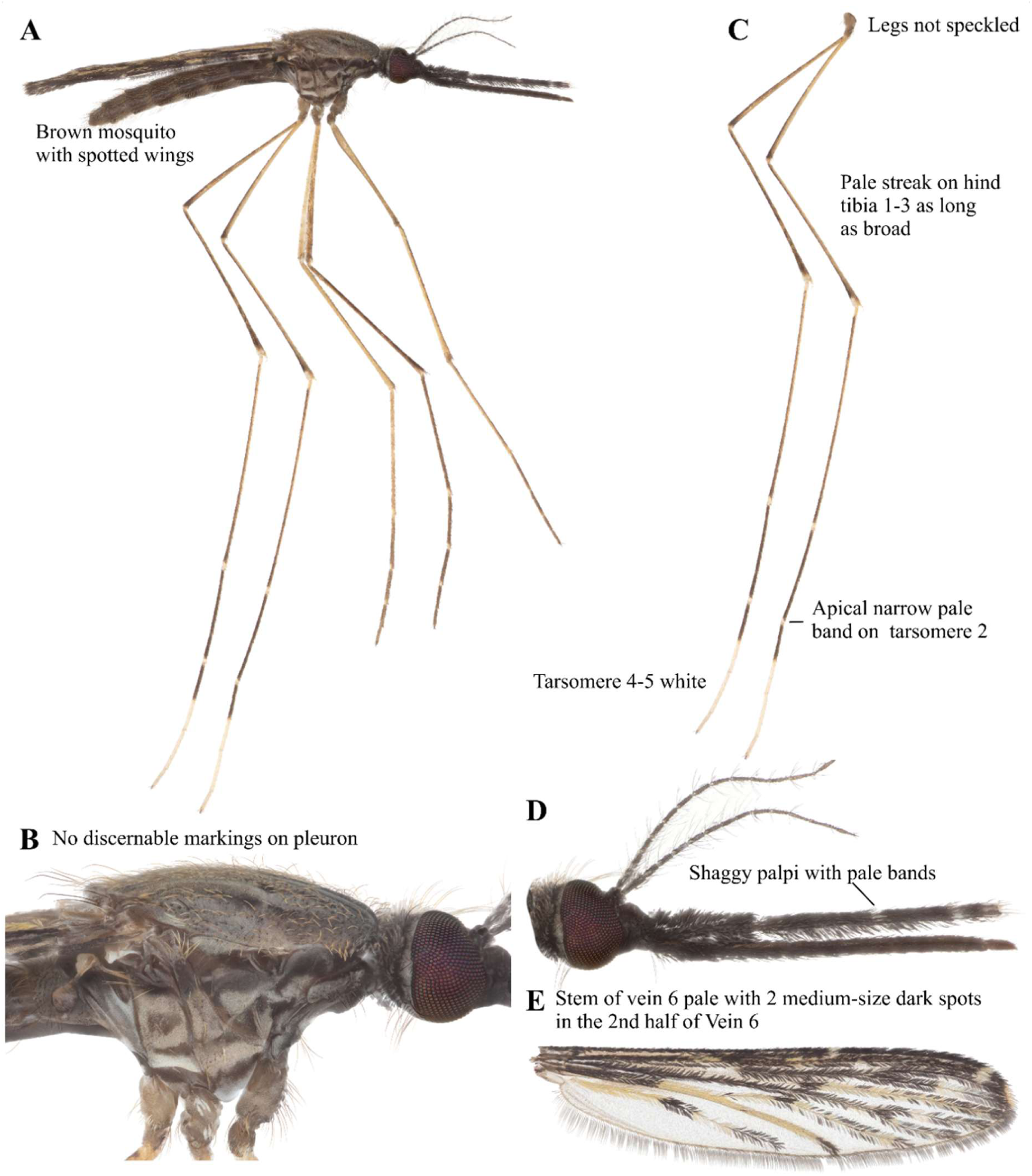
*An. ziemanni* focus-stacked images. A: Lateral view. B: Pleuron. C: Legs. D: Head. E: Wing.

### *Anopheles (Cellia) arabiensis* Patton 1905 (Diptera: Culicidae)

*Anopheles arabiensis* is a brown mosquito with spotted wings and speckled legs (Figure 18A). The scutum has no discernible markings for this species (Figure 18B). Legs are speckled and tarsomere 4-5 are not white (Figure 18C). Palpi have 3 pale bands (Figure 18D). It has a diagnostic white interruption on the R1 wing vein of the third black area (Figure 18E). This species has been previously documented in Macha, Zambia (Kent 2006). It has a wide distribution throughout eastern and western Africa, most notably, Tanzania, Uganda, Sudan, South Africa, Mali, Madagascar, Burkina Faso, Kenya, Zimbabwe and Ethiopia (GBIF Secretariat, 2023).

**FIGURE 18.**
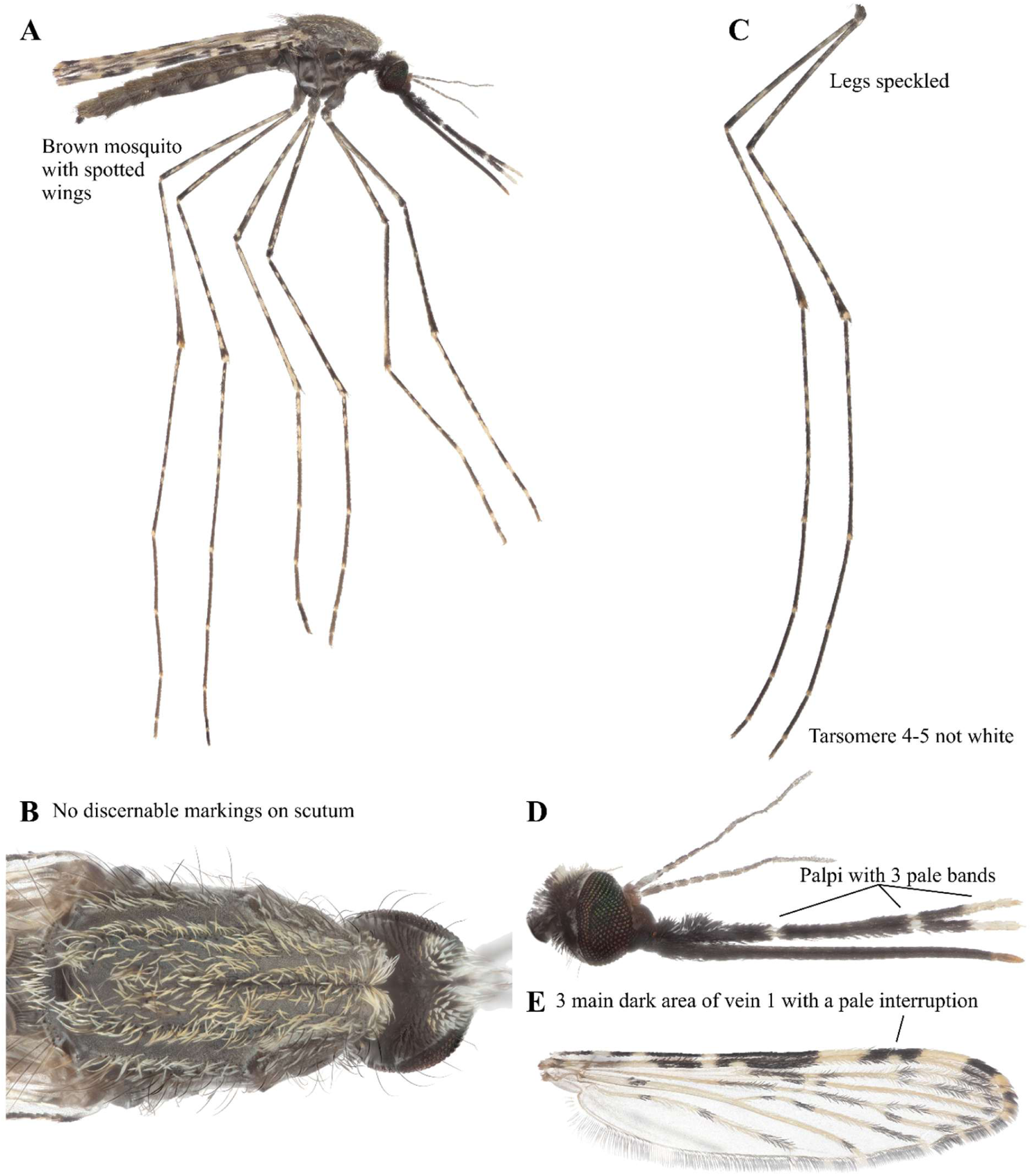
*An. arabiensis* focus-stacked images. A: Lateral view. B: Scutum. C: Legs. D: Head. E: Wing.

### *Anopheles (Cellia) funestus* Giles 1990 (Diptera: Culicidae)

*Anopheles funestus* is a species complex composed of morphologically identical members (Coetzee 2020). The palpi are ∼2X longer than the head. It has solid black legs. This species has been previously documented in various parts of western, eastern, southern and central parts of Africa (Edwards 1941, GBIF Secretariat, 2023). It was detected in Macha, Zambia (Kent 2006), but was not detected in the January 2024 collection trip, and thus high-resolution images are not currently available for this species complex or its listed members.

### *Anopheles (Cellia) longipalpis* Theobald 1903 (Diptera: Culicidae)

*Anopheles longipalpis* is a brown mosquito with spotted wings like other *Anopheles* (Figure 19A). The scutum has no discernable markings and typically is not used for *Anopheles* species identification (Figure 19B). Legs have lighter bands on joints (Figure 19C). It has palpi nearly 3-4 times longer than the head (Figure 19D). Wings of this species are not used for species identification (Figure 19E). Edwards (1941) has reported this species occurring in Kenya, Zimbabwe, Malawi and South Africa. More recently, it has been observed in Zambia and Côte d’Ivoire (Kent 2006, GBIF Secretariat, 2023).

**FIGURE 19.**
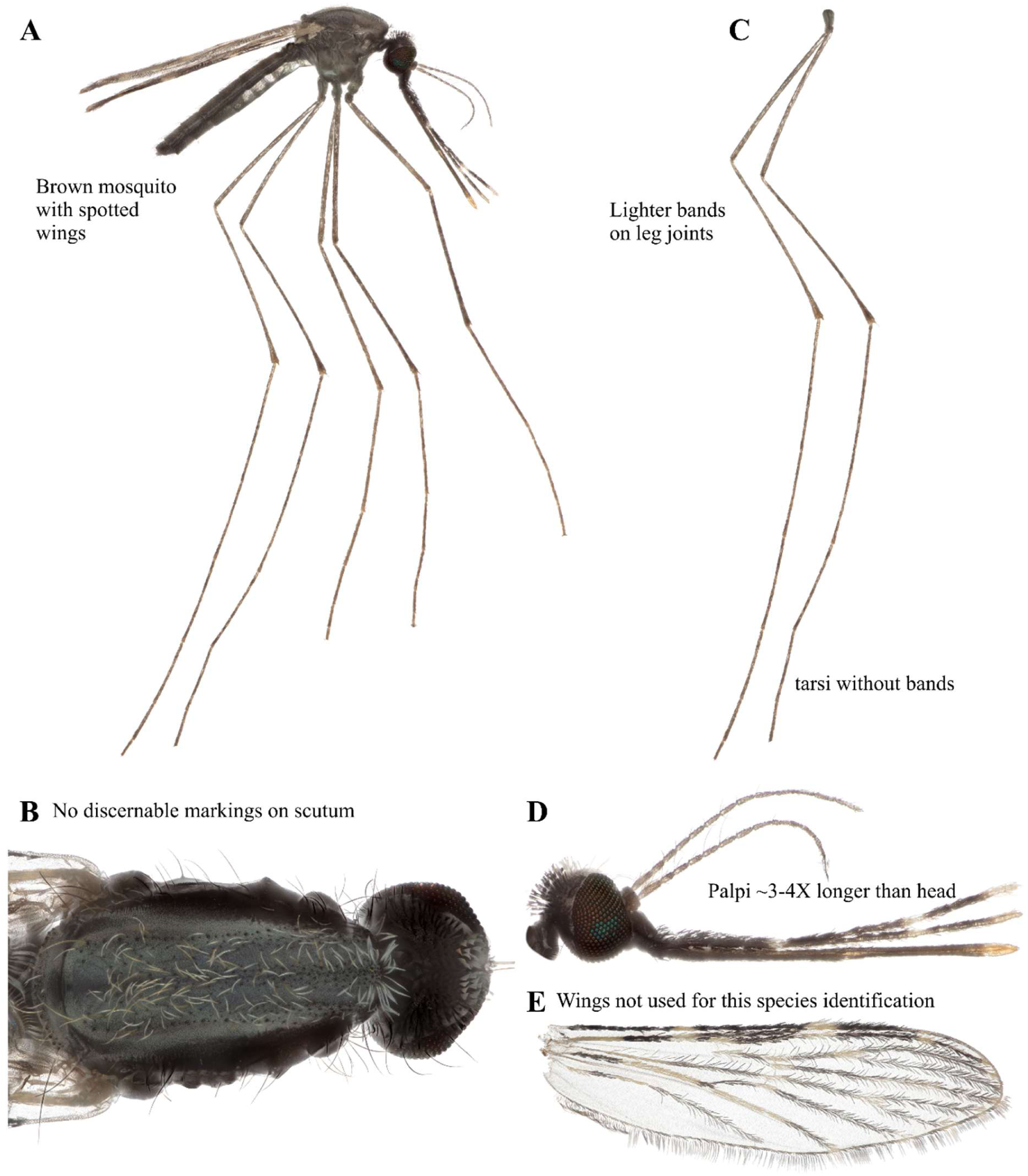
*An. longipalpis* focus-stacked images. A: Lateral view. B: Scutum. C: Legs. D: Head. E: Wing.

### *Anopheles (Cellia) maculipalpis* Giles 1902 (Diptera: Culicidae)

*Anopheles maculipalpis* does not have laterally projecting tufts of scales in the abdomen (Figure 20A). A large dark spot is present on the side of the scutum near the post-spiracular area (Figure 20B). It has a larger than typical triangular dark spot near the scutellum (Figure 20B). The femora and the tibia are speckled (Figure 20C). Tarsi 1 has dark and pale bands approximately the same length; tarsi 2-5 are white (Figure 20C). Palpi have three pale bands usually with some speckling (Figure 20D). The wing has two pale spots on the R1 vein in the second black area (Figure 20E). This species was previously recorded in Ghana, Nigeria, Congo, Malawi, Uganda, Kenya, Tanganyika, Zanzibar, Angola, Zimbabwe, South Africa, Madagascar, and Mascarenes (Edwards1941, GBIF Secretariat, 2023). This is a new record of occurrence in Macha, Zambia.

**FIGURE 20.**
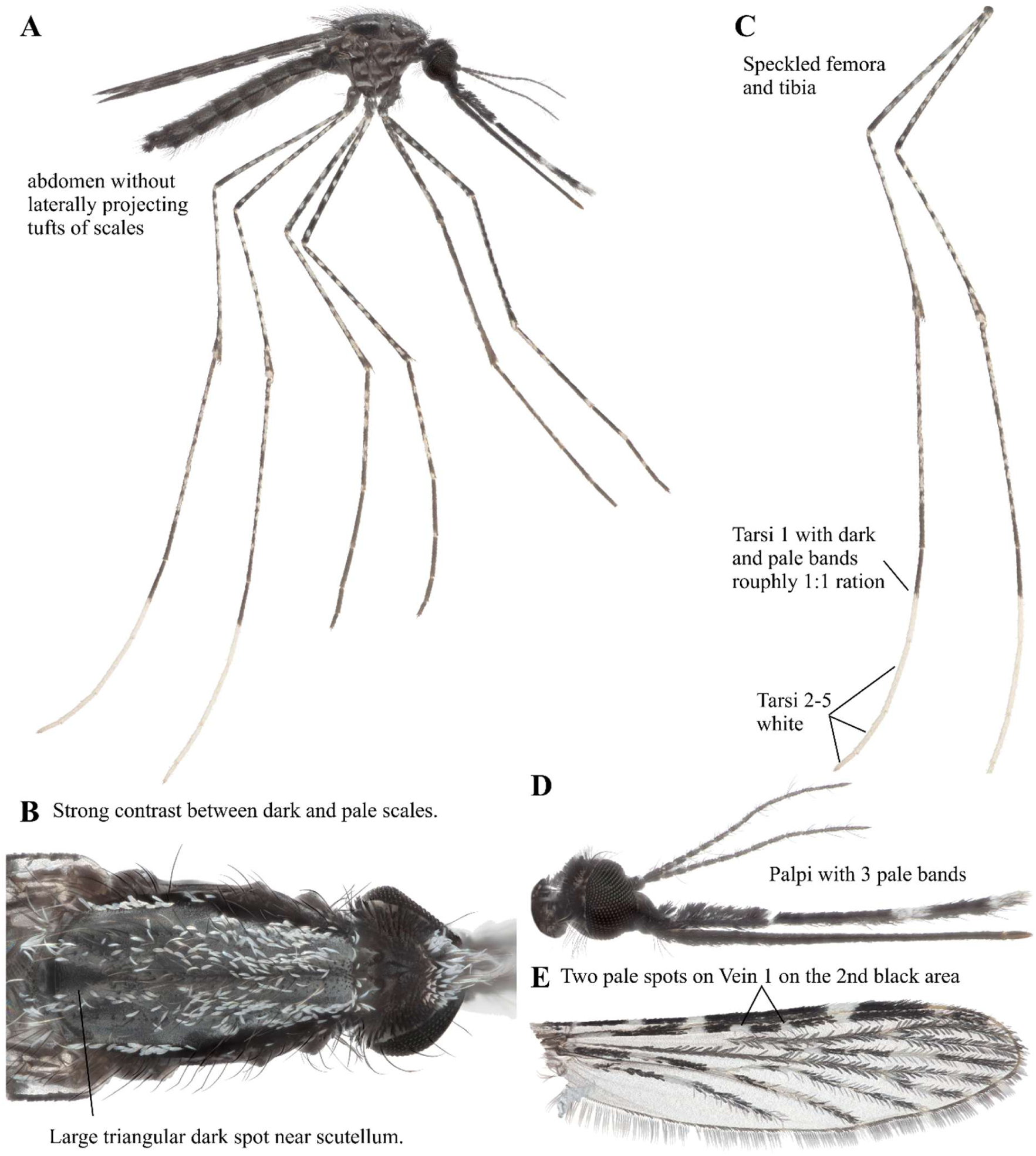
*An. maculipalpis* focus-stacked images. A: Lateral view. B: Scutum. C: Legs. D: Head. E: Wing.

### *Anopheles (Cellia) cf. pharoensis* Theobald 1901 (Diptera: Culicidae)

*Anopheles cf. pharoensis* is a brown mosquito with spotted wings (Figure 21A). It has laterally projecting tufts on the abdomen (Figure 21A). Scutum is not used for species identification, but there are two dark spots in the middle of the scutum and one dark spot near the scutellum (Figure 21B).

**FIGURE 21.**
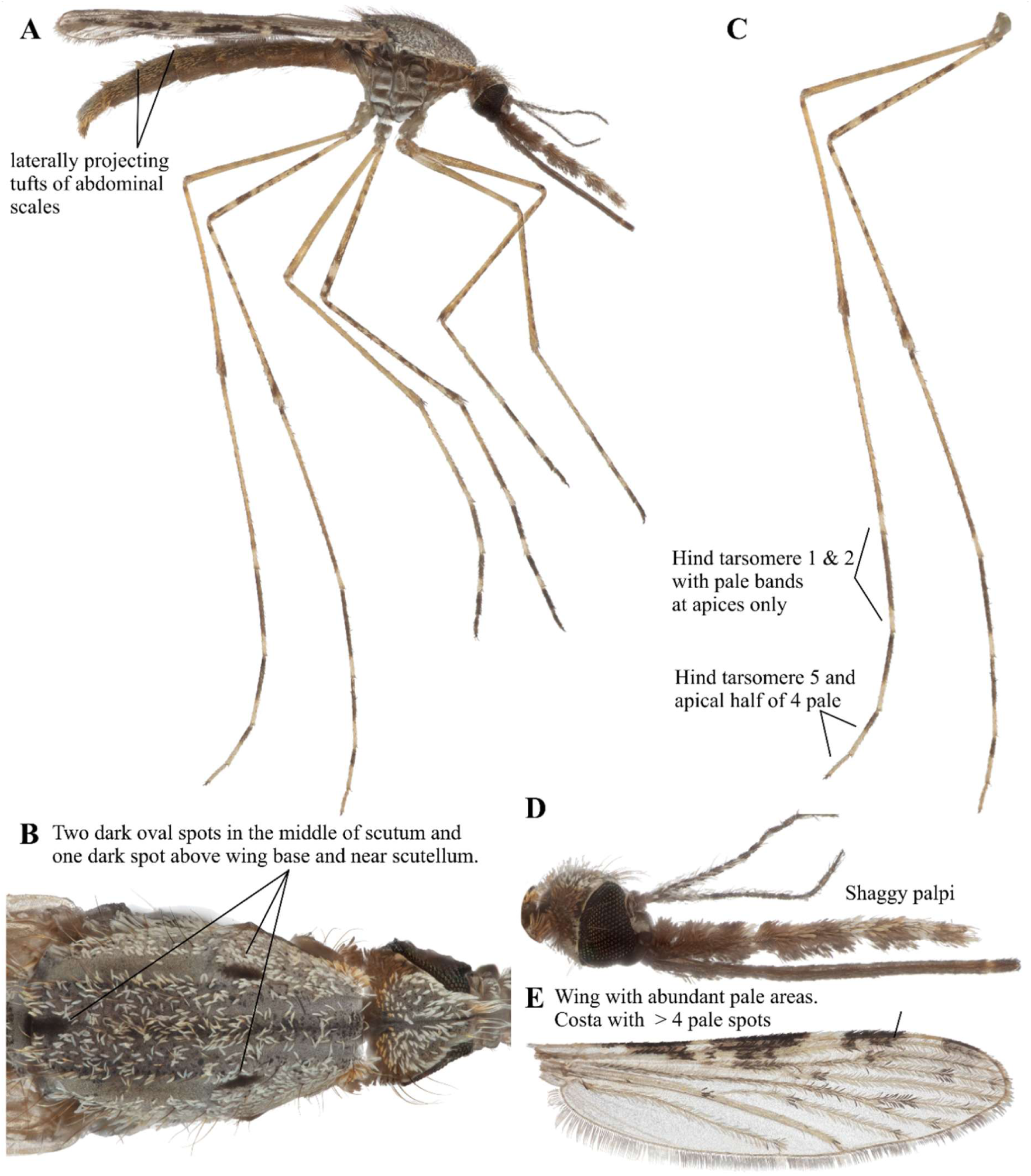
*An. cf. pharoensis* focus-stacked images. A: Lateral view. B: Scutum. C: Legs. D: Head. E: Wing.

Hind tarsomere 1-4 have pale bands, and tarsomere 5 is entirely pale (Figure 21C). Its palpi are shaggy (Figure 21D). Wing has abundant pale areas, and the costa has at least 4 pale spots (Figure 21E). This is a new record of occurrence in Macha, Zambia.

### *Anopheles (Cellia) pharoensis* Theobald 1901 (Diptera: Culicidae)

*Anopheles pharoensis* (Figure 22) shares very similar morphological features to *An. cf. pharoensis* (Figure 21). The COI sequence of this specimen has 95% identity as *An. cf. pharoensis* specimen shown in Figure 21. The distribution of this species has previously been noted in the Central Africa republic, Ethiopia, Gambia, Sierra Leone, Ghana, Nigeria, Congo, Sudan, Uganda, Kenya, Tanganyika, Angola, Zimbabwe, Malawi, Mozambique, South Africa, and Madagascar (Edwards1941, GBIF Secretariat, 2023). This is a new record of occurrence in Macha, Zambia.

**FIGURE 22.**
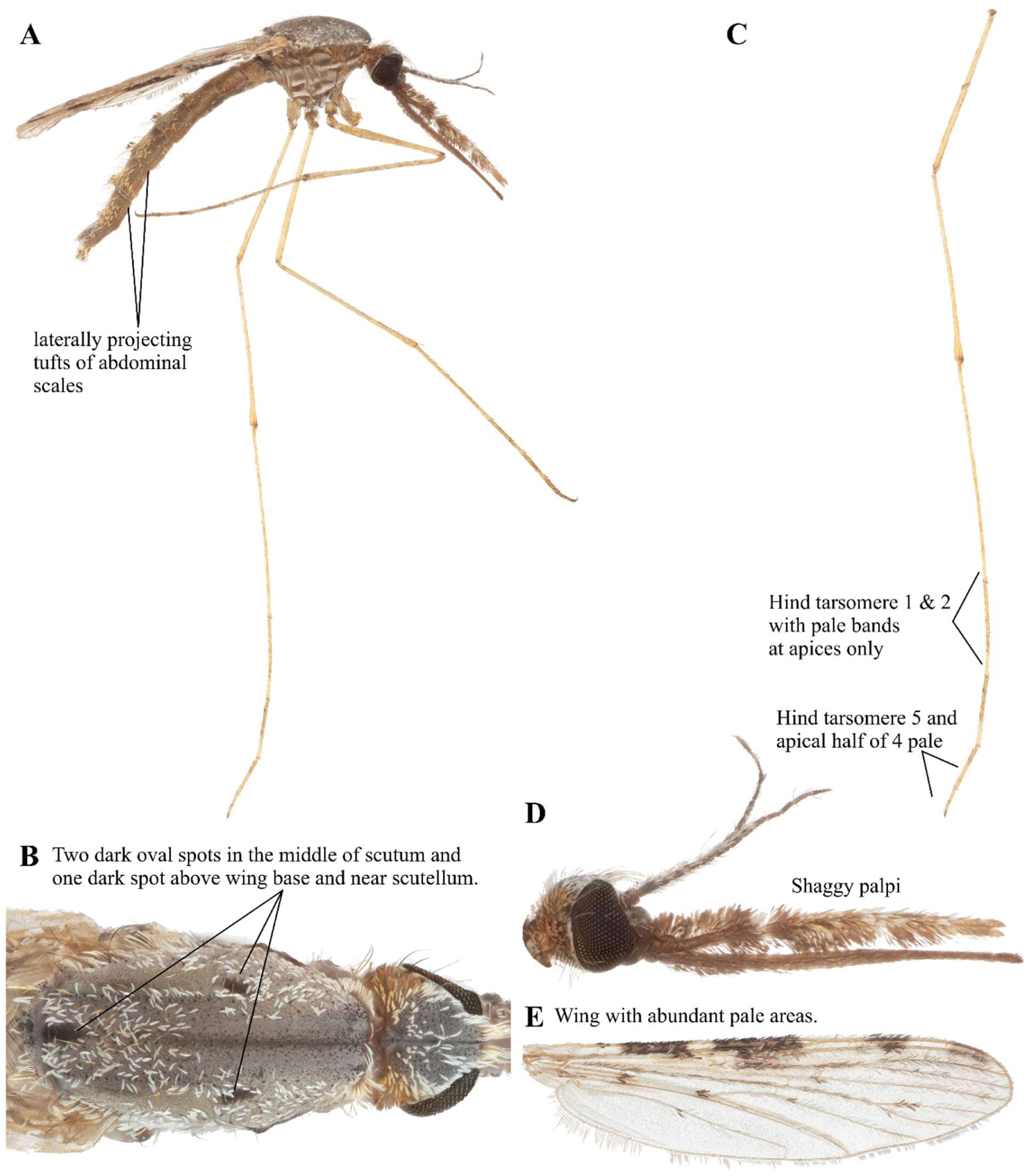
*An. pharoensis* focus-stacked images. A: Lateral view. B: Scutum. C: Legs. D: Head. E: Wing.

### *Anopheles (Cellia) pretoriensis* Theobald 1903 (Diptera: Culicidae)

*Anopheles pretoriensis* does not have laterally projecting tufts of scales on the abdomen (Gillies and Coetzee 1987). Hind tarsi 4-5 are white. Hind tarsi 1 broadly pale at apex (Gillies and Coetzee 1987). Vein 1 of wing with 2 pale spots in 2^nd^ main dark area, similar to *Anopheles maculipalpis* (Coetzee 2020). According to Edwards (1941), this species was distributed in present-day Ethiopia, Eritrea, and Somalia. More recently, this species has been observed in Comoros, Nigeria, Burkina Faso, South Africa, Malawi, Madagascar, Cameroon, Mayotte, and Togo (GBIF Secretariat, 2023). While this species was documented in Macha, Zambia (Kent 2006), we were unable to collect in the January 2024 collection trip, and thus high-resolution images are not currently available for this species.

### *Anopheles (Cellia*) *rufipes* Gough 1910 (Diptera: Culicidae)

*Anopheles rufipes* does not have laterally projecting tufts of scales on the abdomen (Figure 23A). It has a similar scutal pattern to *An. maculipalpis,* but no obvious dark spot near scutellum (Figure 23B). Femora and tibia are not speckled (Figure 23C). Hind tarsi 4-5 are white. This species has smooth palpi with three pale bands (Figure 23D). It has two pale spots on the R1 vein in the second black area with no pale interruption in the third black area (Figure 23E). This species has been previously documented in in Nigeria, Congo, Sudan, Uganda, Kenya, Zimbabwe, and South Africa (Edwards 1941). It has been more recently observed in Macha, Zambia (Kent 2006, GBIF Secretariat, 2023). Notably this specimen matches well (>99%) with Botswana and Zambia *An. rufipes* COI sequences (Genbank Accession #PP815715 and LC473605, respectively) but has lower similarity (∼94%) to those from Mali (Genbank Accession #MK586041) and others from Huestis et al. (2019) or Botswana (Genbank Accession # PP747847). This could reflect the differences between *An. rufipes* (Gough 1910) and *An. broussesi* (Edwards, 1929) that were recently elevated to species by Harbach and Wilkerson (2023).

**FIGURE 23.**
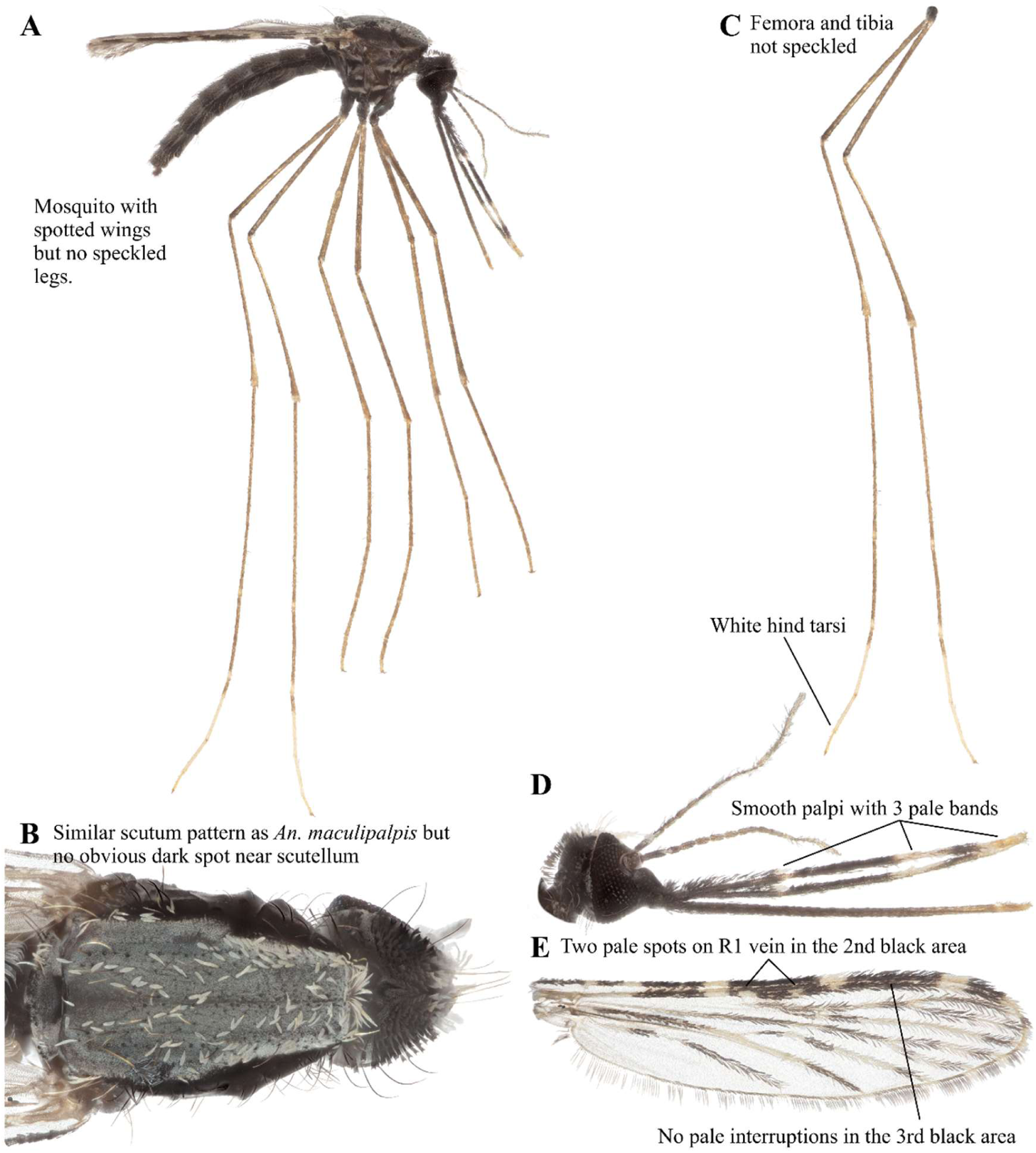
*An. rufipes* focus-stacked images. A: Lateral view. B: Scutum. C: Legs. D: Head. E: Wing.

### *Anopheles (Cellia*) *squamosus* Theobald 1901 (Diptera: Culicidae)

*Anopheles squamosus* is a dark mosquito with laterally projecting tufts on the abdomen (Figure 24A). There are no discernible markings on the scutum (Figure 24B). The femora and tibia are speckled, and the tarsi are banded (Figure 24C). It has shaggy palpi (Figure 24D). The wing has an upper branch on vein 2 either entirely dark apart from the apex or with a few scattered pale scales only (Figure 24E). According to Edwards (1941), the distribution of this species includes Sierra Leone, Ghana, Nigeria, Congo, Sudan, Uganda, Kenya, Tanganyika, Zanzibar, Zimbabwe, Zambia, central-north Africa, parts of South Africa, and Madagascar (GBIF Secretariat, 2023).

**FIGURE 24.**
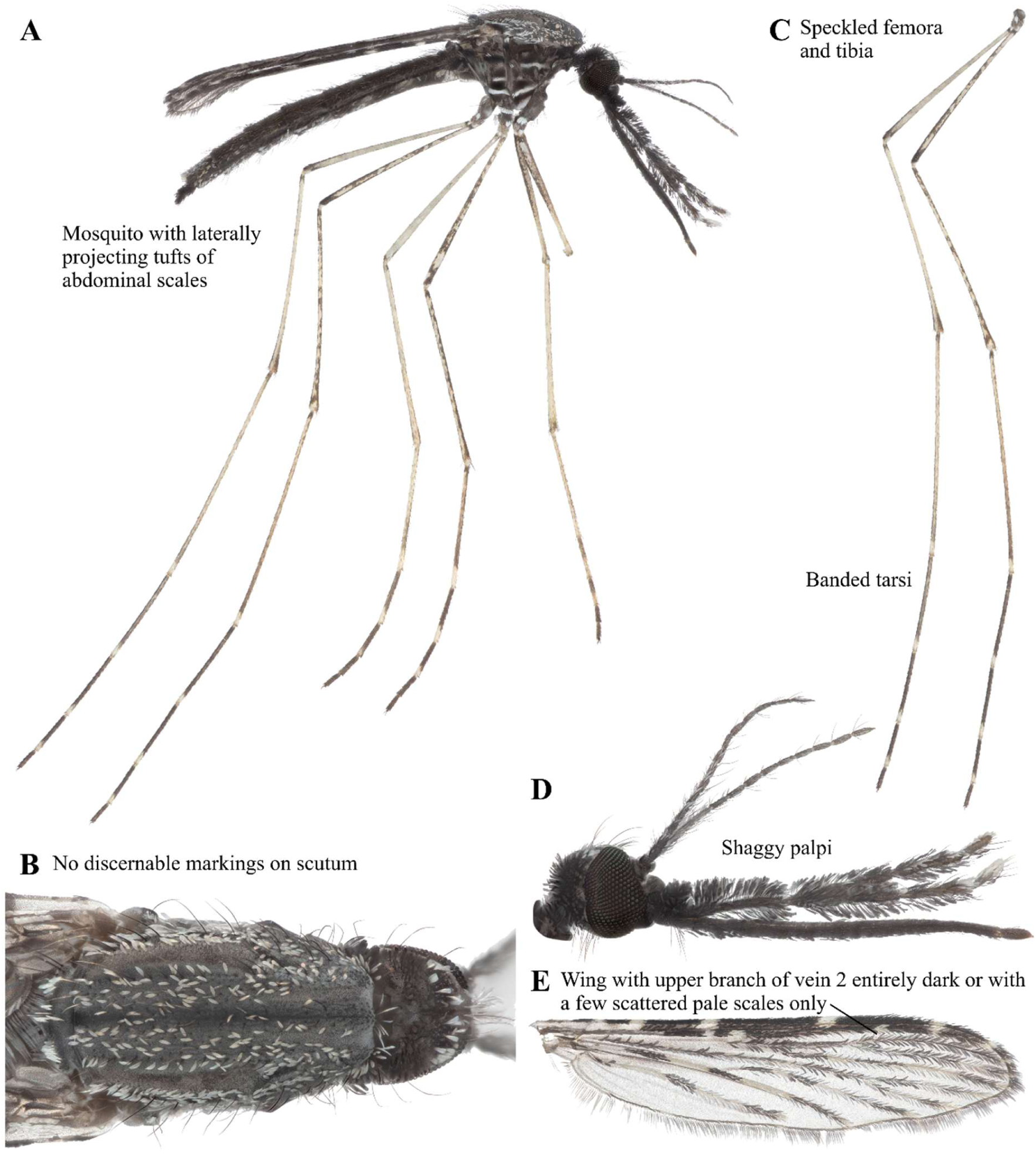
*An. squamosus* focus-stacked images. A: Lateral view. B: Scutum. C: Legs. D: Head. E: Wing.

### *Culex (Culex) antennatus* Becker 1903 (Diptera: Culicidae)

*Culex antennatus* is a small brown mosquito (Figure 25A). It has an unbanded abdomen, but with lateral patches on terminal segments and white abdominal sterna (Figure 25A). No post spiracular scale patches. Its brown scutum has no discernible markings (Figure 25B). It has dark, unbanded legs (Figure 25C). The proboscis is all dark (Figure 25D). This species has been previously documented in all parts of Africa including the Congo, Sudan, Uganda, Nigeria, Chad, Ethiopia, Madagascar, and Zanzibar (Edwars 1941, GBIF Secretariat, 2023). This species has been observed in Macha, Zambia (Kent 2006).

**FIGURE 25.**
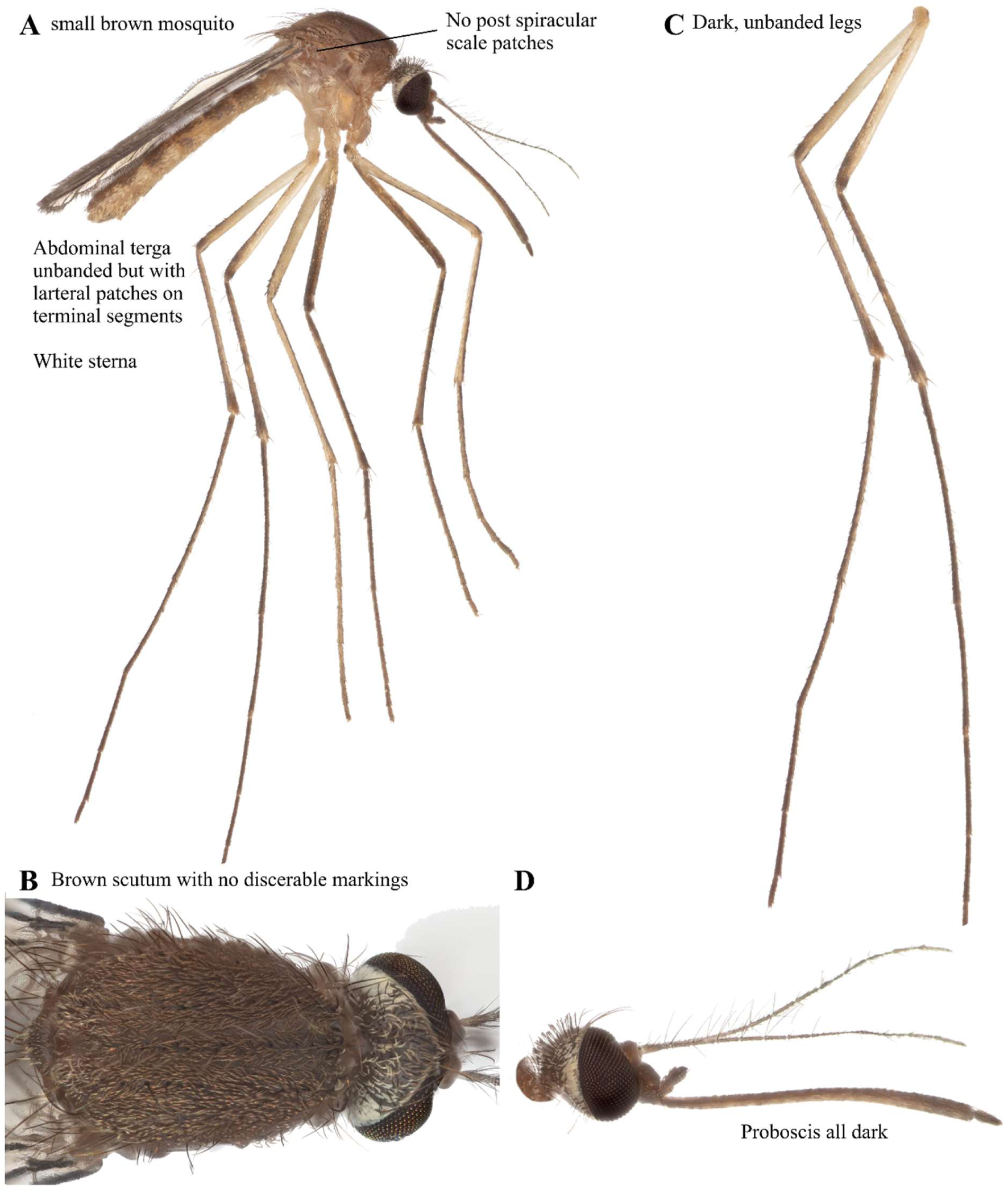
*Cx. antennatus* focus-stacked images. A: Lateral view. B: Scutum. C: Legs. D: Head.

### *Culex (Culex) litwakae* Harbach 1985 (Diptera: Culicidae)

*Culex litwakae* is a small, dark brown mosquito that looks similar to *Cx. antennatus* (Figure 26A). Scale patch on mesepimeron at the level of the upper mesokatepisternal patch and lower mesokatepisternal scales separate from upper scale patch (Figure 26A). The scutum has golden brown scales, lightening slightly at the pre-scutellar area (Figure 26B). Hind femora are mostly pale while tarsi are dark (Figure 26C). The proboscis is dark on top and light on bottom (Figure 26D). This is a new record of occurrence in Zambia.

**FIGURE 26.**
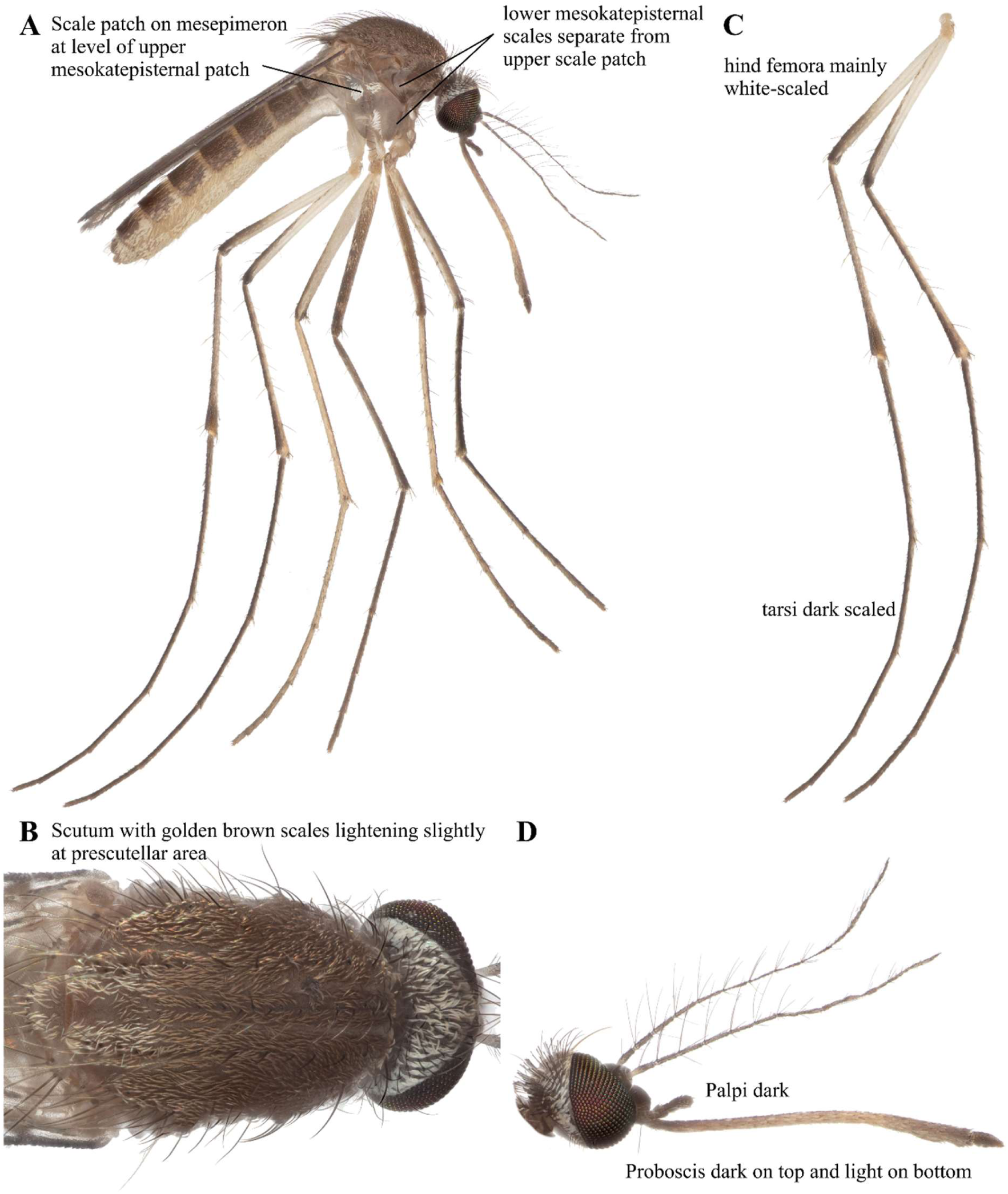
*Cx. litwakae* focus-stacked images. A: Lateral view. B: Scutum. C: Legs. D: Head.

### *Culex (Culex*) *perexiguus* Theobald 1903 (Diptera: Culicidae)

*Culex perexiguus* is a brown mosquito with unbanded dark brown legs (Figure 27A). It has a costa with a few pale scales at the base. The lower mesokatepisternal scales are separated from the upper scale patch. Its brown scutum has golden brown scales in the middle (Figure 27B). The base and venter of the femora are pale, and the apex of the tibia has a white spot (Figure 27C). The rest of the legs are dark brown (Figure 27C). The proboscis and palpi are dark (Figure 27D). While this species had been observed in Mali, South Africa, Egypt, Tunisia, Kenya, Algeria, Libya, Mozambique, Cameroon, and Morocco (GBIF Secretariat, 2023), our observation marks a new record of occurrence in Zambia.

**FIGURE 27.**
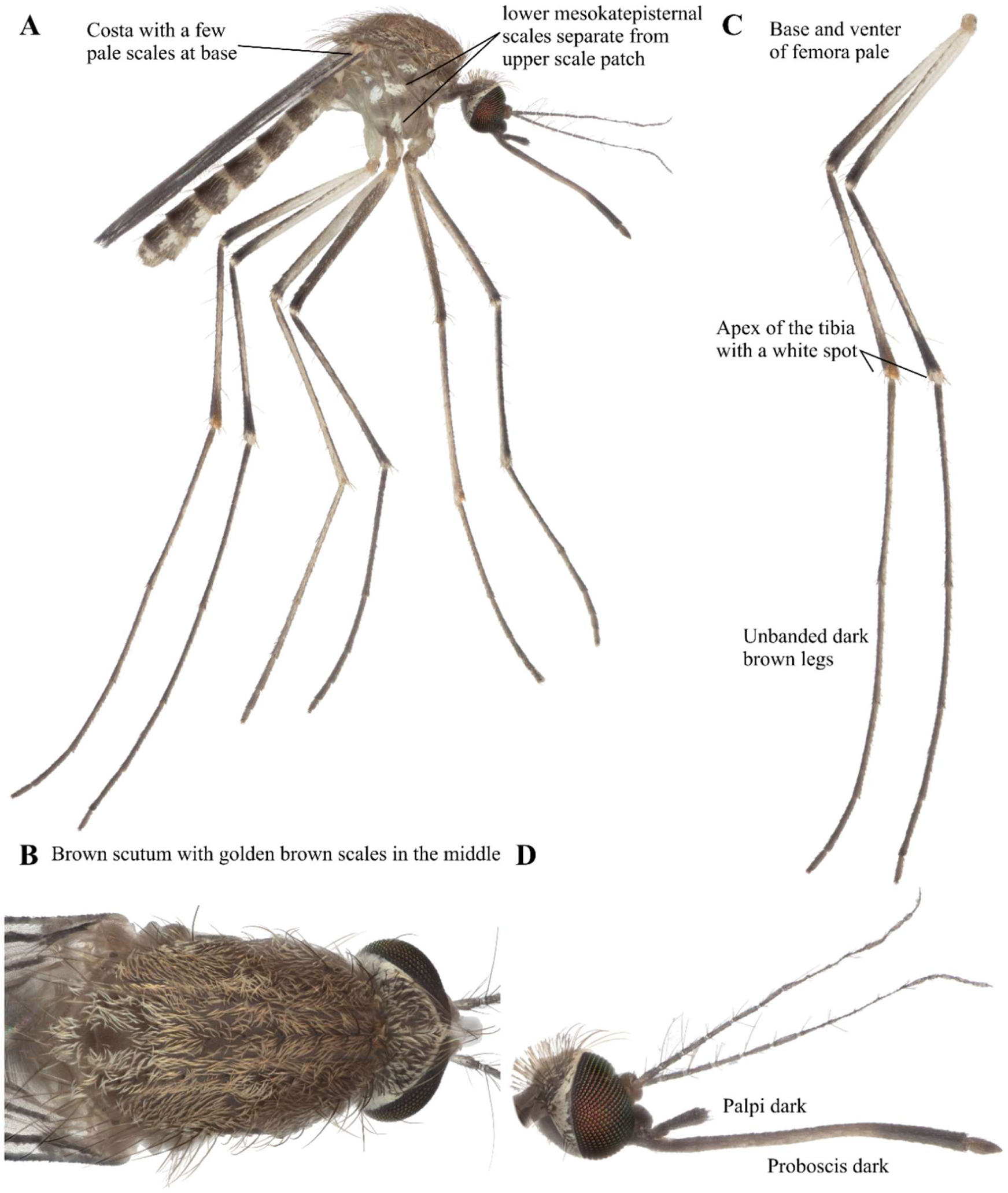
*Cx. perexiguus* focus-stacked images. A: Lateral view. B: Scutum. C: Legs. D: Head.

### *Culex (Culex*) *quinquefasciatus* Say 1823 (Diptera: Culicidae)

*Culex quinquefasciatus* has thick, half-moon shaped basal bands on terga,with mostly white sterna (Figure 28A). Scutum has pale brown scales (Figure 28B). The legs are dark and unbanded (Figure 28C). Proboscis is dark (Figure 28D), and its palpi are reported to be shorter than *Culex pipiens* (Edwards 1941). According to Edwards (1941), this is the most common domestic species spanning tropical Africa, south-west Arabia, and the Gulf of Guinea. This species has been previously documented in Macha, Zambia (Kent 2006). It has been observed in Benin, Uganda, Tanzania, Congo, South Africa, Comoros, Nigeria, France, Madagascar, and Burkina Faso (GBIF Secretariat, 2023).

**FIGURE 28.**
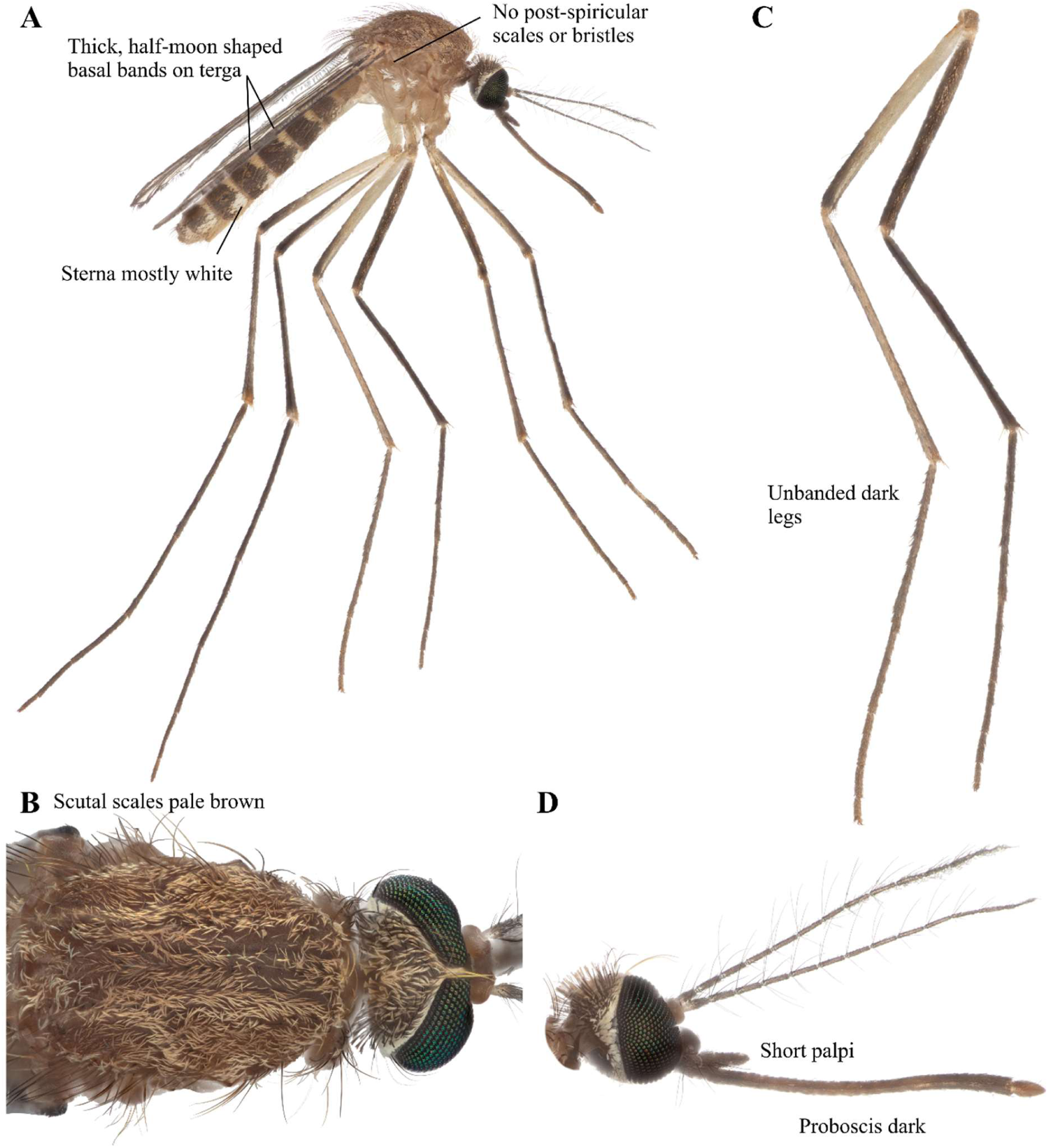
*Cx. quinquefasciatus* focus-stacked images. A: Lateral view. B: Scutum. C: Legs. D: Head.

### *Culex (Culex) telesilla* De Meillon & Lavoipierre 1945 (Diptera: Culicidae)

*Culex telesilla* has sterna with a dark apical transverse band and terga with cream-colored basolateral patches (Figure 29A). The post spiracular scales are absent (Figure 29A). Scutum has no silvery scales (Figure 29B). The hind tibia has no pale apical spots, and pale bands are absent in tarsi (Figure 29C). Proboscis is entirely dark (Figure 29D). The occurrence of this species has been noted in South Africa, Cameroon, and Nigeria (GBIF Secretariat, 2023). Our observation marks a new record of occurrence in Zambia.

**FIGURE 29.**
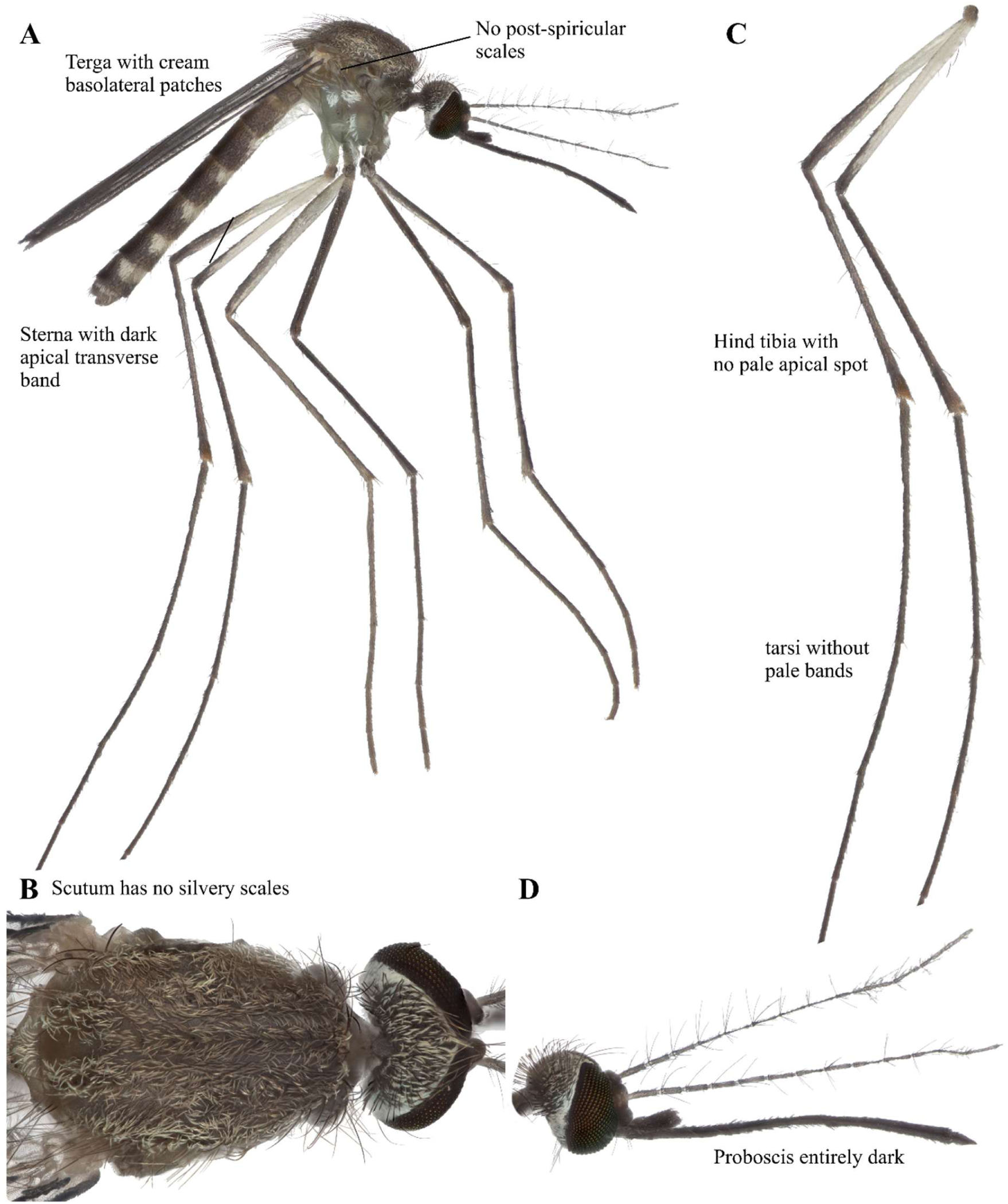
*Cx. telesilla* focus-stacked images. A: Lateral view. B: Scutum. C: Legs. D: Head.

### *Culex (Culex*) *univittatus* Theobald 1901 (Diptera: Culicidae)

*Culex univittatus* has post spiracular scales (Figure 30A). It usually has dark markings on the abdominal sterna (Figure 30A). It has mostly whitish decumbent scales (Figure 30B). The hind tibia has a diagnostic longitudinal stripe (Figure 30C; Edwards 1941, Jupp 1996). Its proboscis is dark but paler in the middle (Figure 30D). Palpi are dark and 1/6 the length of the proboscis (Figure 30D). According to Edwards (1941), this species is commonly distributed throughout the Ethiopian region and Mediterranean region. Occurrences have been reported in Egypt, South Africa, Burkina Faso, Madagascar, Nigeria, Morocco, Senegal, Cameroon, Kenya, and Benin (GBIF Secretariat, 2023). This species has been previously documented in Macha, Zambia (Kent 2006).

**FIGURE 30.**
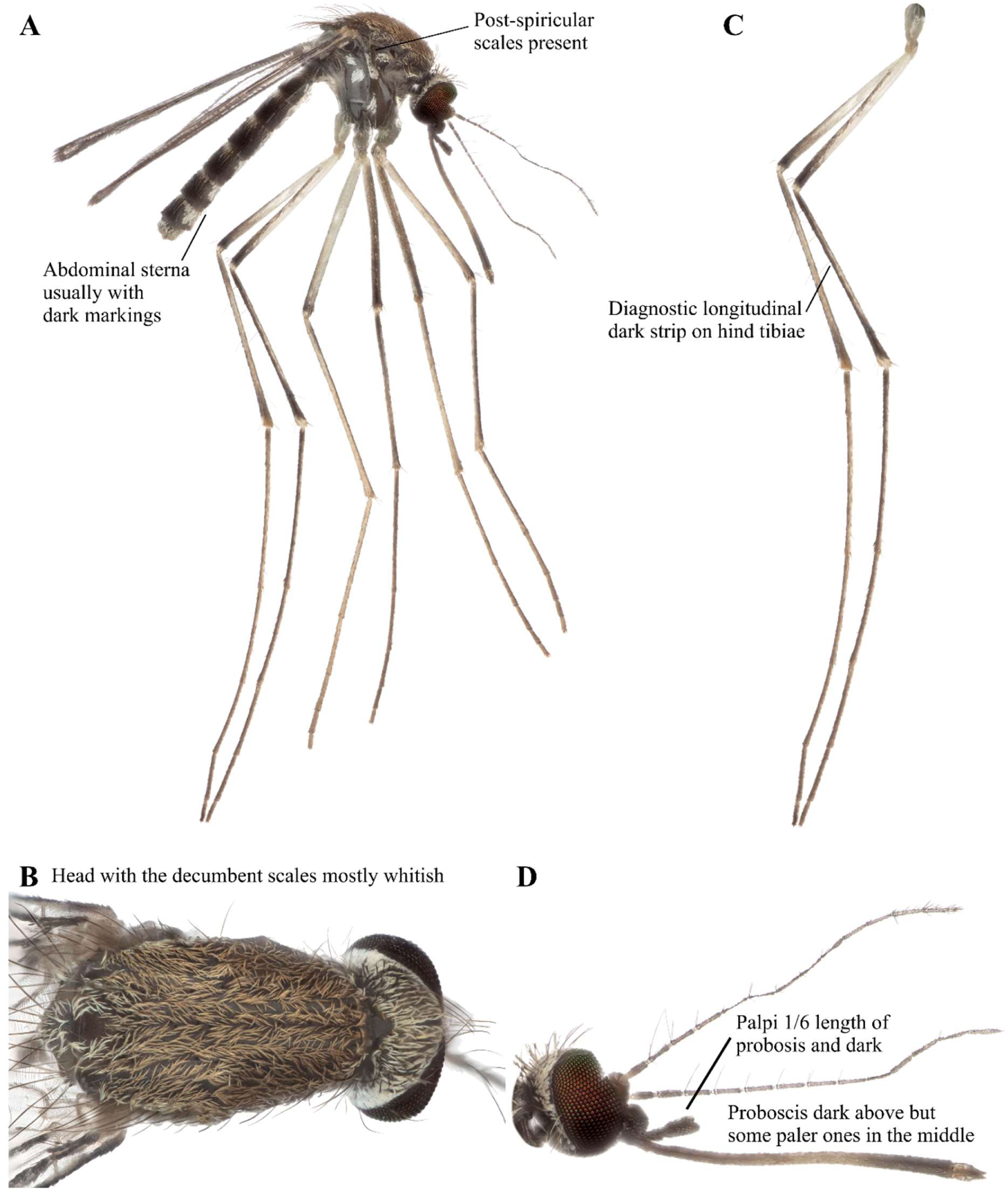
*Cx. univittatus* focus-stacked images. A: Lateral view. B: Scutum. C: Legs. D: Head.

### *Culex (Culciomyia*) *cinereus* Theobald 1901 (Diptera: Culicidae)

*Culex cinereus* is a greyish brown mosquito (Figure 31A) resembling *Cx. nebulosus,* but *Cx. cinereu*s is larger on average (Edwards 1941). A row of white scales lines the posterior edge of mesokatepisternum (Figure 31A). Setae and scales on all coxae. A row of wide, pale scales lines the orbital margin of the eyes (Figure 31B). Legs are not used for diagnosing this species, but the tibia and tarsi are dark (Figure 31C). Proboscis is all dark (Figure 31D). This species has been previously observed in Sierra Leone, Ghana, Nigeria, Morocco, Cameroon, South Africa, Eritrea, Côte d’Ivoire, Mali, Birkin Faso, and Congo (Edwards 1941, GBIF Secretariat, 2023). Our observation marks a new record of occurrence in Macha, Zambia.

**FIGURE 31.**
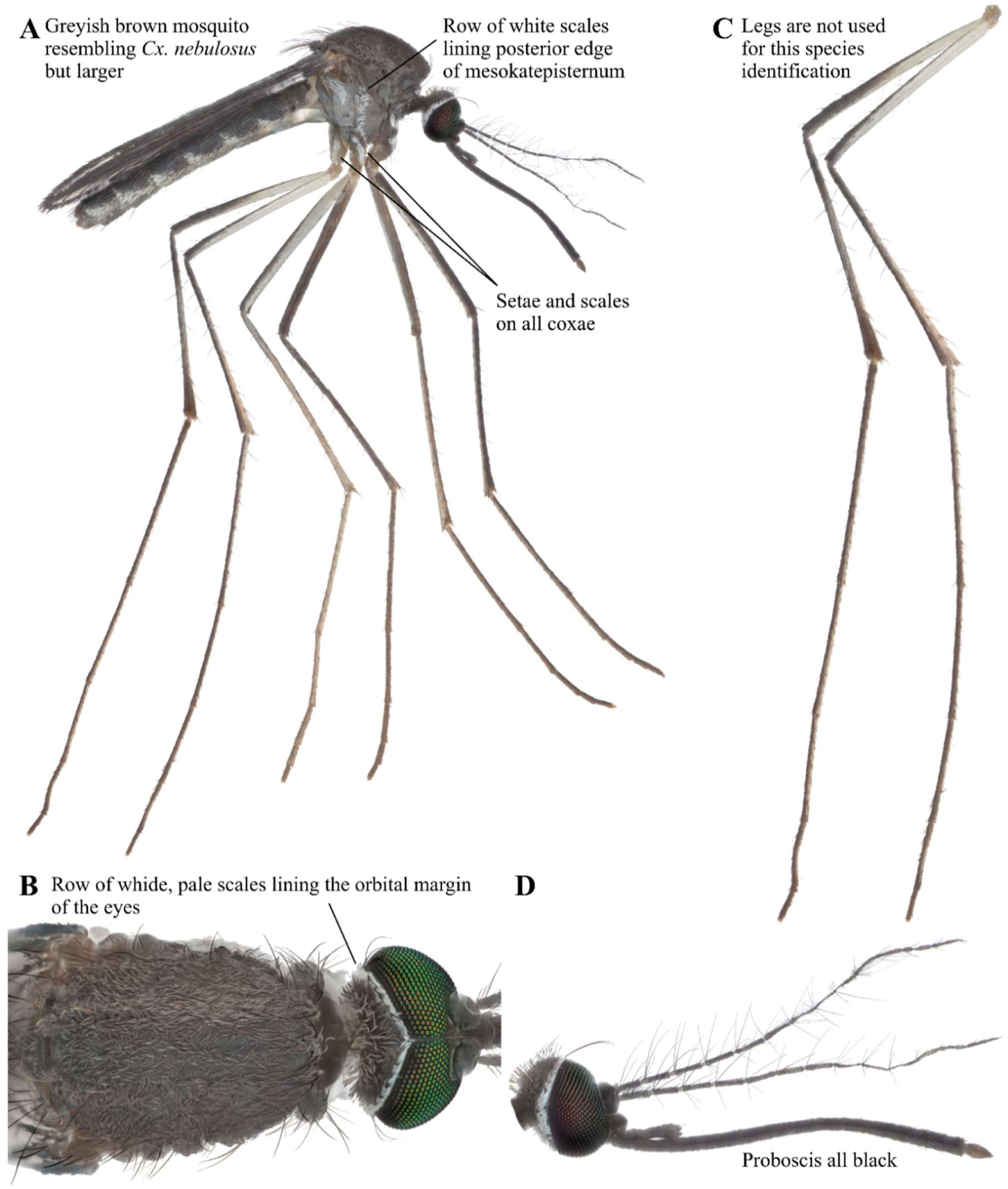
*Cx. cinereus* focus-stacked images. A: Lateral view. B: Scutum. C: Legs. D: Head.

### *Culex (Culiciomyia) nebulosus* Theobald 1901 (Diptera: Culicidae)

*Culex nebulosus* is a grayish brown mosquito with few markings (Figure 32A). It has a vertical row of white scales lining the posterior edge of the mesokatepisternum (Figure 32A). Scutum has a grayish appearance (Figure 32B). Femora, tibia, and tarsi without pale markings or bands (Figure 32C). A row of wide, pale scales lines the orbital margin of the eyes (Figure 32D). Palpi and Proboscis are dark (Figure 32D). Edwards (1941) noted that this species is one of the most widely distributed mosquitoes in Africa, occurring in Sierra Leone, Ghana, Nigeria, Cameroon, Sudan, Congo, Uganda, and Madagascar. This species has been previously documented in Macha, Zambia (Kent 2006). Recent records also documented this species in Burkina Faso, Senegal, Mali, Benin, and Comoros (GBIF Secretariat, 2023).

**FIGURE 32.**
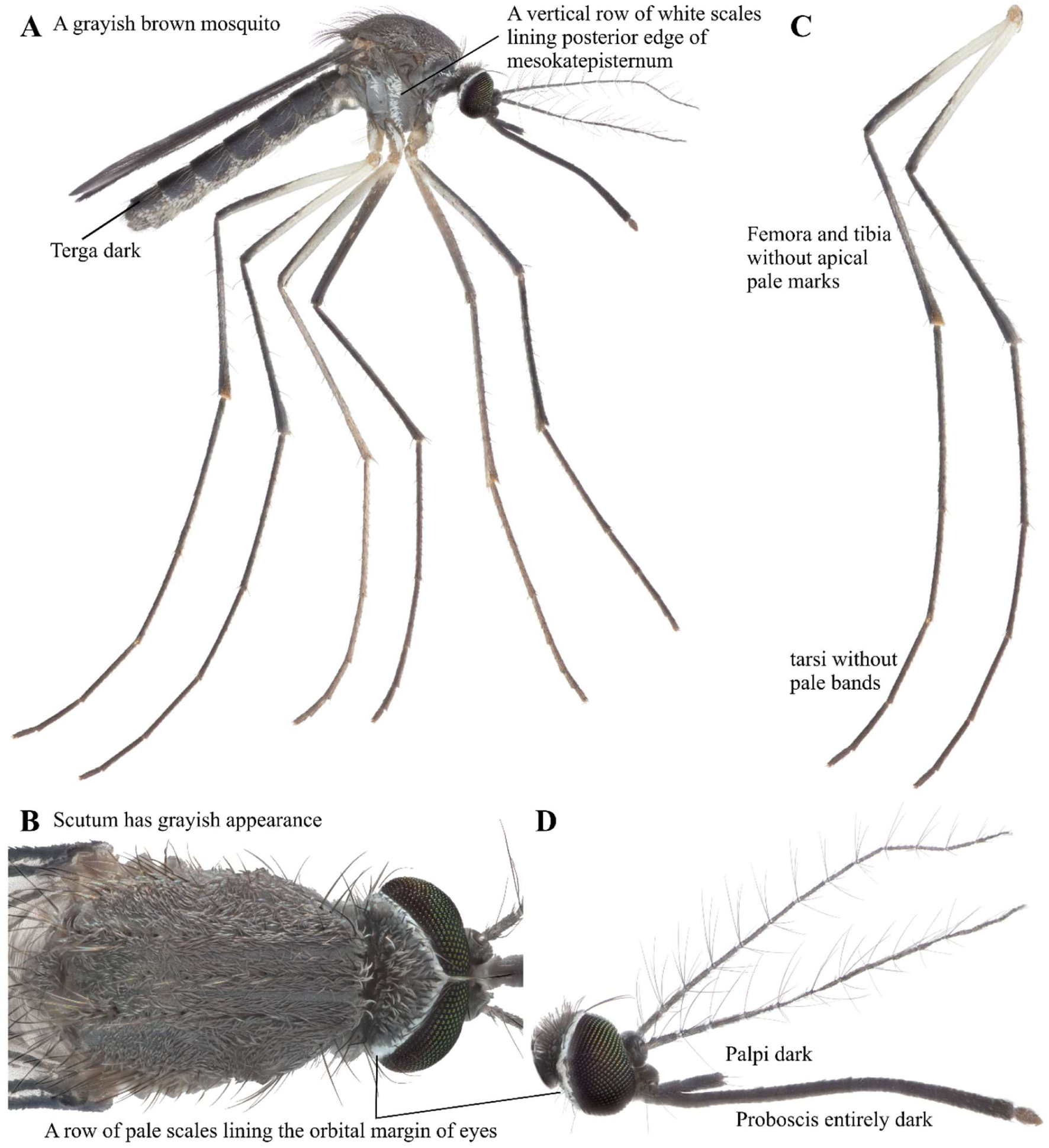
*Cx. nebulosus* focus-stacked images. A: Lateral view. B: Scutum. C: Legs. D: Head.

### *Culex (Eumelanomyia) inconspicuosus* Theobald 1908 (Diptera: Culicidae)

*Culex inconspicuosus* has few ornamental features (Figure 33A). Terga are dark with faint apical pale marks (Figure 33A). Pleural scales are absent (Figure 33A). Scutum is also inornate (Figure 33B). The tibiae and tarsi are dark (Figure 33C). Dark palpi and proboscis (Figure 33D). According to Edwards (194), this species had been observed in Ghana, Sudan, Uganda, Zimbabwe, and parts of South Africa. Recently, occurrences have been reported in Côte d’Ivoire, Burkina Faso, Nigeria, Congo, Cameroon, Benin, and Mali as well (GBIF Secretariat, 2023). However, this is a new record of occurrence in Zambia.

**FIGURE 33.**
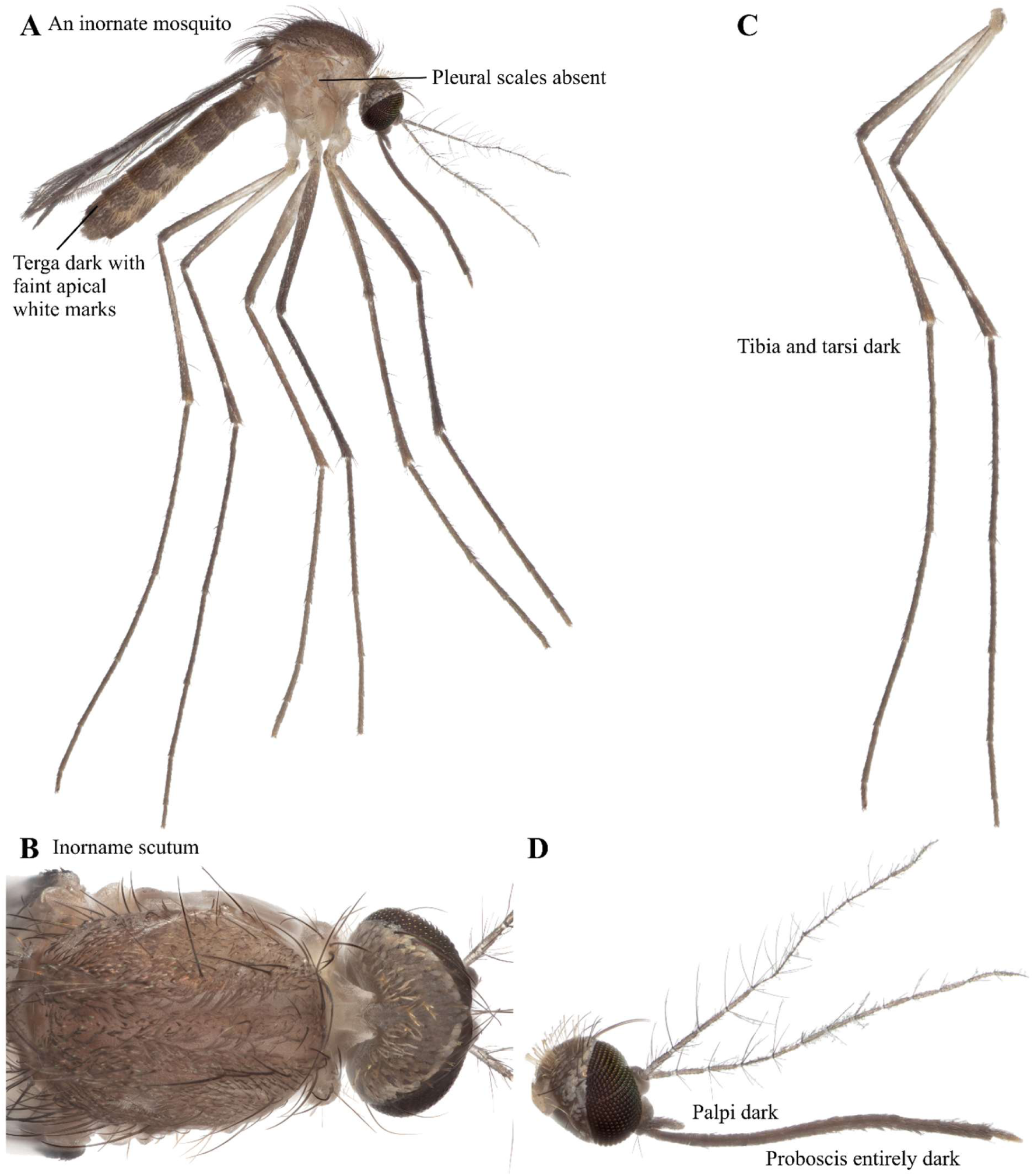
*Cx. inconspicuosus* focus-stacked images. A: Lateral view. B: Scutum. C: Legs. D: Head.

### *Culex (Lasioconops) poicilipes* Theobald 1903 (Diptera: Culicidae)

*Culex poicilipes* has no lower mesanepimeral setae (Figure 34A). Scutum is mostly dark brown or blackish with transversal band behind the middle (Figure 34B). The proboscis has a well-defined pale band (Figure 34D). The fore and mid femora have a row of 6-12 spots (Figure 34C) (Edwards 1941, Jupp 1996). It is one of the more common mosquitoes distributed around the tropical parts of Africa (Edwards 1941). Occurrences have also been noted in Nigeria and parts of South Africa (GBIF Secretariat, 2023). This species has been previously documented in Macha, Zambia (Kent 2006).

**FIGURE 34.**
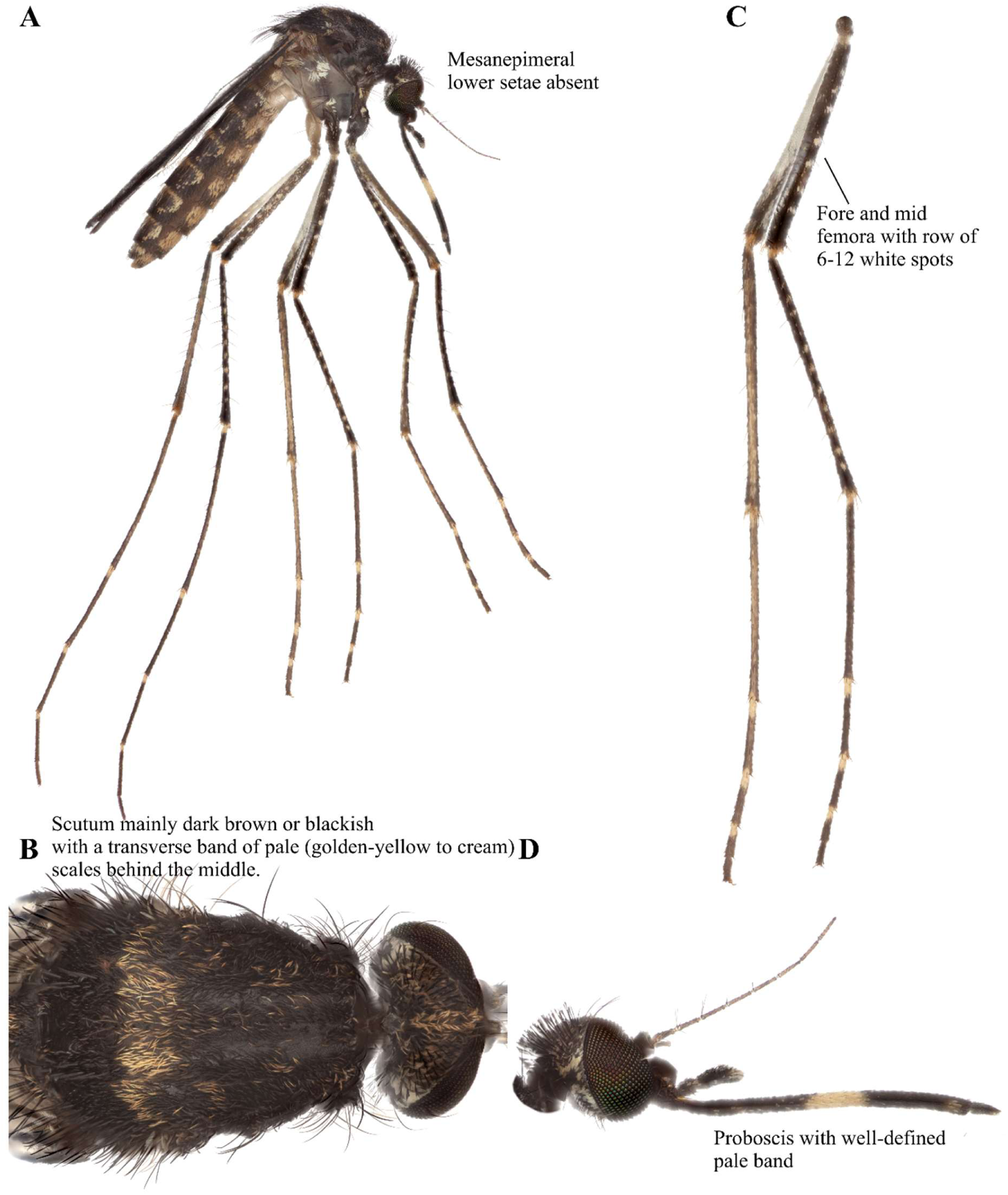
*Cx. poicilipes* focus-stacked images. A: Lateral view. B: Scutum. C: Legs. D: Head.

### *Culex (Oculeomyia*) *bitaeniorhynchus* Edwards 1901 (Diptera: Culicidae)

*Culex bitaeniorhynchus* has a largely black abdomen without a basal band but has apical lateral yellow spots on tergites 5-8 (Figure 35A). It lacks lower mesepimeral bristles. The thorax is largely brown with pale scales near the scutellum (Figure 35B). Legs are sprinkled with pale scales on femora and tibiae, and tarsi are banded (Figure 35C). Proboscis has a distinct medial pale band and pale labellum (Figure 35D). The wing is speckled and has broad, pale, and dark scales (Figure 35E). Occurrence have been recorded in parts of West, East, and South Africa, including Tanzania, Uganda, Sudan, Kenya, and Congo (Edwards 1941, GBIF Secretariat, 2023). This species has been previously documented in Macha, Zambia (Kent 2006).

**FIGURE 35.**
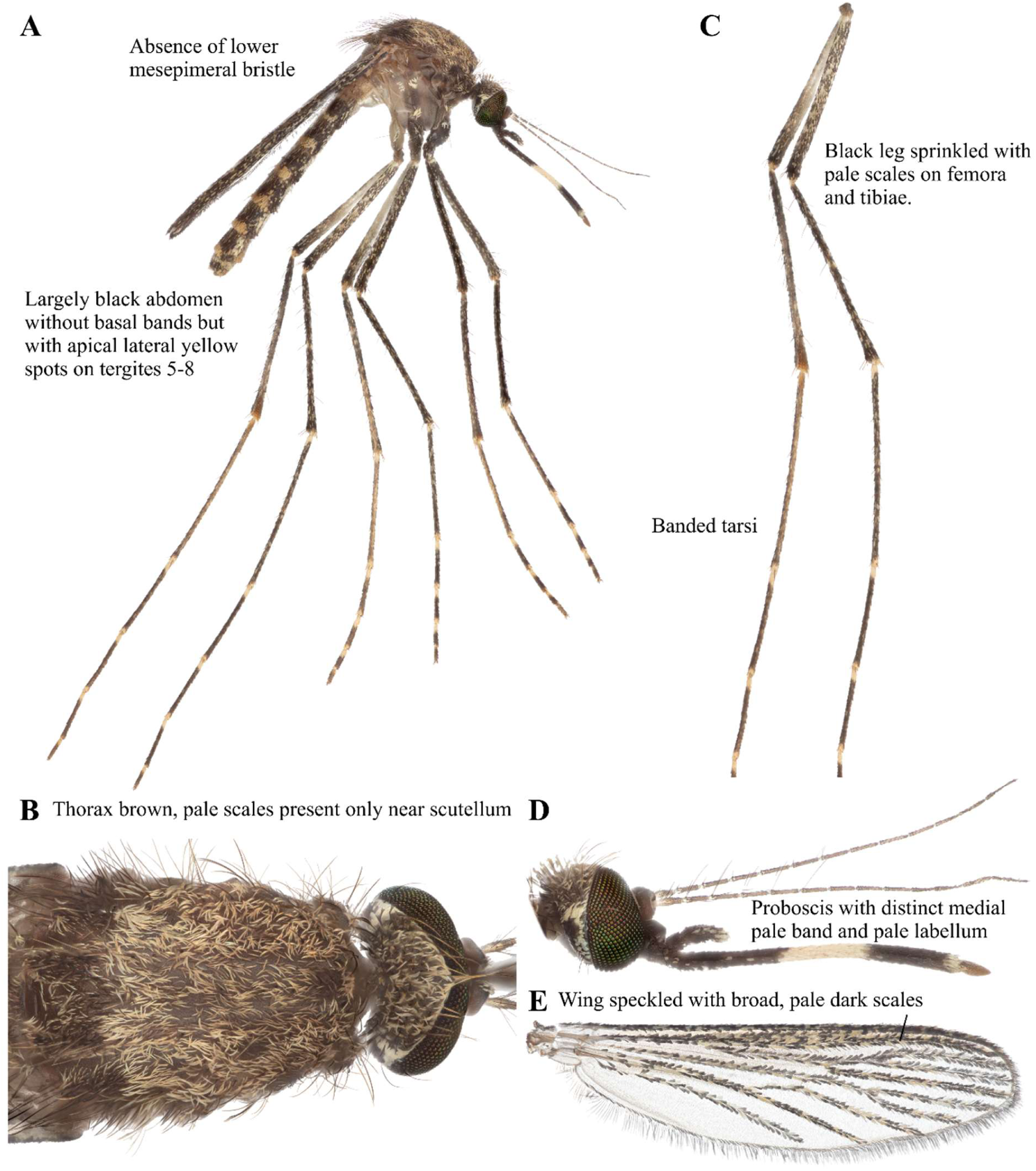
*Cx. bitaeniorhynchus* focus-stacked images. A: Lateral view. B: Scutum. C: Legs. D: Head. E: Wing.

### *Ficalbia circumtestacea* Theobald 1908 (Diptera: Culicidae)

*Ficalbia circumtestacea* has a patch of grey scales in the center of the mesokatepisternum. The palpi are 0.2x the length of the proboscis. The proboscis is slightly swollen at the tip. The first flagellomere is 3x longer than the second (Edwards 1941, Jupp 1996). This species has been previously documented in Macha, Zambia (Kent 2006), but was not detected in the January 2024 collection trip, and thus high-resolution images are not currently available for this species. Of note, this species has been reported from Benin and Burkina Faso in West Africa (Ibrahim and Granouillac 2012, GBIF Secretariat, 2023). According to Edwards (1941), the occurrence had also been noted in Sudan and Sierra Leone. The Kent (2006) record marks the first report of this species in southern African country of Zambia.

### *Ficalbia uniformis* Theobald 1904 (Diptera: Culicidae)

*Ficalbia uniformis* has a pale yellowish pleuron with no scales (Figure 36A). Terga are mainly dark with basal bands (Figure 36A). The scutum has very numerous and strong dorso-central bristles but is otherwise bare (Figure 36B). Legs are dark except for the lower and posterior surface of the femora, which are yellowish (Figure 36C). The palps and proboscis are dark (Figure 36D). The proboscis is swollen at the tip (Figure 36D). While the occurrence of this species has been noted in South Africa, our observation marks the first record of occurrence in Zambia (GBIF Secretariat, 2023). Edwards (1941) has also reported the occurrence in Ghana, Nigeria, Congo, Uganda, and Sudan.

**FIGURE 36.**
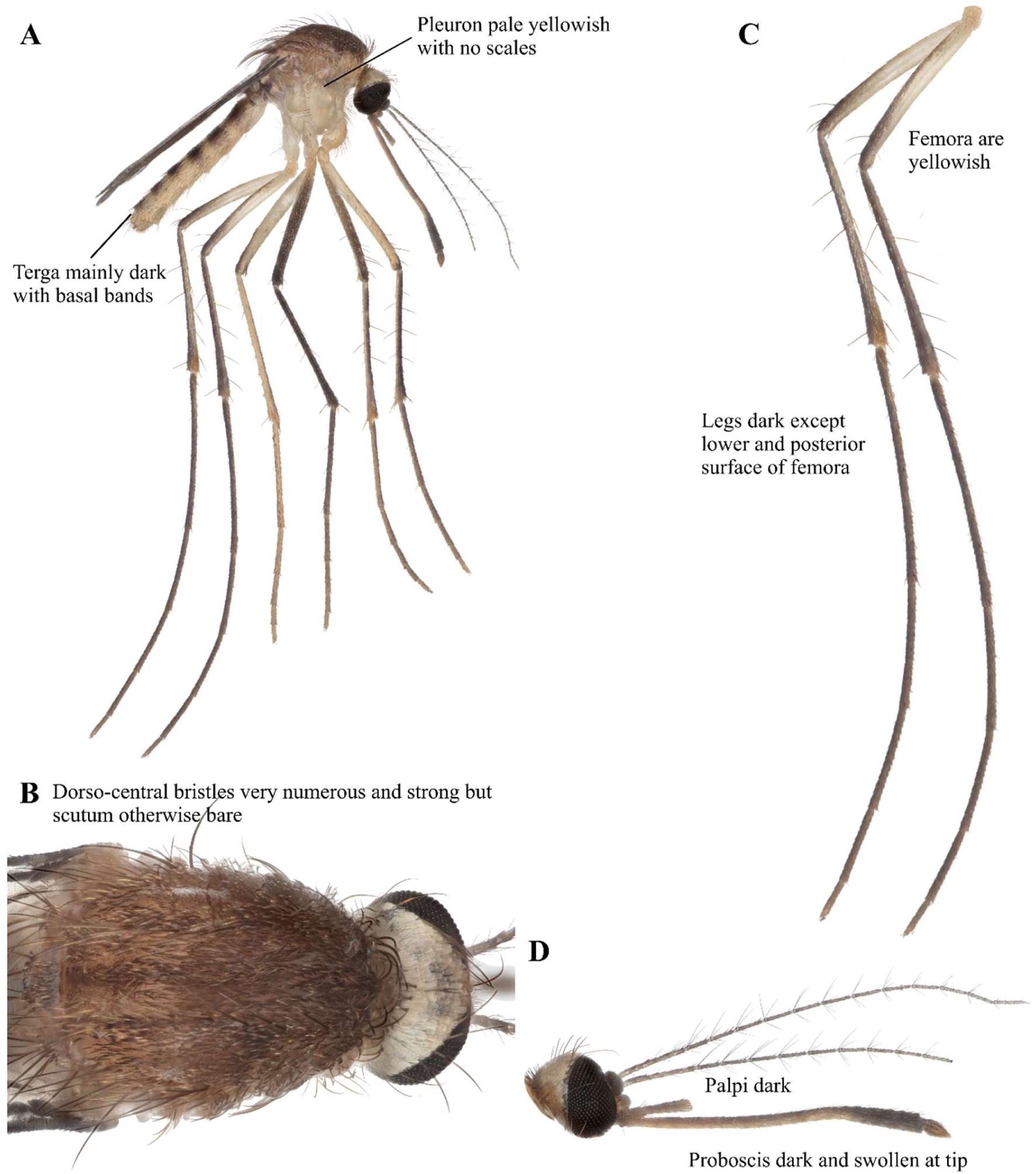
*Fi. uniformis* focus-stacked images. A: Lateral view. B: Scutum. C: Legs. D: Head.

### *Lutzia (Metalutzia*) *tigripes* De Grandpré & De Charmoy 1901 (Diptera: Culicidae)

*Lutzia tigripes* is a large mosquito with four mesanepimeral setae, apical abdominal bands and large patches of white scales on the upper part of mesepimeron (Figure 37A). Scutum has mostly brown scales with three pale spots: 2 near the middle of the scutum and a median spot towards the front margin (Figure 37B). The femora and tibia are speckled, and the tarsi do not have pale bands (Figure 37C). Proboscis is pale in the middle (Figure 37D). According to Edwards (1941), this species commonly occurs around the Ethiopian region. This species has been previously documented in Nigeria and parts of East Africa (GBIF Secretariat, 2023). The observation by Kent (2006) marks the first occurrence in Macha, Zambia.

**FIGURE 37.**
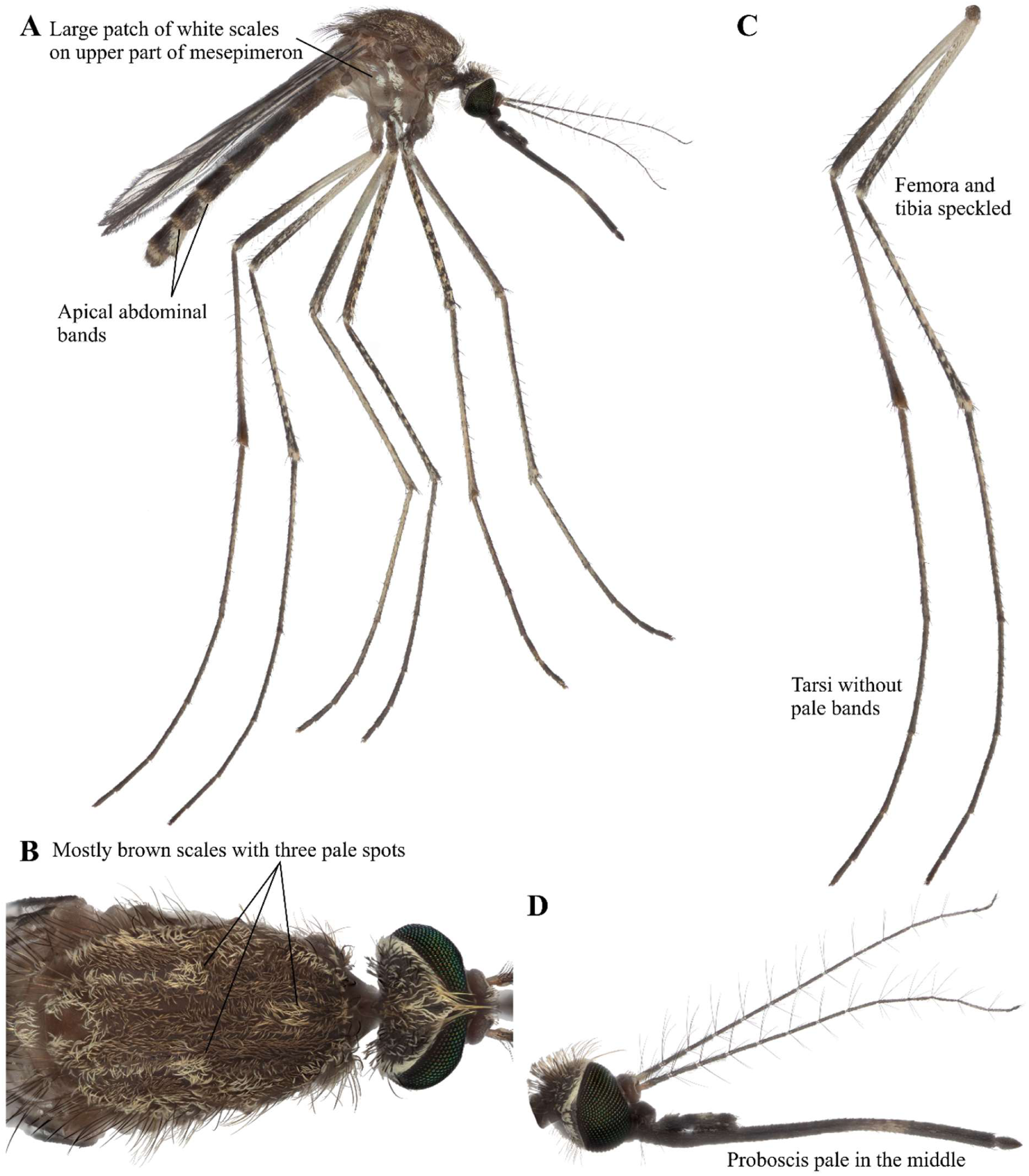
*Lt. tigripes* focus-stacked images. A: Lateral view. B: Scutum. C: Legs. D: Head.

### *Mansonia (Mansonioides) uniformis* Theobald 1901 (Diptera: Culicidae)

*Mansonia uniformis* has a blunt abdomen (Figure 38A). Scutum with bands of pale scales (Figure 38B). Confluent white spots present on hind tibiae and tarsi are banded (Figure 38C). Proboscis has a broad, creamy band in the middle (Figure 38D). Broad, speckled scales are present in the wing (Figure 38E). Edwards (1941) reported that the distribution of this species in Africa is widespread: Gambia, Ghana, Nigeria, Congo, Uganda, Sudan, Ethiopia, Kenya, Tanganyika, Zanzibar, Mozambique, Malawi, Botswana, Central-North Africa, and Madagascar (GBIF Secretariat, 2023). The observation by Kent (2006) marks the first occurrence in Macha, Zambia. Distribution has also been noted in parts of Southeast Asia (Edwards 1941).

**FIGURE 38.**
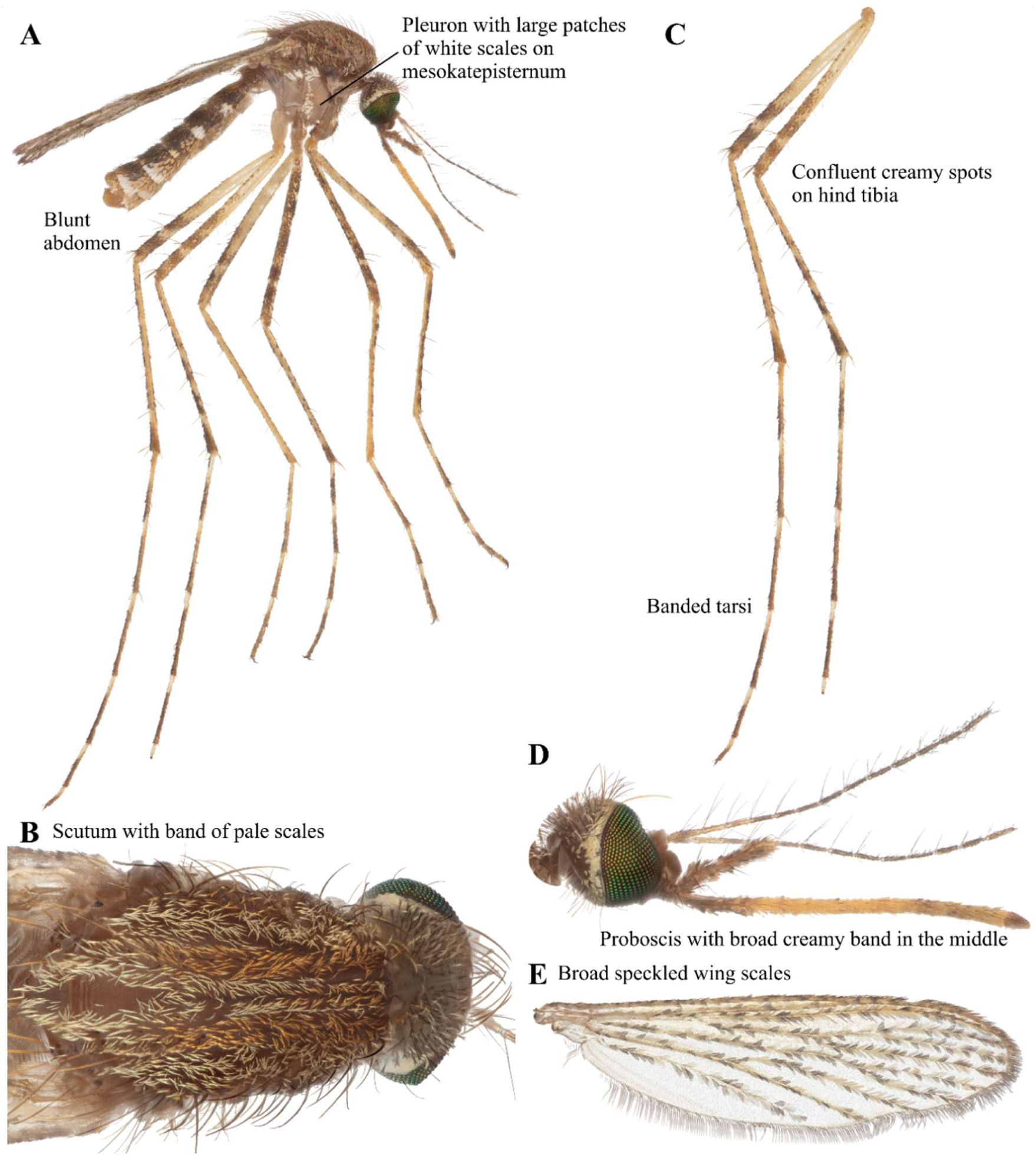
*Ma. uniformis* focus-stacked images. A: Lateral view. B: Scutum. C: Legs. D: Head. E: Wing.

### *Mimomyia (Etorleptiomyia*) *mediolineata* Theobald 1904 (Diptera: Culicidae)

*Mimomyia mediolineata* is a small mosquito with dark brown terga (Figure 39A). Its coxae have small spots of black scales. The scutum has yellow scales in the center and black scales along the sides (Figure 39B). Tibia has a mixture of dark and yellowish scales (Figure 39C). The proboscis is slightly swollen at the tip and has a broad yellow band in the middle (Figure 39D). Its palpi have white tips. The wing scales are sometimes heart-shaped (Figure 39E). Edwards (1941) reported the distribution of this species in Nigeria, the coast of Ghana, Uganda, Sudan, Malawi, Zanzibar, Congo, and parts of North-Central Africa. This species had also been identified in Botswana and Mozambique (GBIF Secretariat, 2023); however, our observation marks a new record of occurrence in Zambia.

**FIGURE 39.**
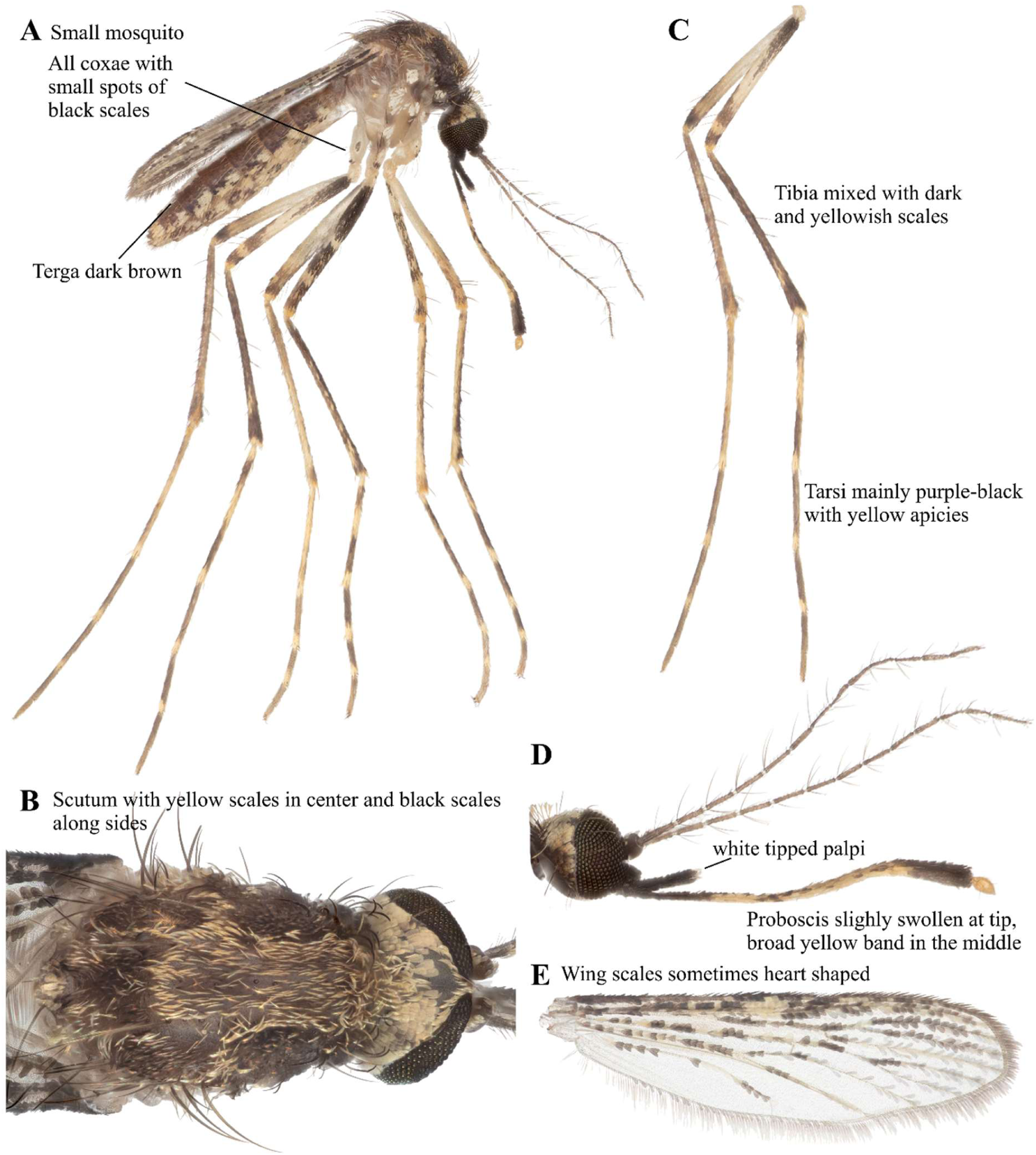
*Mi. mediolineata* focus-stacked images. A: Lateral view. B: Scutum. C: Legs. D: Head. E: Wing.

### *Mimomyia (Mimomyia) hispida* Theobald 1910 (Diptera: Culicidae)

*Mimomyia hispida* is a medium-sized mosquito with terga 2-6 having median and basolateral pale patches (Figure 40A). Scutum is dark brown in contrast with the yellow-scaled head (Figure 40B). Hind femora are largely dark, and hind tarsi are all dark (Figure 40C). The proboscis is swollen at the tip (Figure 40D). This species had been previously recorded in Sudan, Uganda, the coast of Ghana, Nigeria, Congo, and parts of southern Africa (Edwards 1941, GBIF Secretariat, 2023). Our observation marks a new record of occurrence in Macha, Zambia.

**FIGURE 40.**
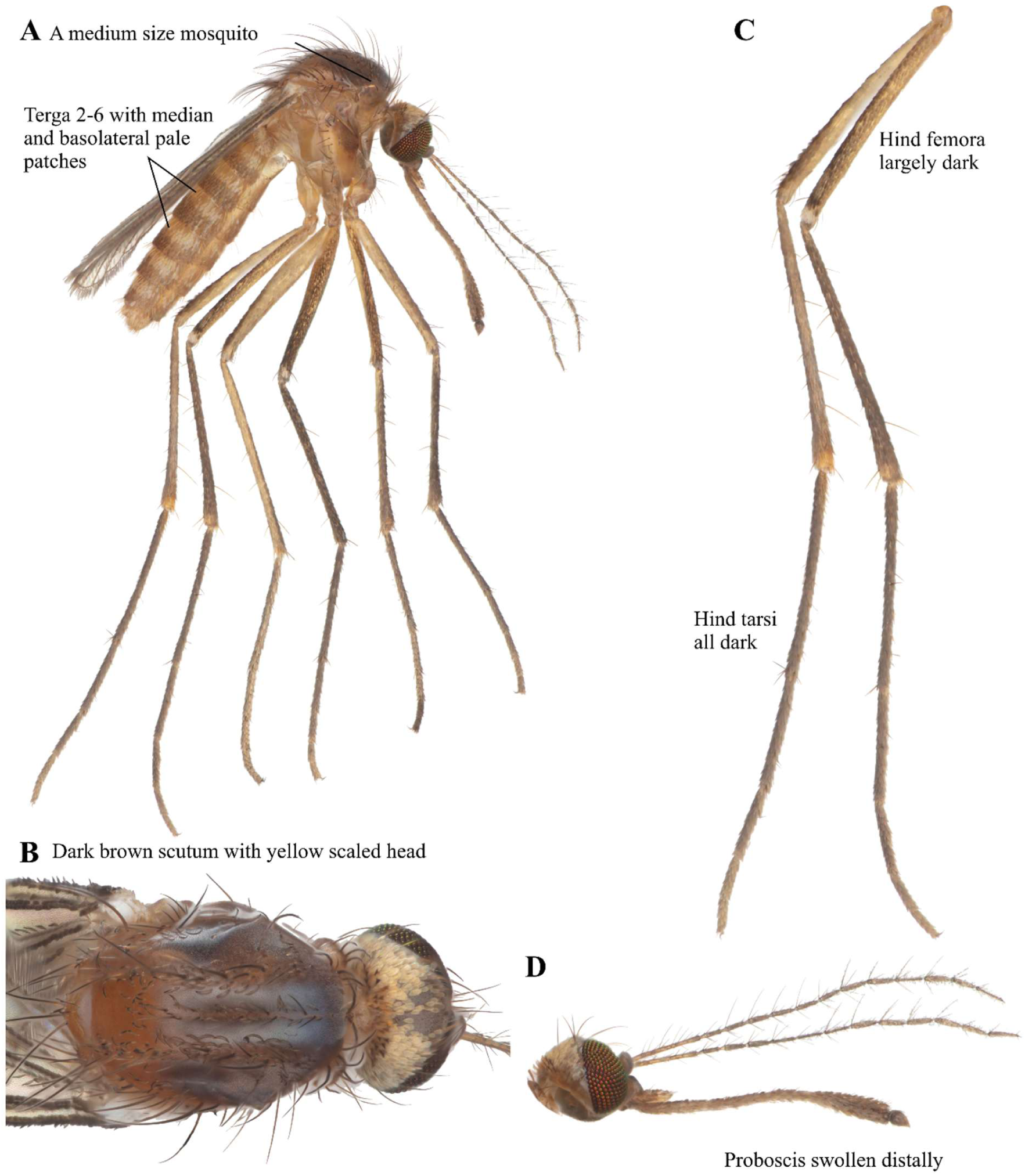
*Mi. hispida* focus-stacked images. A: Lateral view. B: Scutum. C: Legs. D: Head.

### *Toxorhynchites (Toxorhynchites*) *brevipalpis* Theobald 1901 (Diptera: Culicidae)

*Toxorhynchites brevipalpis* is a very large and colorful mosquito with predominantly blue color (Figure 41A). The proboscis is strongly curved, and this mosquito feeds only on plant nectar. The coxae have white scales. White tuft on tergite VI; black setal tufts on tergite VII; and orange tufts on tergite VIII (Figure 41A). No gold scales on thorax (Figure 41B). The legs are banded and mostly white on tarsomere 2 of all legs (Figure 41C). Its head has a metallic tint and its palpi and proboscis are blue (Figure 41D). The alula and upper calypter of its wing are mostly bare (Figure 41E). Edwards (1941) reported the occurrence of this species “confined to east and south Africa”. This species has been previously documented in Macha, Zambia (Kent 2006).

**FIGURE 41.**
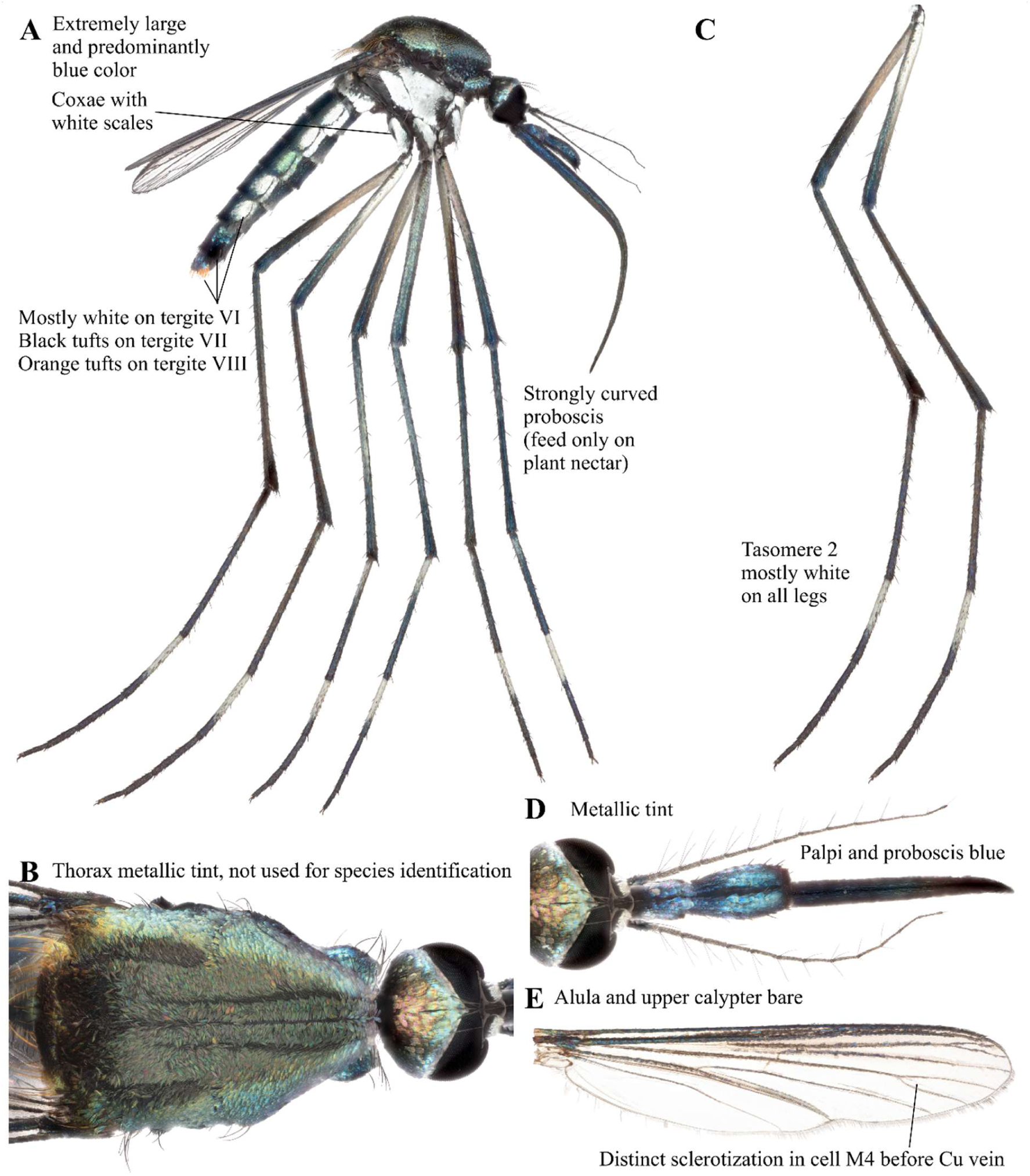
*Tx. brevipalpis* focus-stacked images. A: Lateral view. B: Scutum. C: Legs. D: Head. E: Wing.

### *Uranotaenia (Pseudoficalbia) mashonaensis* Theobald 1901 (Diptera: Culicidae)

*Uranotaenia mashonaensis* has a black spot in the thoracic integument above the wing-root (Figure 42A). Pleuron is yellowish. Scutum is brown (Figure 42B). Legs are brownish, but the underside of the femora is pale (Figure 42C). Flagellum and proboscis are dark (Figure 42D). Flat creamy scales are present around the eyes (Figure 42B and D). Edwards (1941) reported occurrence in the Eastern and western parts of Africa as well as in Congo. While this species has been observed in Rwanda and Zimbabwe (GBIF Secretariat, 2023), our observation marks the first record of occurrence in Zambia.

**FIGURE 42.**
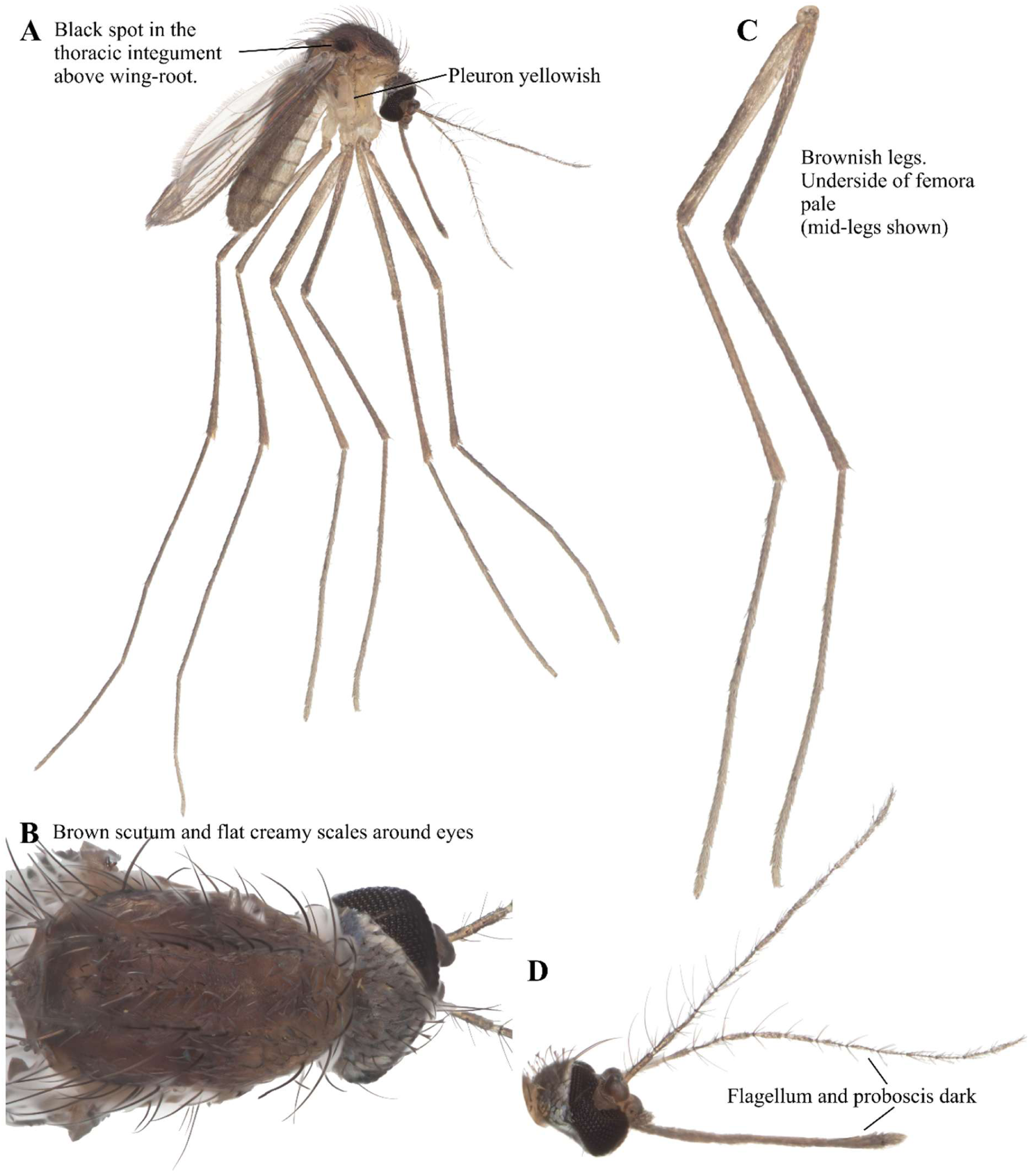
*Ur. mashoensis* focus-stacked images. A: Lateral view. B: Scutum. C: Legs. D: Head.

### *Uranotaenia (Uranotaenia) balfouri* Theobald 1904 (Diptera: Culicidae)

*Uranotaenia balfouri* is a small, dark mosquito with white scales on the margin of the scutum and pleuron (Figure 43A). The abdominal terga are dark (Figure 43A). There is white at the extreme base of R1 wing vein (Edwards 1941). The scutum is dark brown (Figure 43B). The legs are dark (Figure 43C). It has short palpi and dark proboscis (Figure 43D). This species has been previously documented in Macha, Zambia (Kent 2006) and western Zambia in 1944 (WRBU, 2018). Edwards (1941) reported observations in West and East Africa.

**FIGURE 43.**
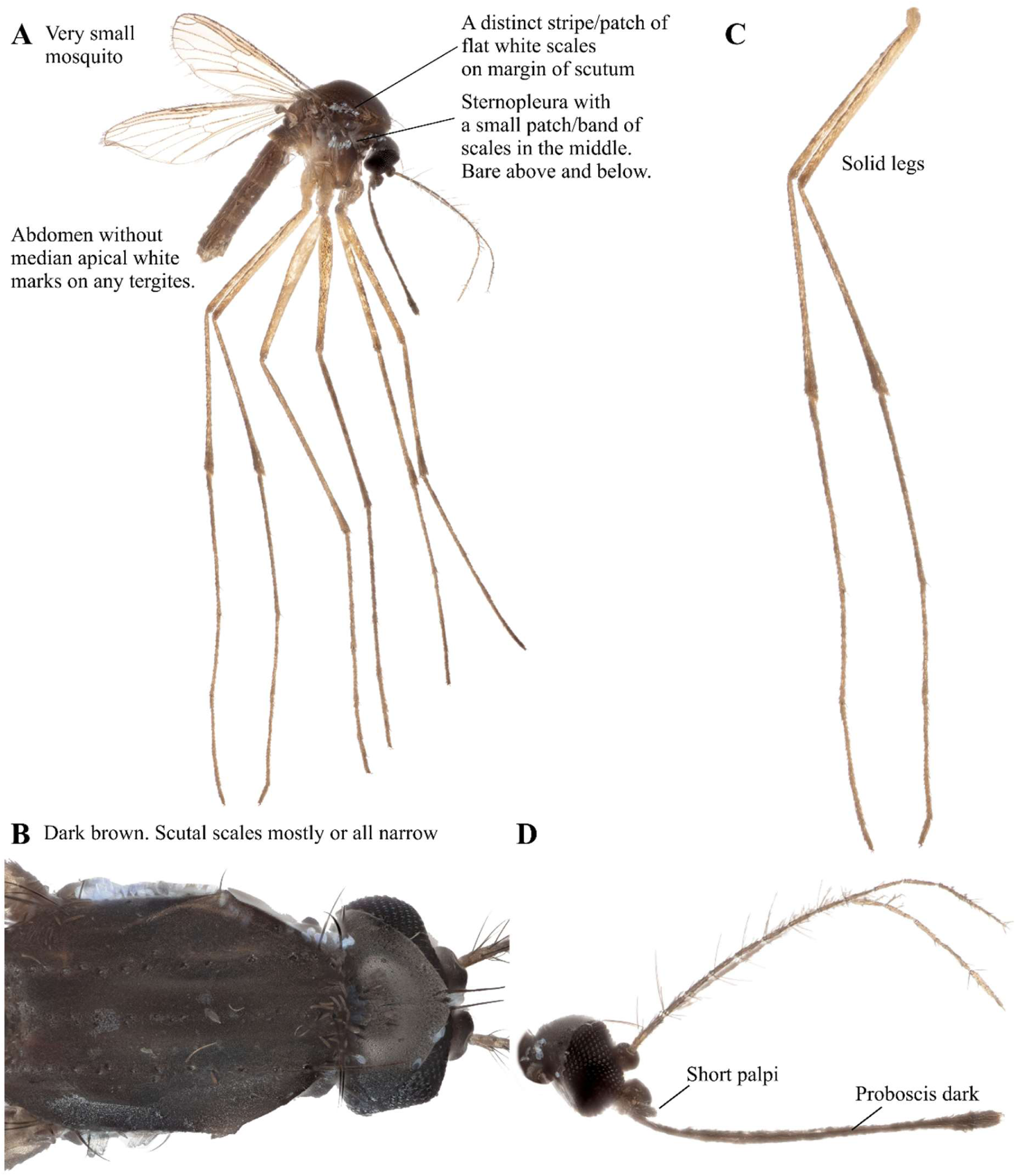
*Ur. balfouri* focus-stacked images. A: Lateral view. B: Scutum. C: Legs. D: Head.

### Phylogenetic relationships of mosquito species

Our data includes new COI sequences for five species which had no prior genetic information. These species include *Mimomyia hispida, Ficalibia uniforms, Aedes contiguus, Aedes embuensis,* and *Culex litwakae.* Other COI sequences had high sequence similarity (>98%) with corresponding conspecific COI sequences reported elsewhere (Figure 44). The exception was the Zambian *An. pharoensis* which has only 93.3-93.5% sequence similarity with other *An. pharoensis* found in Mali (MK586019) and Kenya (KU380430). *An. cf. pharoensis* from Zambia was similar (99.1%) to the same species found in Kenya (MW555793). The sequence similarity between Zambian *An. pharoensis* and *An. cf. pharoensis* was 95.2%. The COI sequences newly generated from this study is available from NCBI Nucleotide Database (PX317491-PX317524).

**FIGURE 44.**
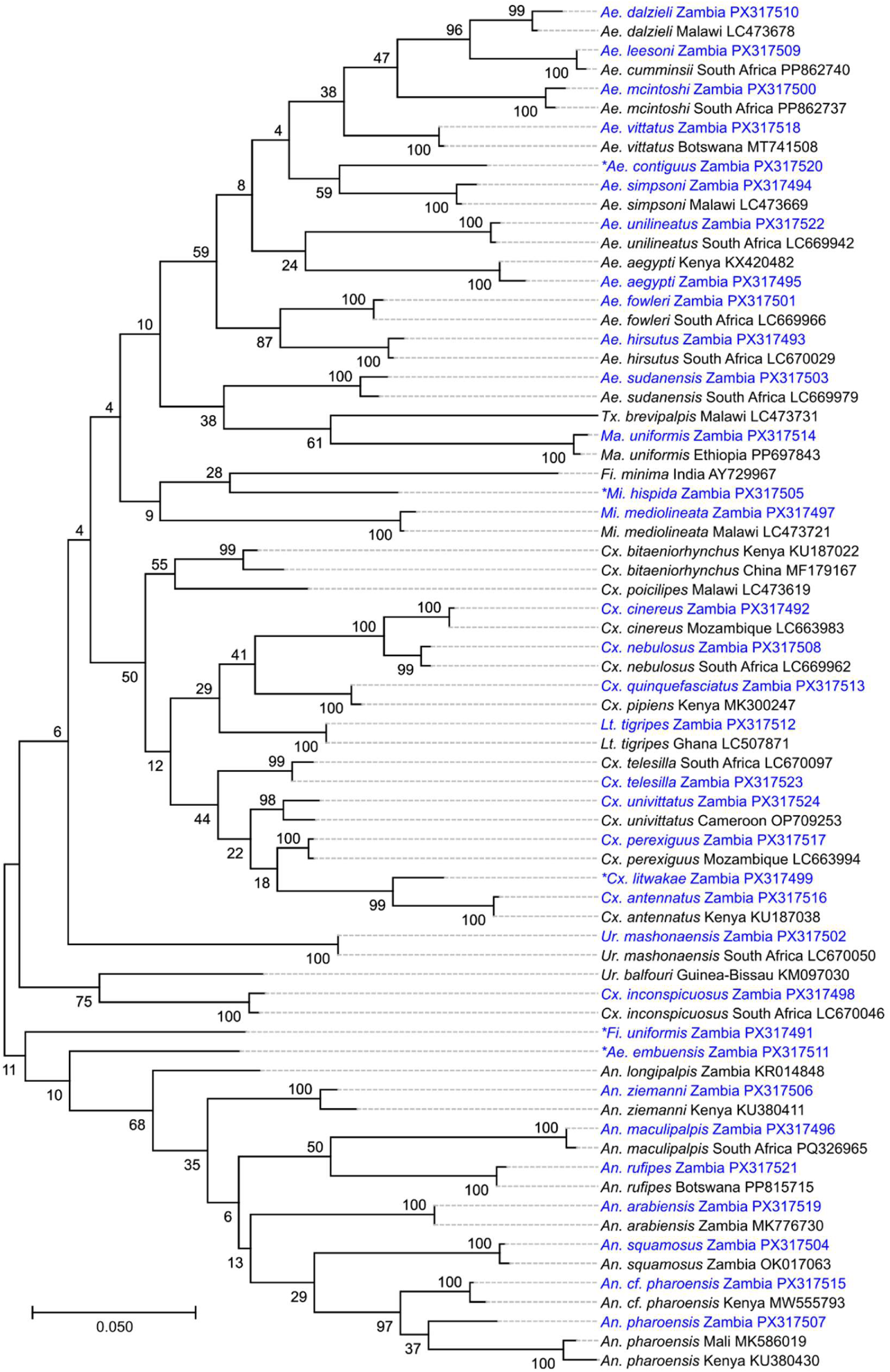
Maximum likelihood tree based on COI sequences of mosquitoes from southern Zambia with similar matches on GenBank. Samples from this study are marked in blue text. The numbers ranging from 4-100 at the branching point indicate the % agreement among 500 bootstraps. GenBank accession number and country of origin is shown after species name. Species with new COI records include *Mimomyia hispida, Ficalibia uniforms, Aedes contiguus, Aedes embuensis,* and *Culex litwakae* and marked with * in front of each species name.

The maximum likelihood tree grouped the mosquito species by genus in general (Figure 44). All *Anopheles* species fell under a single lineage. *Aedes* genus forms mostly one linage with the exception of *Ae. embuensis. Mansonia uniformis* and *Tx. brevipalpis* group were the closest kin to the *Aedes* group. Other genus like *Culex, Lutzia,* and *Uranotaenia* had a more complex relationship. *Culex inconspicuosus* was separated from the rest of the *Culex* species. *Uranotanenia balfouri* did not group together with *Ur. machonaensis. Lutzia tigripes* fell within the *Culex* group.

## Discussion

Here, we provide new species records of mosquitoes from Macha in southern Zambia as well as the updated list of species from that region. In total, 46 species have been reported to occur in southern Zambia, with 19 new species records reported in this paper since 2006 (Kent 2006). This is a 70% increase in mosquito species recorded in this region from the previous record (Kent 2006). We believe that this is a reflection of the poor investment in biodiversity research in Africa, rather than environmental changes enhancing biodiversity over two decades. Species richness reported in Lehmann et al. (2023) shows that Zambia’s biodiversity is as expected as other African countries given the size of the country. Our large increase in species records in Macha alone highlights that we may be grossly underestimating the richness of the mosquito species in Africa overall.

We provided high-resolution focus-stacked photographs for 36 of 46 species. Some of these species were only available as text or drawing descriptions in mosquito identification books and articles. This has added educational value when training new entomologists to recognize subtle morphological features important for discerning species. While we report mosquitoes from Macha and neighboring towns in southern Zambia (Sinazongwe and Livingston), many of the species listed here also occur in other eastern and southern African regions. Therefore, the high-resolution pictorial keys provide important resources for species identification training beyond the specific region of Zambia we sampled. By presenting the organism’s image, the records we generated may uncover diagnostic features that have not been used in dichotomous keys when more data of other species and variation within species are documented in the future.

Our data includes new COI sequences of five mosquito species which had no prior genetic information. The low similarity between Zambian *An. pharoensis* with other conspecific COI sequence elsewhere in Africa suggest that *An. pharoensis* group may have more than two members within the species group. While previous report suggested the likelihood of species complex within *An. squamosus,* based on COI sequences, we only found one species of *An. squamosus* in Zambia thus far. Although we cannot rule out the possibility of species differentiation based on nuclear markers like ITS2 as shown in *An. gambiae* complex (Hanemaaijer et al. 2018).

Understanding the biodiversity of mosquitoes provides a fuller picture of the ecosystem it occupies. While some blood-feeding insects are generalists in their blood feeding, some are specialists with a narrow range of animals they feed on. In some cases, finding blood-fed insects is easier than sighting or capturing the animal for biodiversity assessment (Mwakasungula et al. 2022). This process, called xenosurveillance, also gained attention for pathogen surveillance (Grubaugh et al. 2015, McMinn et al. 2023).

The accurate records of native species also have practical implications when monitoring invasive species and the biosecurity risks when a new vector or pathogen is introduced into the area. Monitoring species that cross continental boundaries like *An. stephensi* (Liu et al. 2024) could be a relatively straightforward process with appropriate training for corresponding species identification. The species movement within a continent, especially for understudied regions like Africa, can be dramatically undermined when the accurate records of native species are not present.

## Acknowledgement

We thank the entomology staff at Macha Research Trust who assisted with field collection.

This work was supported by grants from the United States National Institute of Health Small Grant Program (R03AI178041) and the United States Department of Agriculture (Cooperative Agreement 58-3022-3-028). Additional support was provided by the United States Department of Agriculture National Institute of Food and Agriculture Hatch (006108) and multistate Hatch project (7007941), the College of Agricultural and Life Sciences Dean’s Award to VTN at the University of Florida, and Global Fellows Award to YL at the University of Florida International Center.

